# Integrated bioinformatics analysis reveals novel key biomarkers and potential candidate small molecule drugs in gestational diabetes mellitus

**DOI:** 10.1101/2021.03.09.434569

**Authors:** Basavaraj Vastrad, Chanabasayya Vastrad, Anandkumar Tengli

## Abstract

Gestational diabetes mellitus (GDM) is one of the metabolic diseases during pregnancy. The identification of the central molecular mechanisms liable for the disease pathogenesis might lead to the advancement of new therapeutic options. The current investigation aimed to identify central differentially expressed genes (DEGs) in GDM. The transcription profiling by array data (E-MTAB-6418) was obtained from the ArrayExpress database. The DEGs between GDM samples and non GDM samples were analyzed with limma package. Gene ontology (GO) and REACTOME enrichment analysis were performed using ToppGene. Then we constructed the protein-protein interaction (PPI) network of DEGs by the Search Tool for the Retrieval of Interacting Genes database (STRING) and module analysis was performed. Subsequently, we constructed the miRNA-hub gene network and TF-hub gene regulatory network by the miRNet database and NetworkAnalyst database. The validation of hub genes was performed through receiver operating characteristic curve (ROC). Finally, the candidate small molecules as potential drugs to treat GDM were predicted by using molecular docking. Through transcription profiling by array data, a total of 869 DEGs were detected including 439 up regulated and 430 down regulated genes. Biological process analysis of GO enrichment analysis showed these DEGs were mainly enriched in reproduction, nuclear outer membrane-endoplasmic reticulum membrane network, identical protein binding, cell adhesion, supramolecular complex and signaling receptor binding. Signaling pathway enrichment analysis indicated that these DEGs played a vital in cell surface interactions at the vascular wall and extracellular matrix organization. Ten genes, HSP90AA1, EGFR, RPS13, RBX1, PAK1, FYN, ABL1, SMAD3, STAT3, and PRKCA in the center of the PPI network, modules, miRNA-hub gene regulatory network and TF-hub gene regulatory network were associated with GDM, according to ROC analysis. Finally, the most significant small molecules were predicted based on molecular docking. Our results indicated that HSP90AA1, EGFR, RPS13, RBX1, PAK1, FYN, ABL1, SMAD3, STAT3, and PRKCA could be the potential novel biomarkers for GDM diagnosis, prognosis and the promising therapeutic targets. The current might be essential to understanding the molecular mechanism of GDM initiation and development.

## Introduction

Gestational diabetes mellitus (GDM) is the diabetes diagnosed during pregnancy, which affecting 2–5% of all pregnant women worldwide [1–2]. Risk factors associated with GDM includes obesity, previous occurrence of diabetes, family history of type 2 diabetes, preeclampsia, hypertension, cardiovascular diseases and genetic factors [3]. In GDM blood glucose levels are elevated during the third trimester of pregnancy [4]. Moreover, the elevated glucose level in pregnancy is closely linked with detrimental consequences in the newborn babe, such as fetal hyperglycemia and cardiovascular disease [5]. Therefore, investigating the molecular mechanisms of GDM and early screening of patients with GDM are essential to restrain the occurrence and progression of GDM.

It is therefore essential to find new genes and pathways that are associated with GDM and patient prognosis, which might not only help to explicate the underlying molecular mechanisms associated, but also to discover new diagnostic molecular markers and therapeutic targets. Transcription profiling by array can rapidly detect gene expression on a global basis and are particularly useful in screening for differentially expressed genes (DEGs) [6]. Gene chips allow the analysis of gene expression in a high throughput way with great sensitivity, specificity and repeatability. A symbolic amount of data has been produced via the use of gene chips and the majority of such gene expression data has been uploaded and stored in public databases. Previous investigation concerning GDM transcription profiling by array have found hundreds of DEGs [7–8]. The availability of bioinformatics analysis based on high-throughput technology enabled the investigation of the modification in gene expression and the interaction between differential genes in GDM, to provide novel insights for further in-depth The availability of bioinformatics analysis based on high-throughput technology enabled the investigation of the alterations in mRNA expression and the interaction between differential genes in GDM, to provide novel insights for further in depth GDM investigations.

In the current investigation, public transcription profiling by array data of E-MTAB-6418 from ArrayExpress database was downloaded. A total of 38 GDM samples and 70 non GDM samples data in E-MTAB-6418 were available. DEGs between GDM and non GDM were filtered and obtained using bioconductor package limma in R software. Gene Ontology (GO) and REACTOME pathway enrichment analyses of the DEGs were performed. The functions of the DEGs were further assessed by PPI network and modular analyses to identify the hub genes in GDM. Subsequently, miRNA-hub gene regulatory network and TF-hub gene regulatory network were constructed to identify the target genes, miRNAs and TFs in GDM. Hub genes were validated by receiver operating characteristic curve (ROC). Finally, screening of small drug molecules carried out by using molecular docking. The investigation was designed to obtain deep insights during the pathogenesis of GDM.

## Materials and methods

### Transcription profiling by array data information

The mRNA expression profile E-MTAB-6418 [9] based on A-MEXP-2072 - Illumina HumanHT-12_V4_0_R2_15002873_B was downloaded from the ArrayExpress database (https://www.ebi.ac.uk/arrayexpress/) [10], which included 38 GDM samples and 70 non GDM samples.

### Identification of DEGs

To obtain differentially expressed genes (DEGs) between GDM samples and non GDM samples. After limma package in R analysis [11], results including adjusted P values (adj. P. Val) and log FC were provided. Cut-off criterion was set as adj. P. Val <0.05, |log FC| > 1.158 for up regulated genes and |log FC| < -0.83 for down regulated genes. A list of candidate DEGs was obtained via the above methods.

### Gene ontology and pathway enrichment of DEGs analysis

Gene ontology (GO) analysis (http://geneontology.org/) [12] and REACTOME (https://reactome.org/) [13] pathway enrichment analysis are both integrated in the ToppGene (ToppFun) (https://toppgene.cchmc.org/enrichment.jsp) [14] program. Therefore, ToppGene is capable of providing comprehensive annotations for functional and pathway interpretations. In this experiment, DEGs were uploaded onto ToppGene in order to perform related GO and REACTOME pathway enrichment analyses. The cut-off criterion was set as P<0.05.

### PPI network establishment and modules selection

Search Tool for the Retrieval of Interacting Genes StringDB interactome (https://string-db.org/) is a database of known and predicted protein-protein interactions [15]. All candidate DEGs were posted into the STRING website, with a confidence score of ≥ set as the cut-off criterion for PPI network construction.

Then, Cytoscape (version 3.8.2, http://www.cytoscape.org/) [16] software was utilized to construct protein interaction relationship network. The Network Analyzer plugin was performed to scale node degree [17], betweenness centrality [18], stress centrality [19] and closeness centrality [20] of the PPI network. Significant modules in the visible PPI network were screened using the PEWCC1 (http://apps.cytoscape.org/apps/PEWCC1) [21] plugin. Degree cut-off=2, node score cut-off=0.2, k-core=2, and max depth=100 were set as the cut-off criterion. Three highest-degree modules were extracted, and the potential mechanisms of each module were investigated with ToppGene. A degree of ≥ was set as the filter criterion. Hub genes with high degree were selected as the potential key genes and biomarkers.

### miRNA-hub gene regulatory network construction

The miRNet database (https://www.mirnet.ca/) [22] is an open-source platform mainly focusing on miRNA-target interactions. miRNet utilizes fourteen established miRNA-target prediction databases, including TarBase, miRTarBase, miRecords, miRanda, miR2Disease, HMDD, PhenomiR, SM2miR, PharmacomiR, EpimiR, starBase, TransmiR, ADmiRE, and TAM 2.0. In this study, miRNAs were considered the targeted miRNAs of hub genes. Subsequently, the network of the hub genes and their targeted miRNAs was visualized by Cytoscape software.

### TF-hub gene regulatory network construction

The NetworkAnalyst database (https://www.networkanalyst.ca/) [23] is an open-source platform mainly focusing on TF-target interactions. NetworkAnalyst utilizes three established TF-target prediction databases, including ENCODE, JASPAR, ChEA. In this study, TFs were considered the targeted TFs of hub genes based on ChEA database. In this study, TFs were considered the targeted TFs of hub genes. Subsequently, the network of the hub genes and their targeted TFs was visualized by Cytoscape software.

### Receiver operating characteristic (ROC) curve analysis

The receiver operating characteristic curve (ROC) was constructed by predicting the probability of a diagnosis being of high or low integrated score of significant hub gene expression in GDM. Area under curve (AUC) analysis was operated to calculate the diagnostic ability by using the statistical package pROC in R software [24].

### RT-PCR Analysis

The HTR-8/SVneo (ATCC CRL3271) cell line procured from ATCC. For normal HTR-8/SVneo (ATCC CRL3271) cell line was grown in RPMI-1640 medium added with 10% fetal bovine serum, containing 5.5 mM glucose, and 1% penicillin/streptomycin. Incubate this cell line at 37°C in a 5% CO2 in humidified cell culture incubator. Similarly, for GDM HTR-8/SVneo (ATCC CRL3271) cell line was grown in RPMI-1640 medium added with 10% fetal bovine serum, containing 5.5 mM glucose, and 1% penicillin/streptomycin. Incubate this cell line at 37°C in a 5% CO2 in humidified cell culture incubator for 24 hrs, then stimulated with various concentrations 40 mM of D-glucose for 6 h. TRIzol (cat. no. 9109; Takara Bio, Inc.) was used to isolate total RNA from HTR-8/SVneo cell line and HTR-8/SVneo cell line treated with glucose according to the manufacturer’s instructions. TRI Reagent (Sigma, USA).was used to isolate total RNA from each tissue sample according to the manufacturer’s instructions. Then, total RNA was reverse transcribed into cDNAs using the FastQuant RT kit (with gDNase; Tiangen Biotech Co., Ltd.). RT-qPCR was performed to measure the levels of cDNAs using a QuantStudio 7 Flex real-time PCR system (Thermo Fisher Scientific, Waltham, MA, USA). RT-PCR procedure was performed as follows: Pre-denaturation at 95°C for 30 sec for 1 cycle followed by 40 cycles of 95°C for 5 sec and 60°C for 20 sec. The relative expression level of the hub genes was calculated following comparative CT method [25]. β actin was used to normalize the mRNA expression level. The primer sequences are listed in Table 1.

**Table 1.**
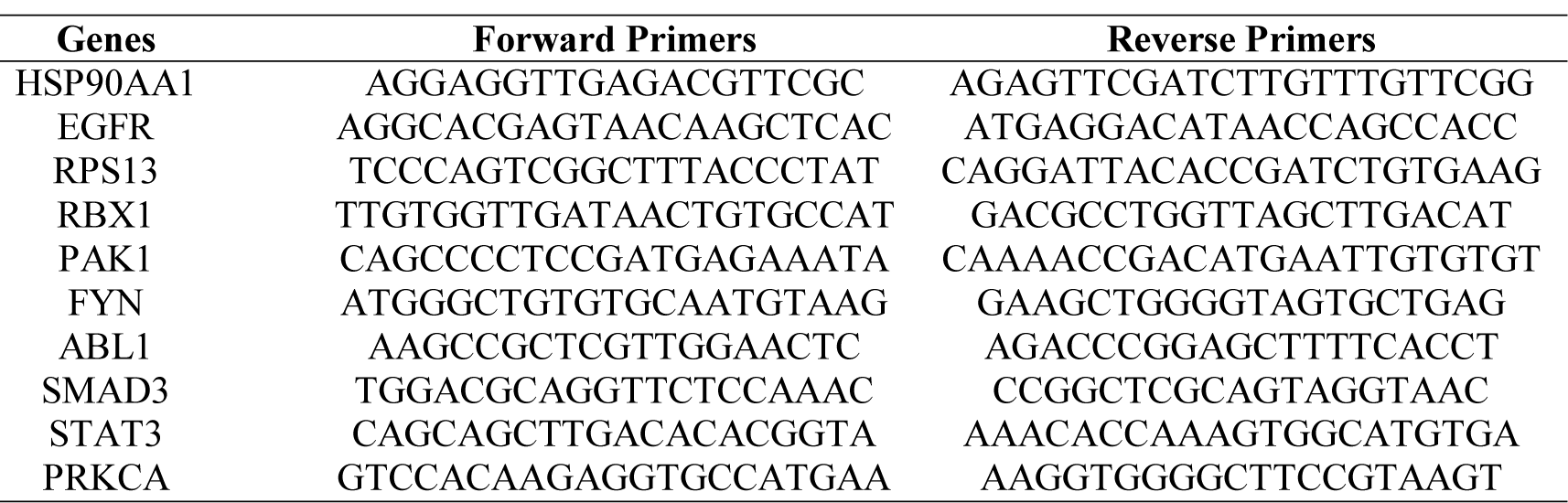
The sequences of primers for quantitative RT-PCR

### Molecular docking experiments

Molecular docking was used to find biologically active hits among the designed ligands. Using perpetual software module BIOVIA Discovery Studio (Perpetual), Surflex-Docking docking studies were conducted on active constituents. The lowest binding energy conformation was presumed to form a stable complex within the active site of the over expressed proteins. The 2D structures were sketched using Chemdraw software, imported and saved into sdf. format using Open Babelfree software. The protein structure was processed after introduction of the protein, the co-crystallized ligand and all the water molecule were excluded from the crystal structure; morehydrogen was added and refined the side chain. This study employed CDOCKER, a grid-based molecular docking approach that utilizes the CHARMm force field. A higher number indicates a stronger bond. The CDOCKER score is expressed as a negative number (-CDOCKER ENERGY).The H-bonds, van der Waals, and electrostatic interactions between the target protein and the ligand were used to measure the CDOCKER energy. The modeled protein’s binding site was determined using the template protein’s crystal data and proteins in which do not Co-crystallized ligand generated binding site automatically. To make it easier for ligands to interact with amino acids, the binding site sphere center was set at 9 Å radius. Furthermore, using smart minimizer algorithm, CHARMm force field was applied followed by energy minimization to define local minima (lowest energy conformation) of the modeled over expressed proteins with an energy gradient of 0.1 kcal mol^−1^ Å^−1^ respectively. The energy minimized receptor protein and the set of 44 natural molecules which was reported as effective in diabetes mellitus and the well-known commonly used allopathic drug Metformin and Glyburide were used as standard and to compare the binding interactions with natural molecules on over expressed proteins in gestational diabetes. The binding site sphere radius set at *X* = 29.50, *Y* = −31.38 and *Z* = −38.79 were submitted to the CDOCKER parameter and also calculated binding energy. The X-ray co-crystallized structure and were extracted from Protein Data Bank of PDB code of 4UV7, 5NJX, 3Q4Z and 3FNI of over expressed genes of Epidermal growth factor receptor (EGFR), Heat shock protein 90 alpha family class A member 1 (HSP90AA1), P21 RAC1 activated kinase 1 (PAK1) and Ring-box 1 (RBX1) respectively in gestational diabetes were selected for docking studies [26–29]. The best position was inserted into the molecular area between the protein and the ligand. The 2D and 3D interaction of amino acid molecules was achieved using the free online Discovery Studio Visualizer.

## Results

### Identification of DEGs

Transcription profiling by array data sets was obtained from the ArrayExpress database containing GDM samples and non GDM samples; E-MTAB-6418. Then, the R package named “limma” was processed for analysis with adjusted P < 0.05, |log FC| > 1.158 for up regulated genes and |log FC| < -0.83 for down regulated genes. All DEGs were displayed in volcano maps (Fig. 1). A total of 869 genes were finally obtained including 439 up regulated genes and 430 down regulated genes in the GDM samples compared to the non GDM samples and are listed Table 1. Top 869 genes in this dataset were displayed in the heatmap (Fig. 2).

**Fig. 1.**
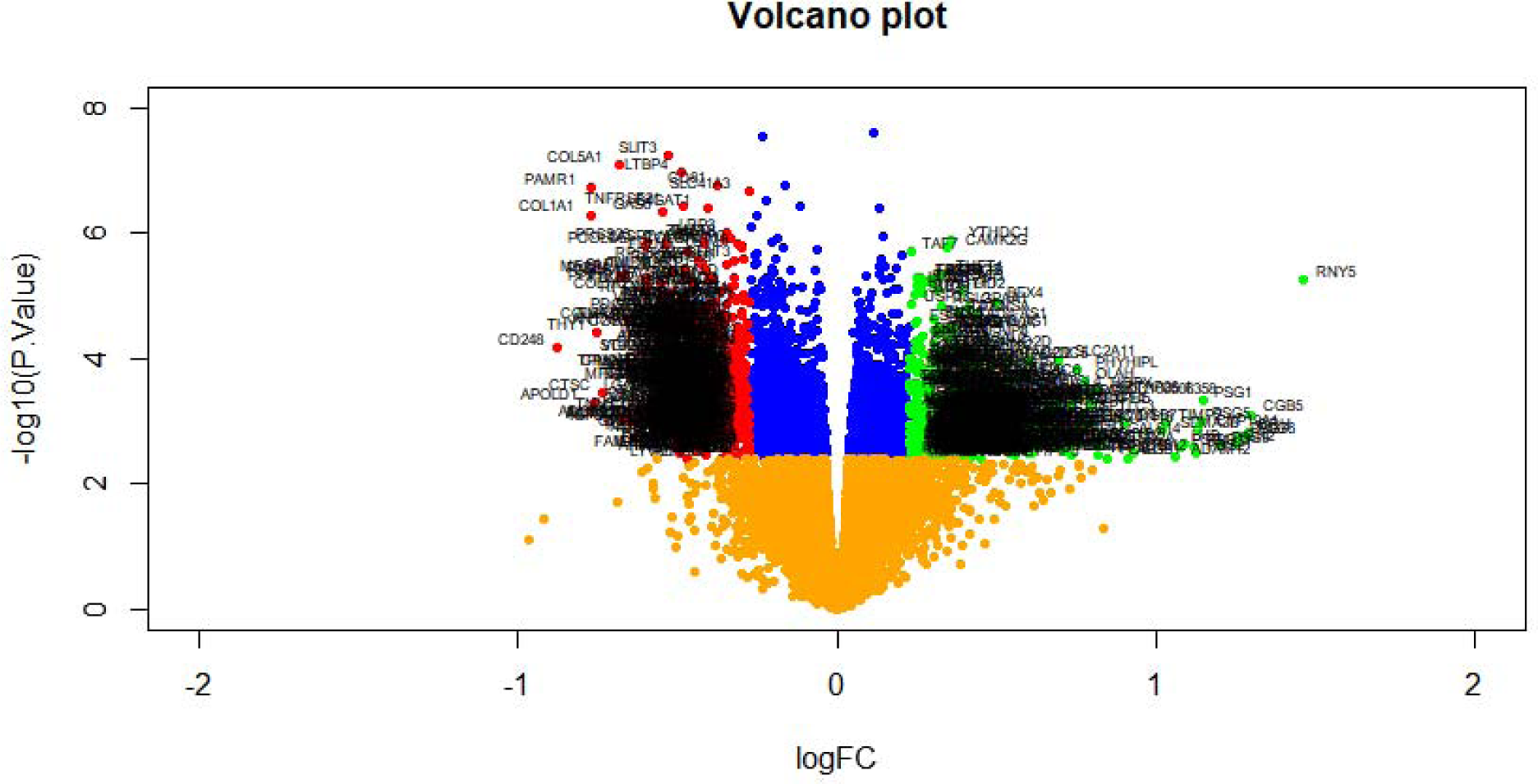
Volcano plot of differentially expressed genes. Genes with a significant change of more than two-fold were selected. Green dot represented up regulated significant genes and red dot represented down regulated significant genes.

**Fig. 2.**
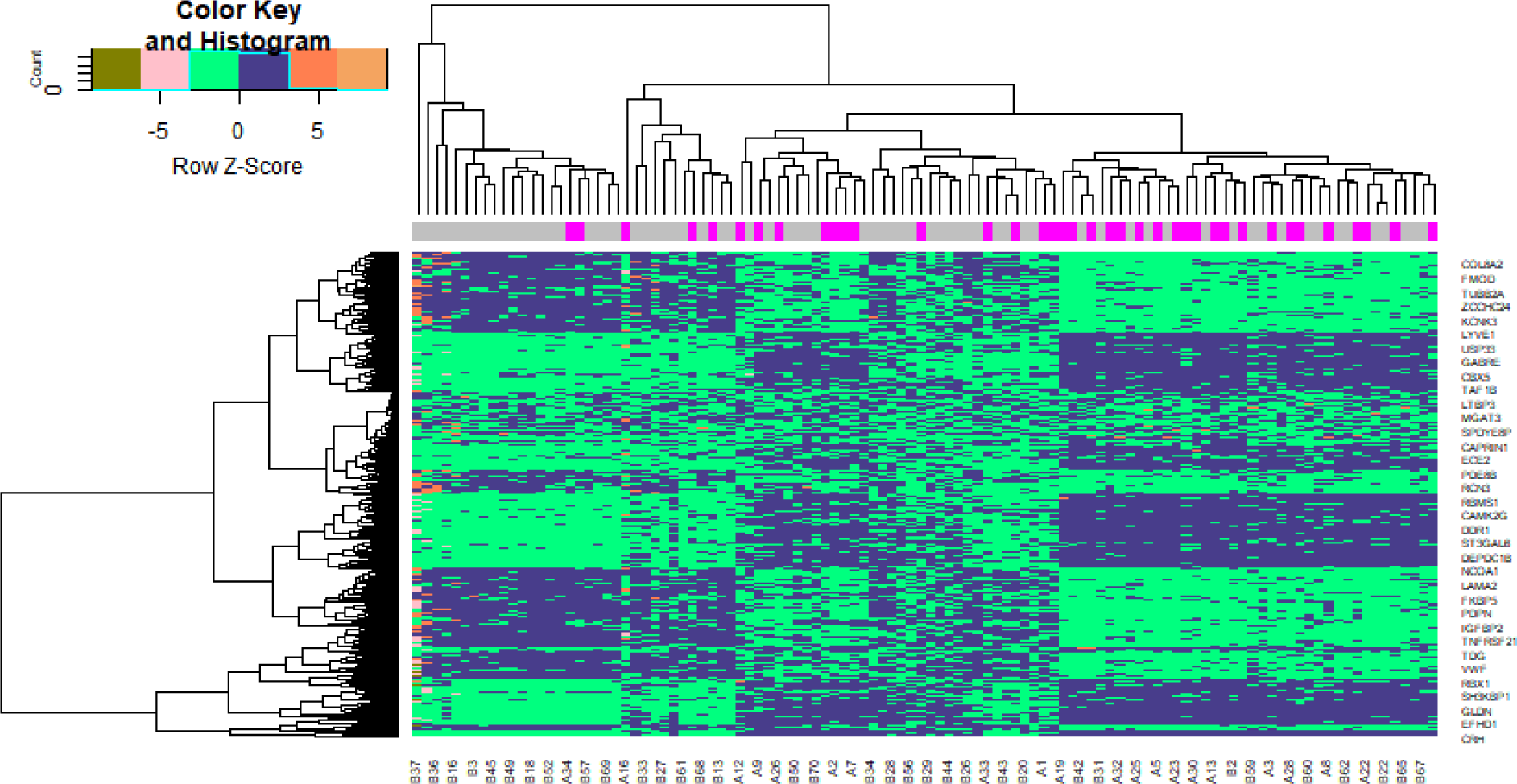
Heat map of differentially expressed genes. Legend on the top left indicate log fold change of genes. (A1 – A38= Gestational diabetes mellitus; B1 – B70 = Gestational diabetes mellitus)

### Gene ontology and pathway enrichment of DEGs analysis

To clarify the major functions of these DEGs, we first explored the associated biological processes and REACTOME pathways. The top highly enriched GO terms were divided into three categories: biological process (BP), cellular component (CC), and molecular function (MF) and are listed in Table 2. The most enriched GO terms in BP was reproduction, macromolecule catabolic process, cell adhesion and localization of cell, that in CC was nuclear outer membrane-endoplasmic reticulum membrane network, golgi apparatus, supramolecular complex and cell junction, and that in MF were identical protein binding, molecular function regulator, signaling receptor binding and molecular function regulator. In the REACTOME pathway enrichment analysis, the DEGs were mostly enriched in cell surface interactions at the vascular wall, epigenetic regulation of gene expression, extracellular matrix organization and axon guidance and are listed in Table 3.

**Table 2.**
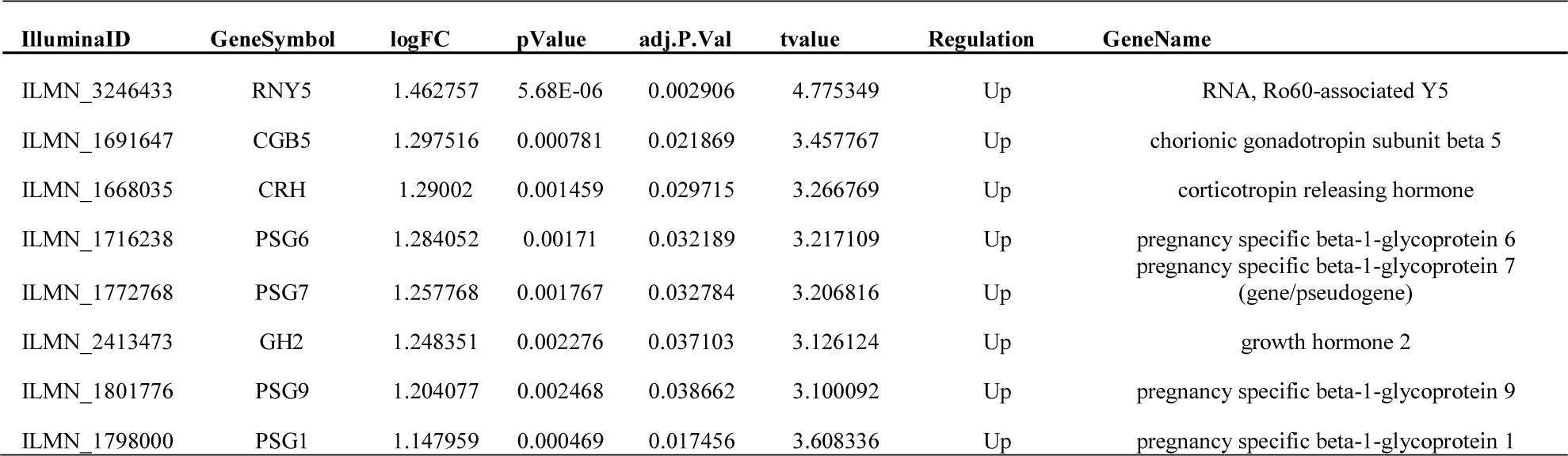
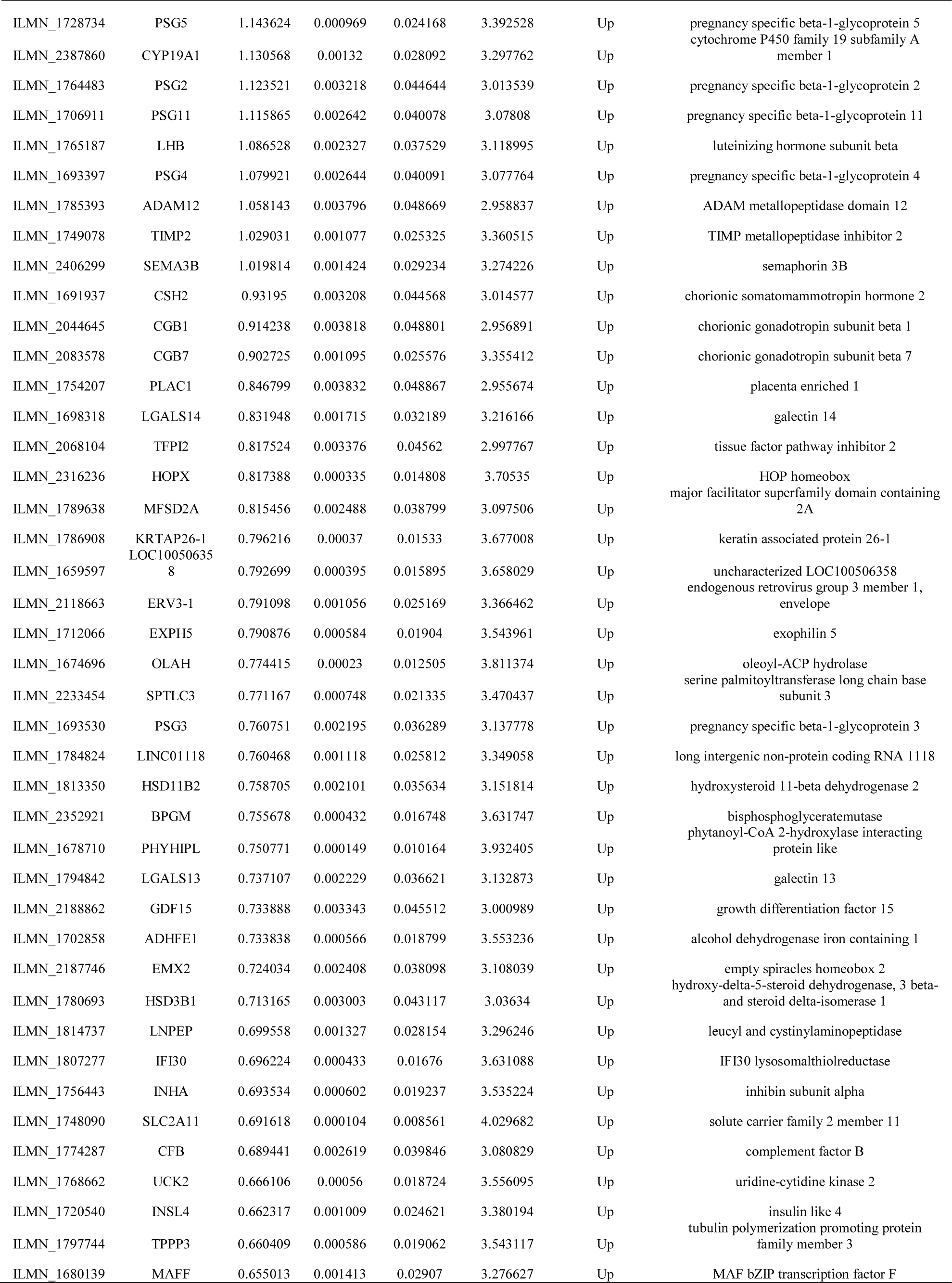
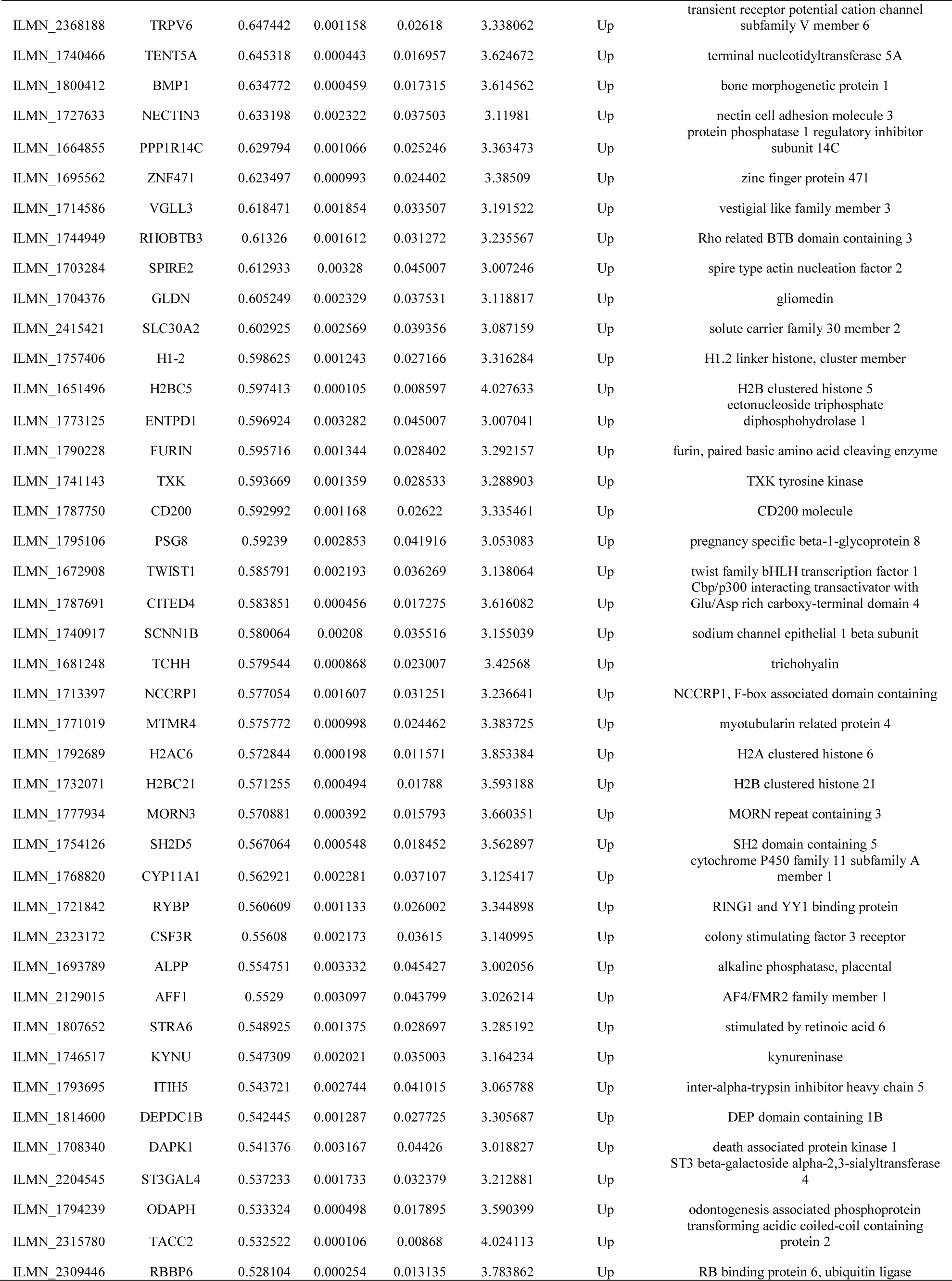
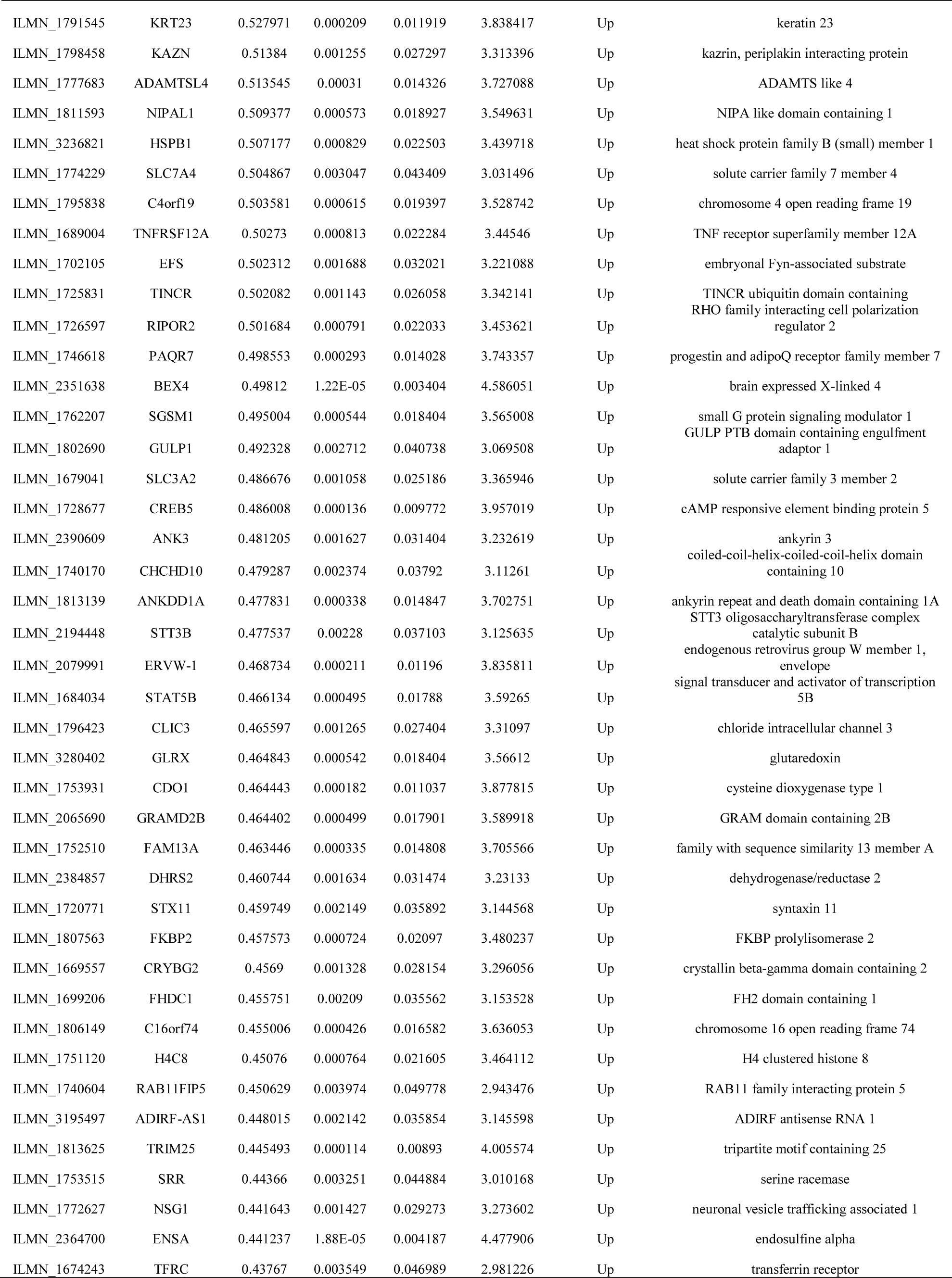
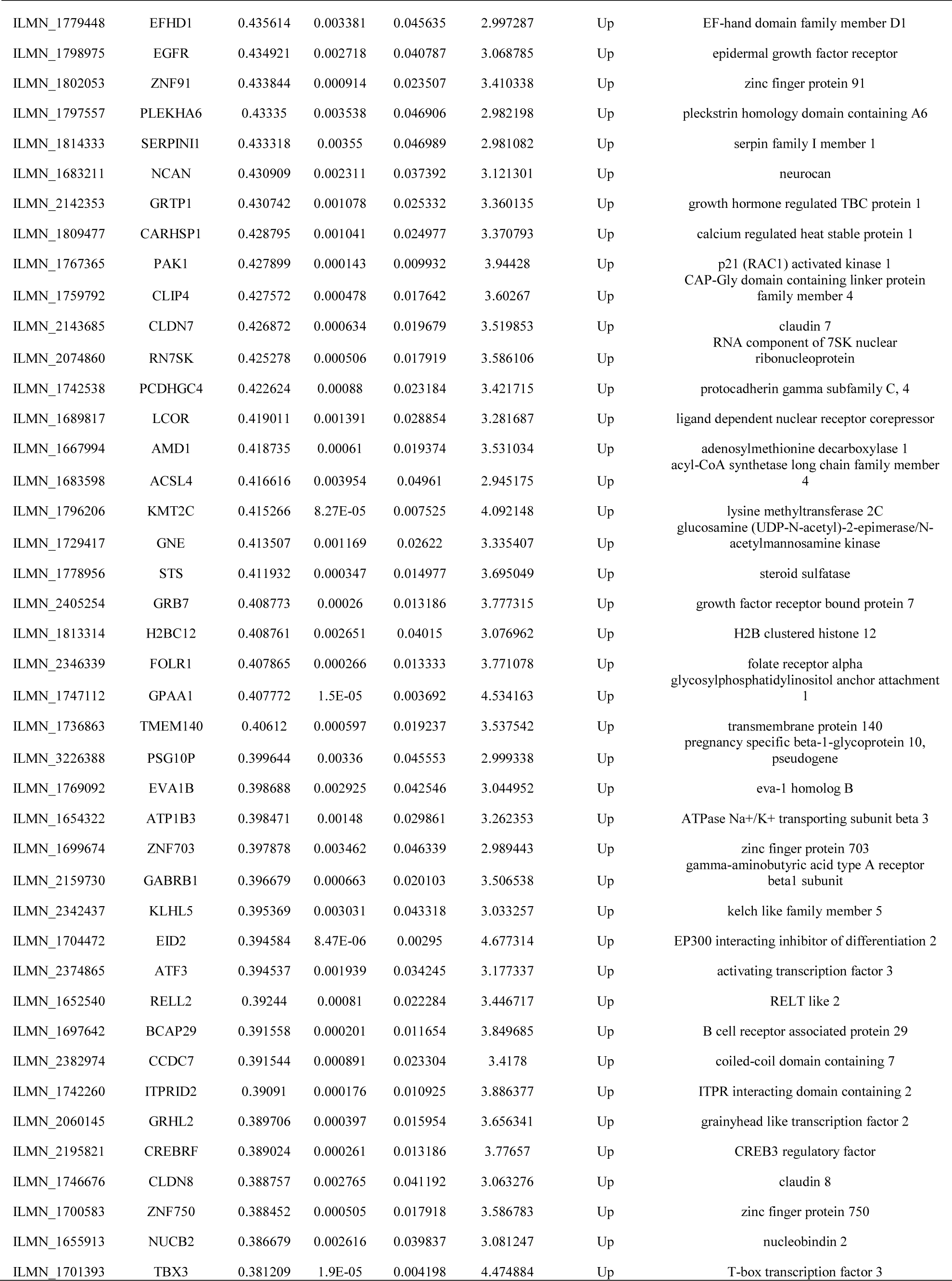
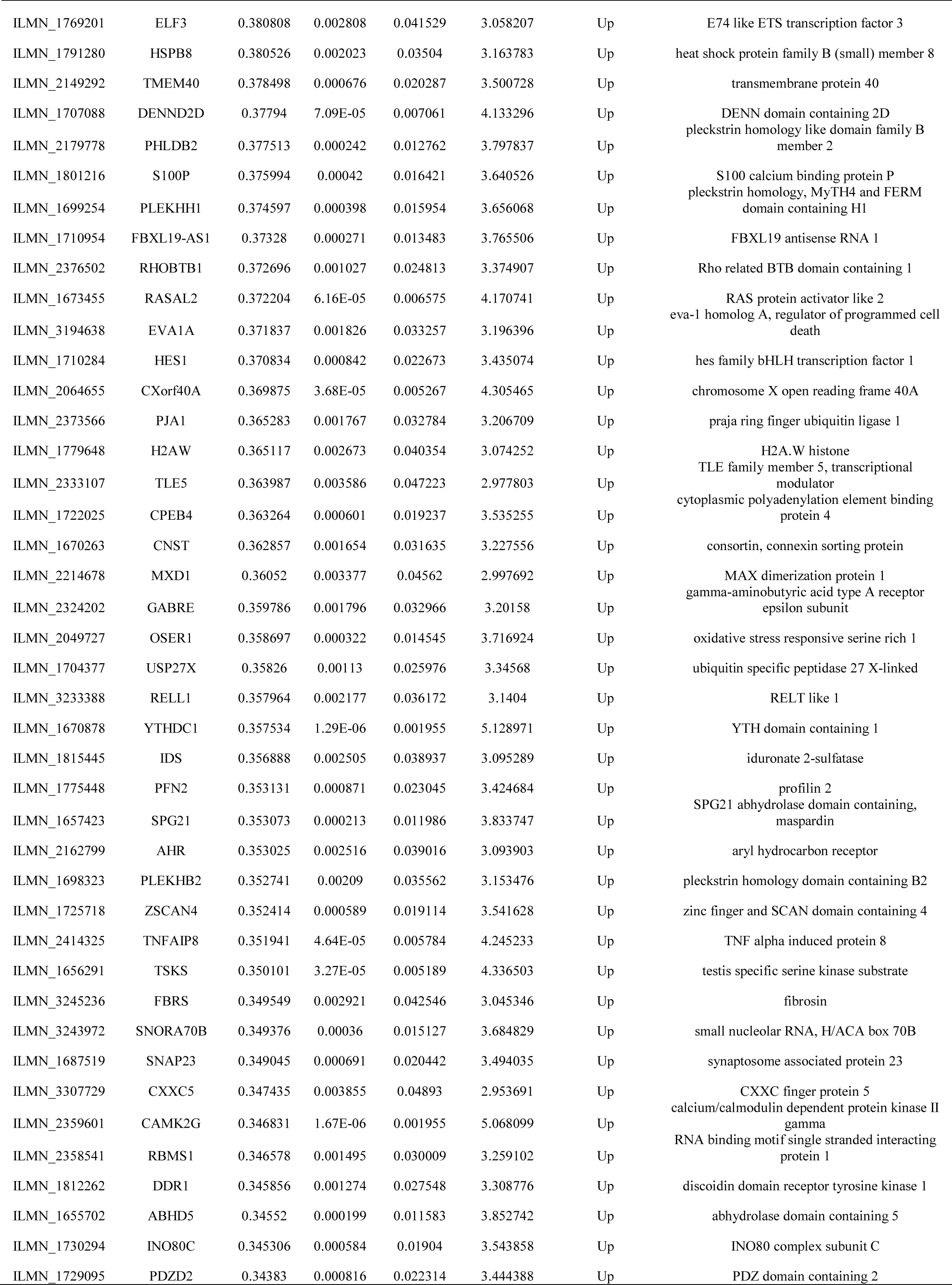
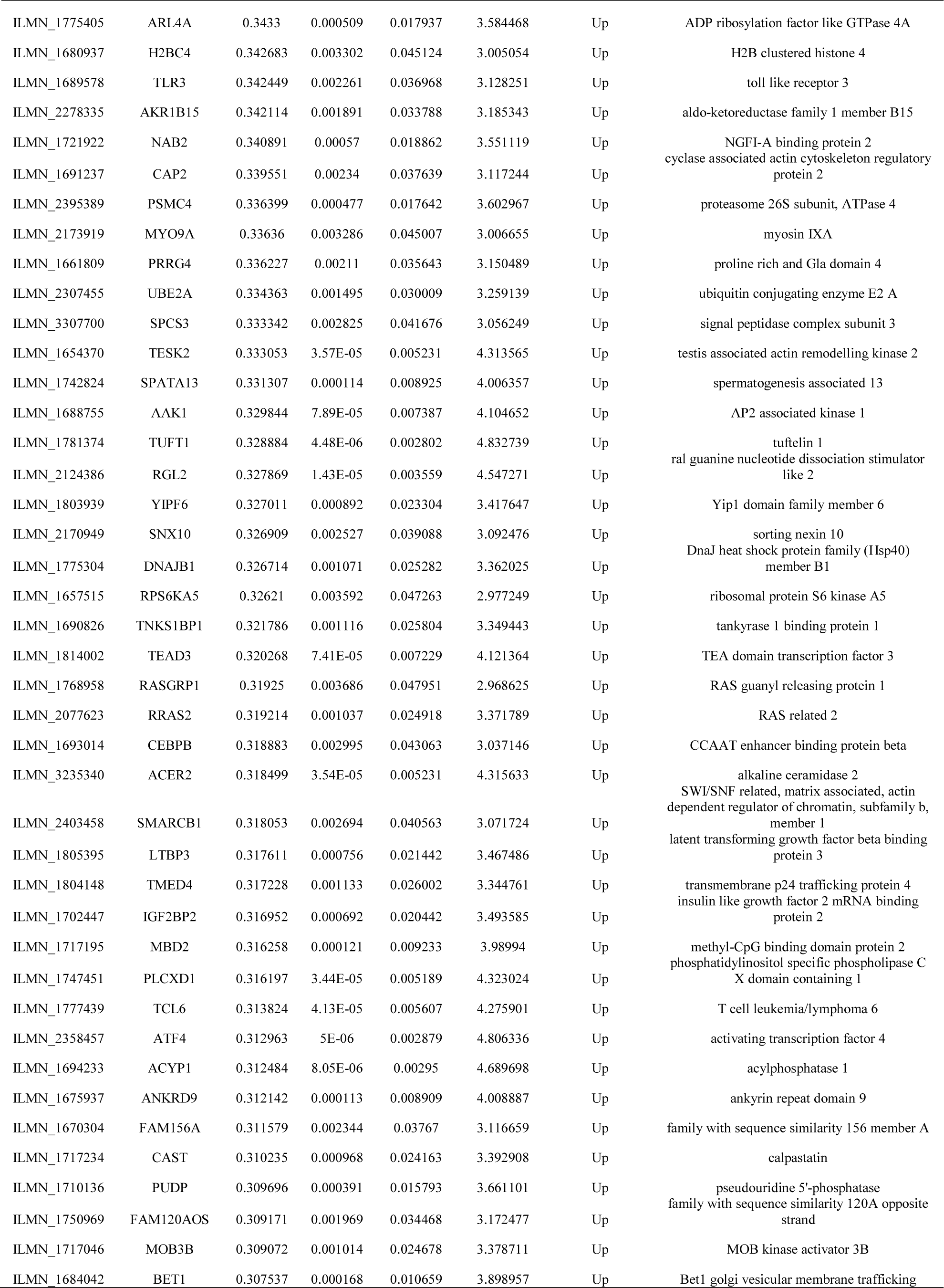
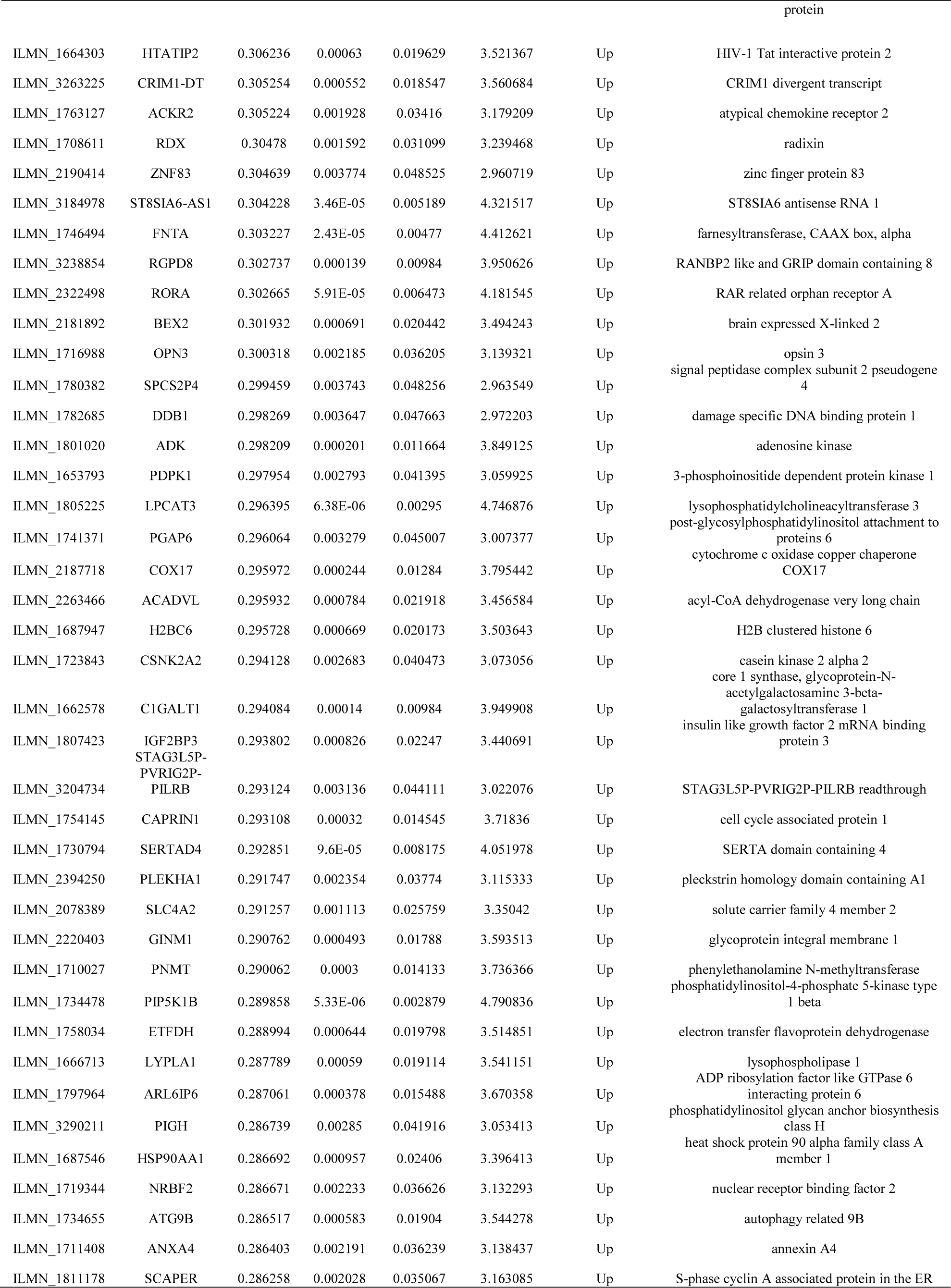
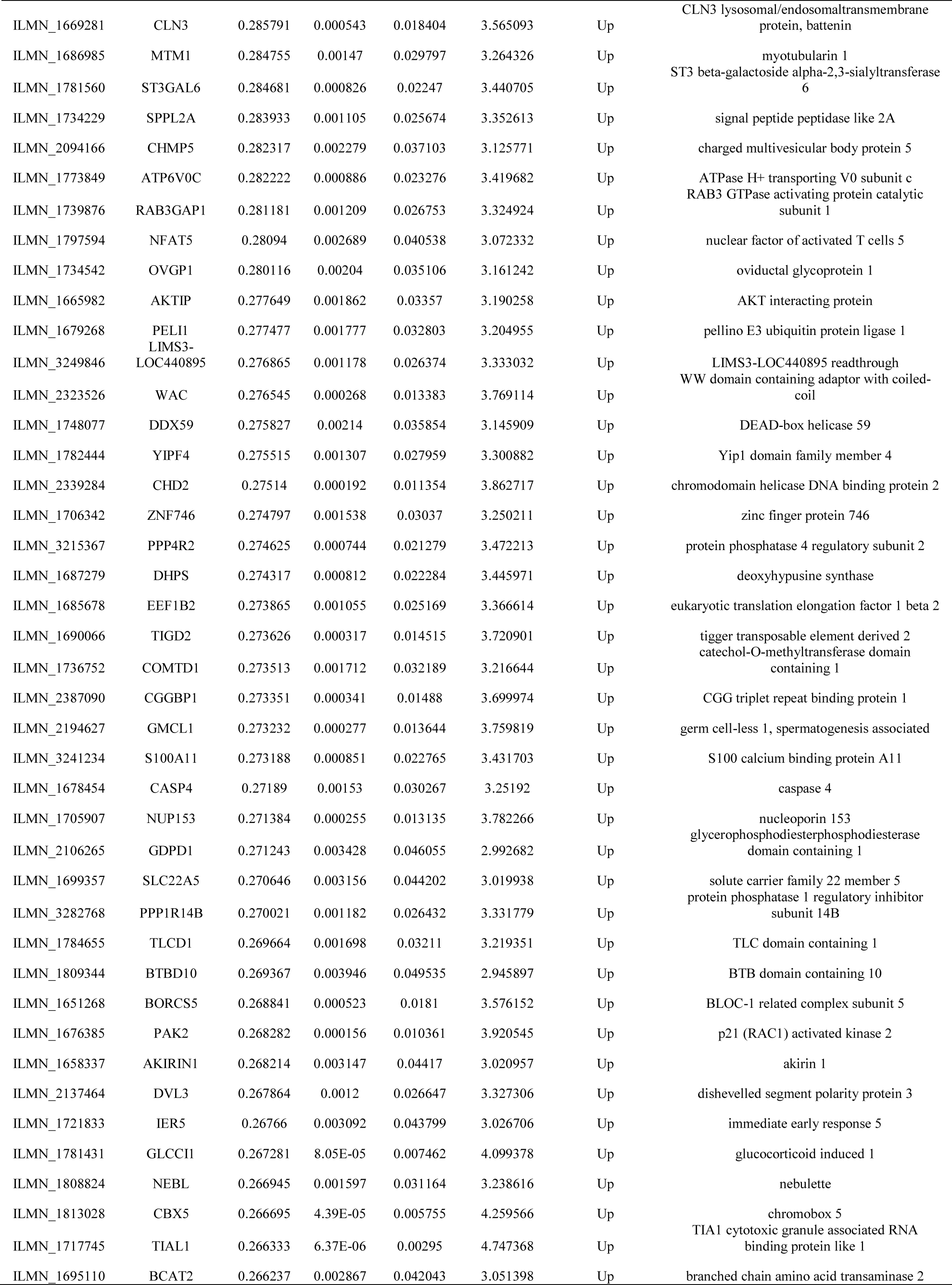
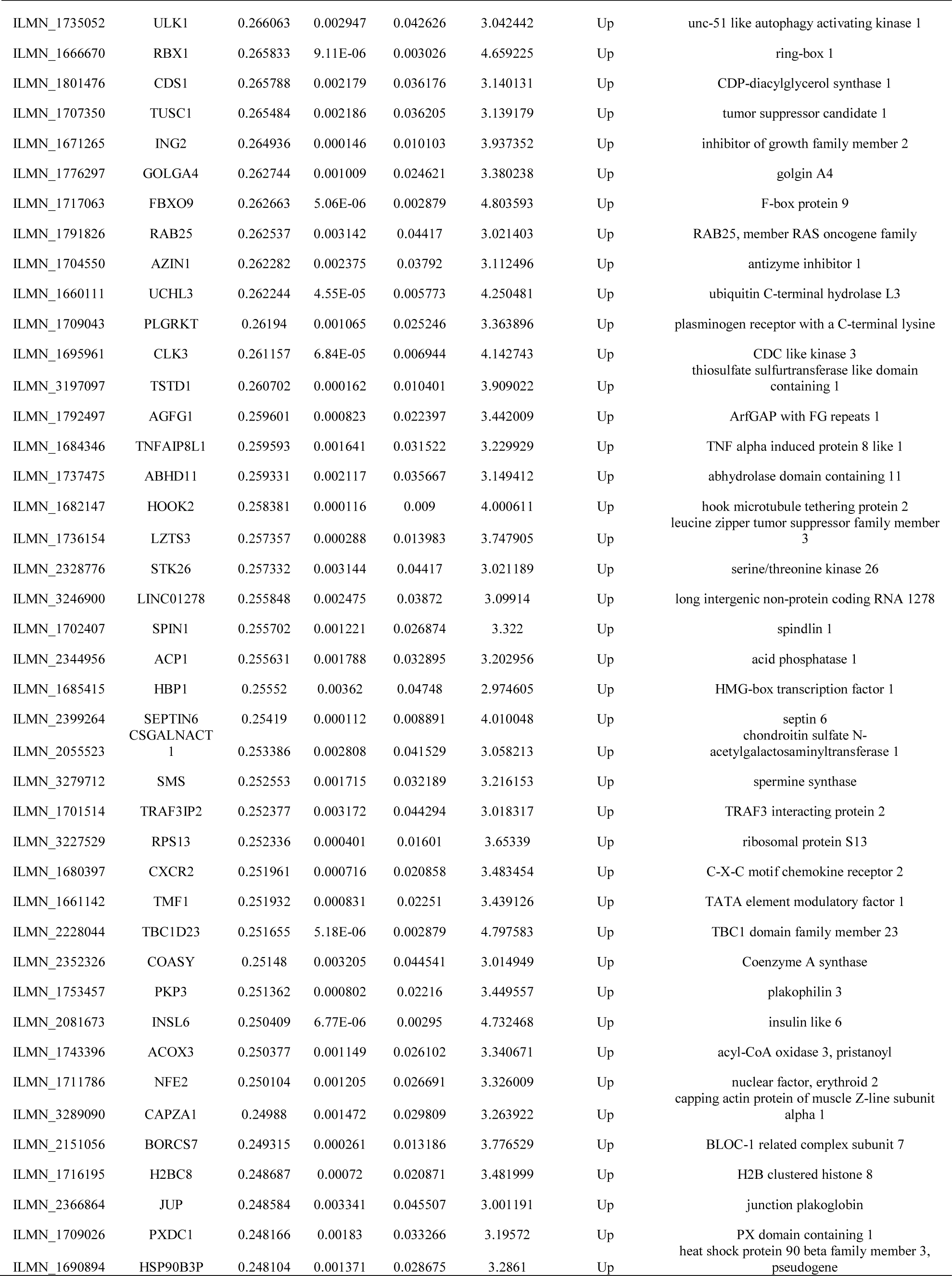
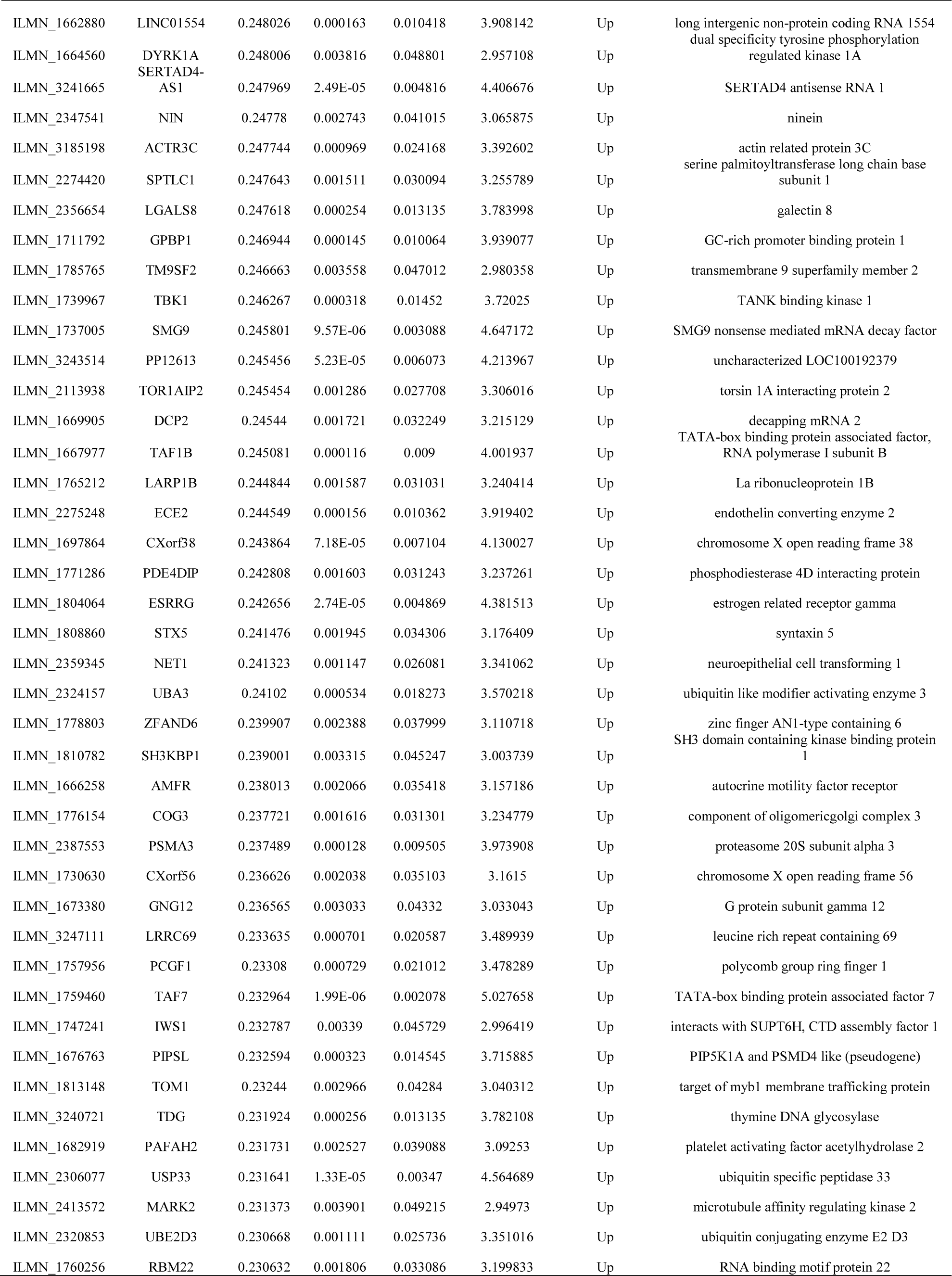
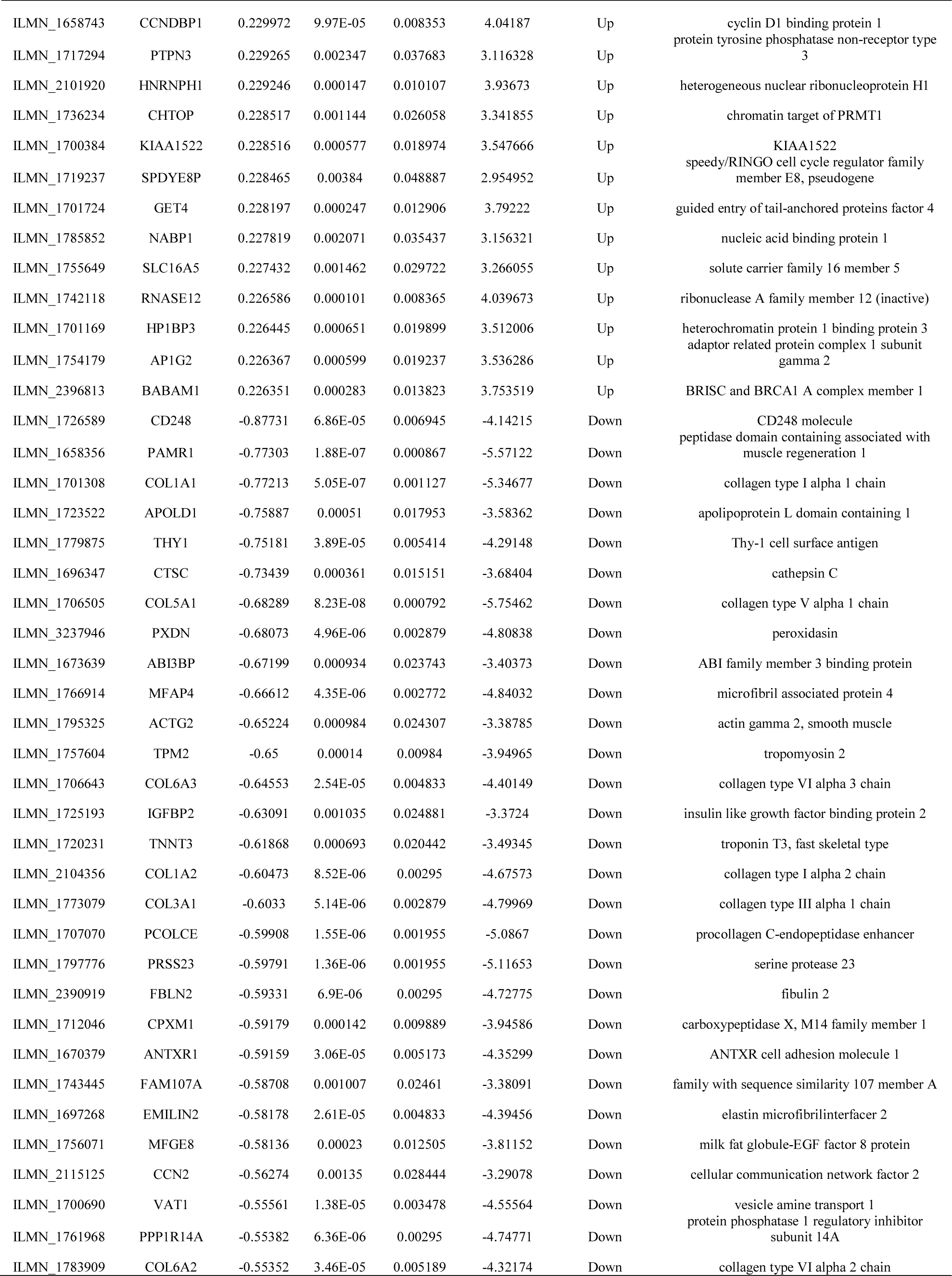
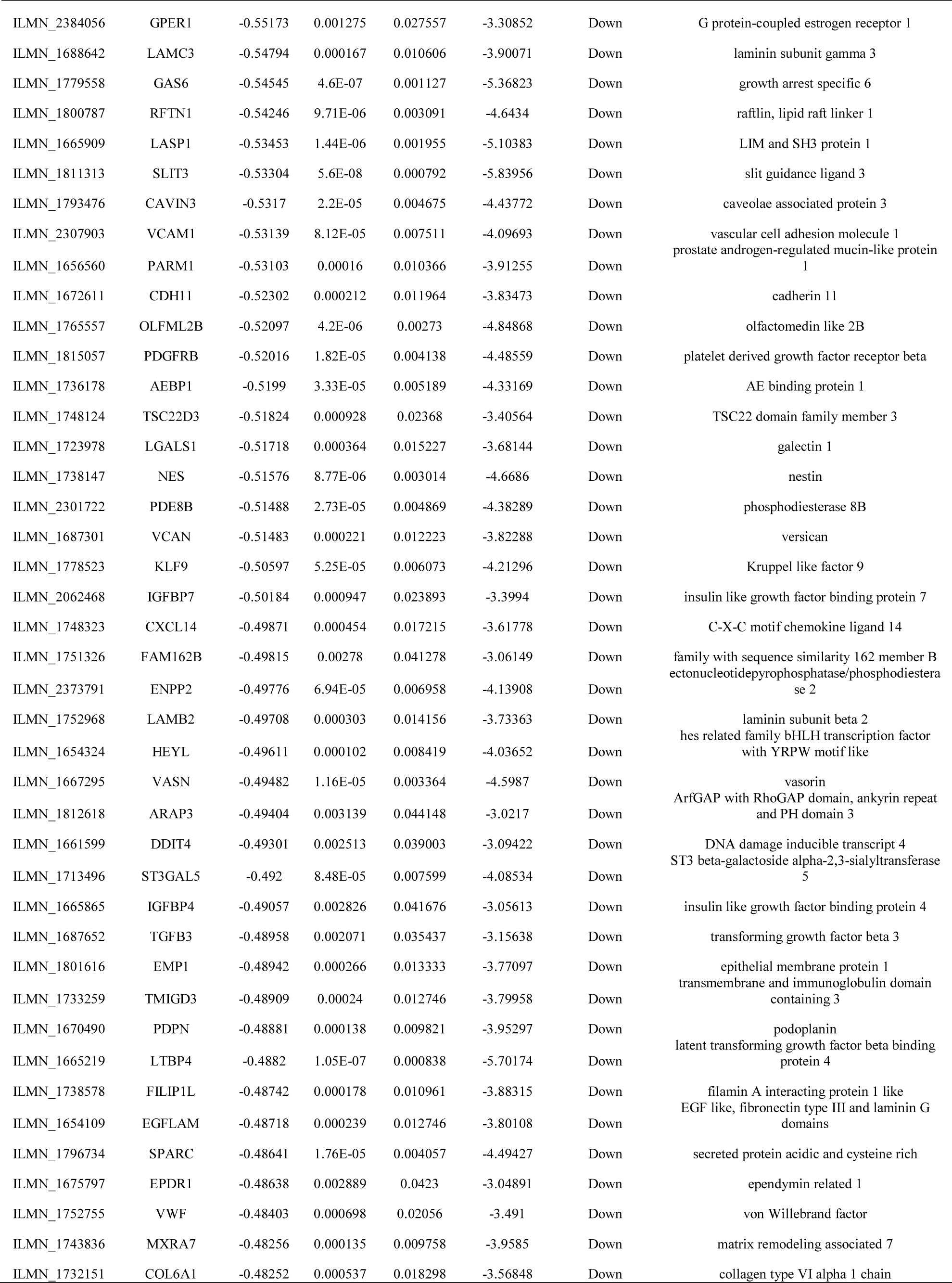
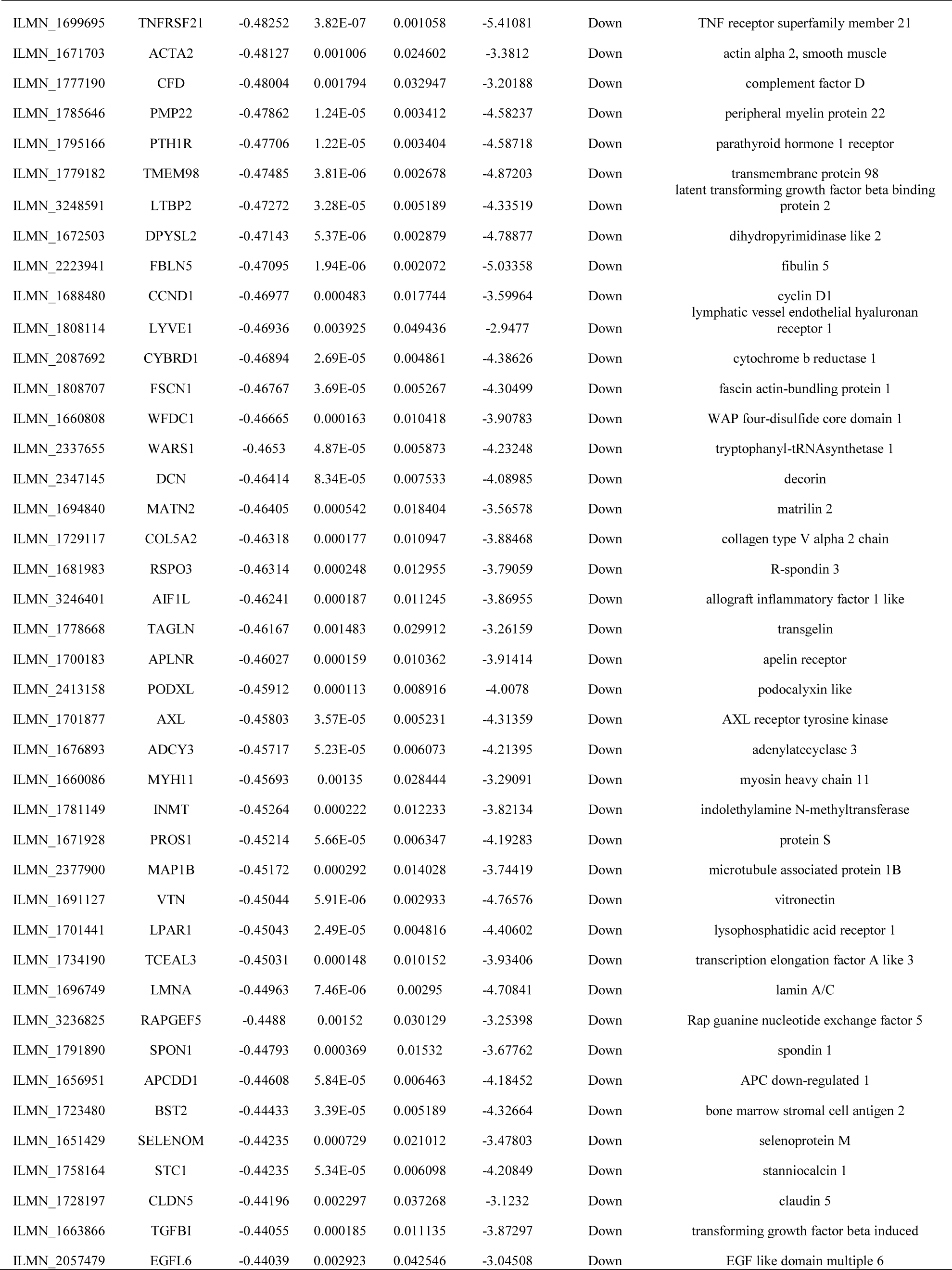
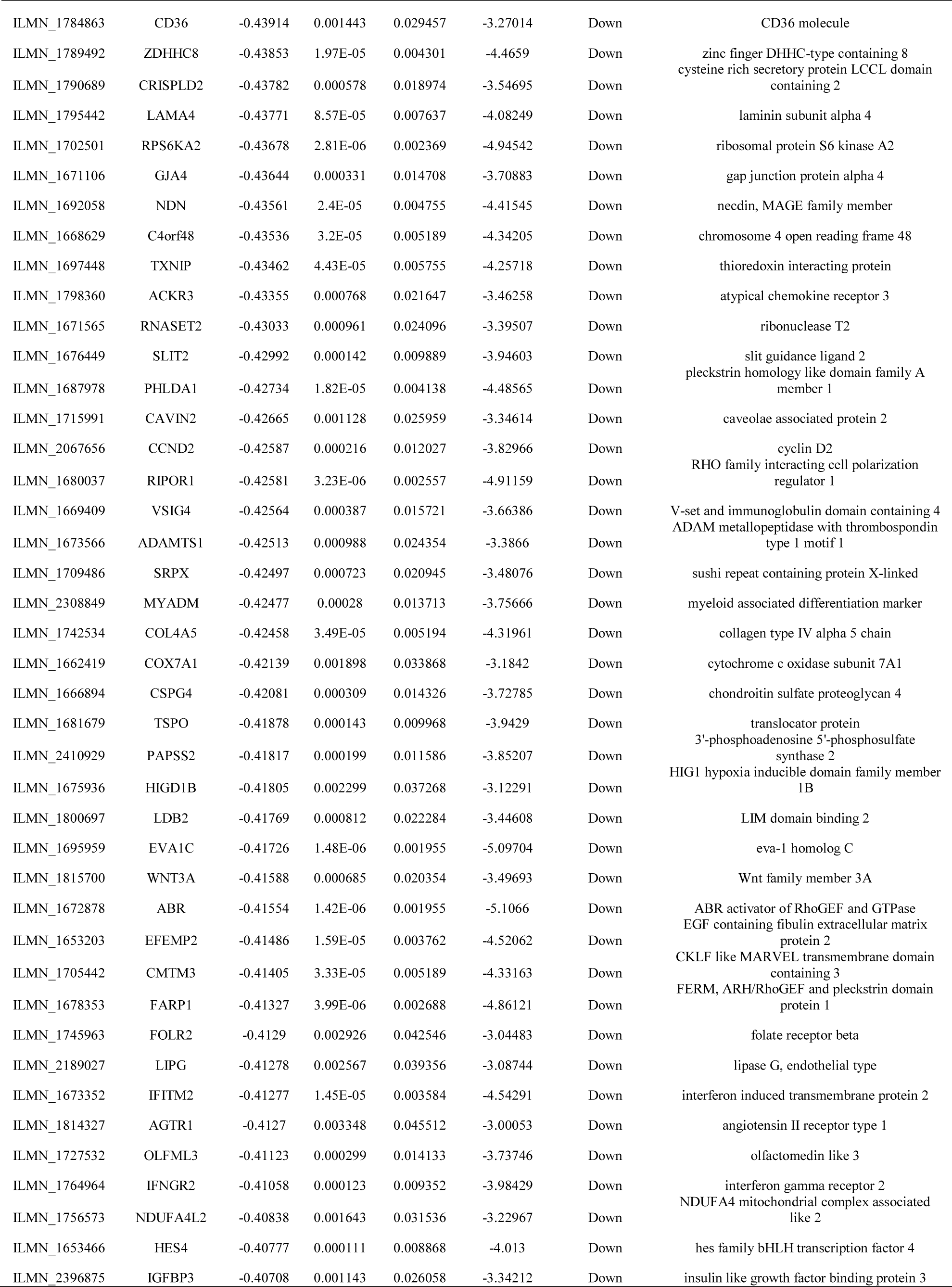
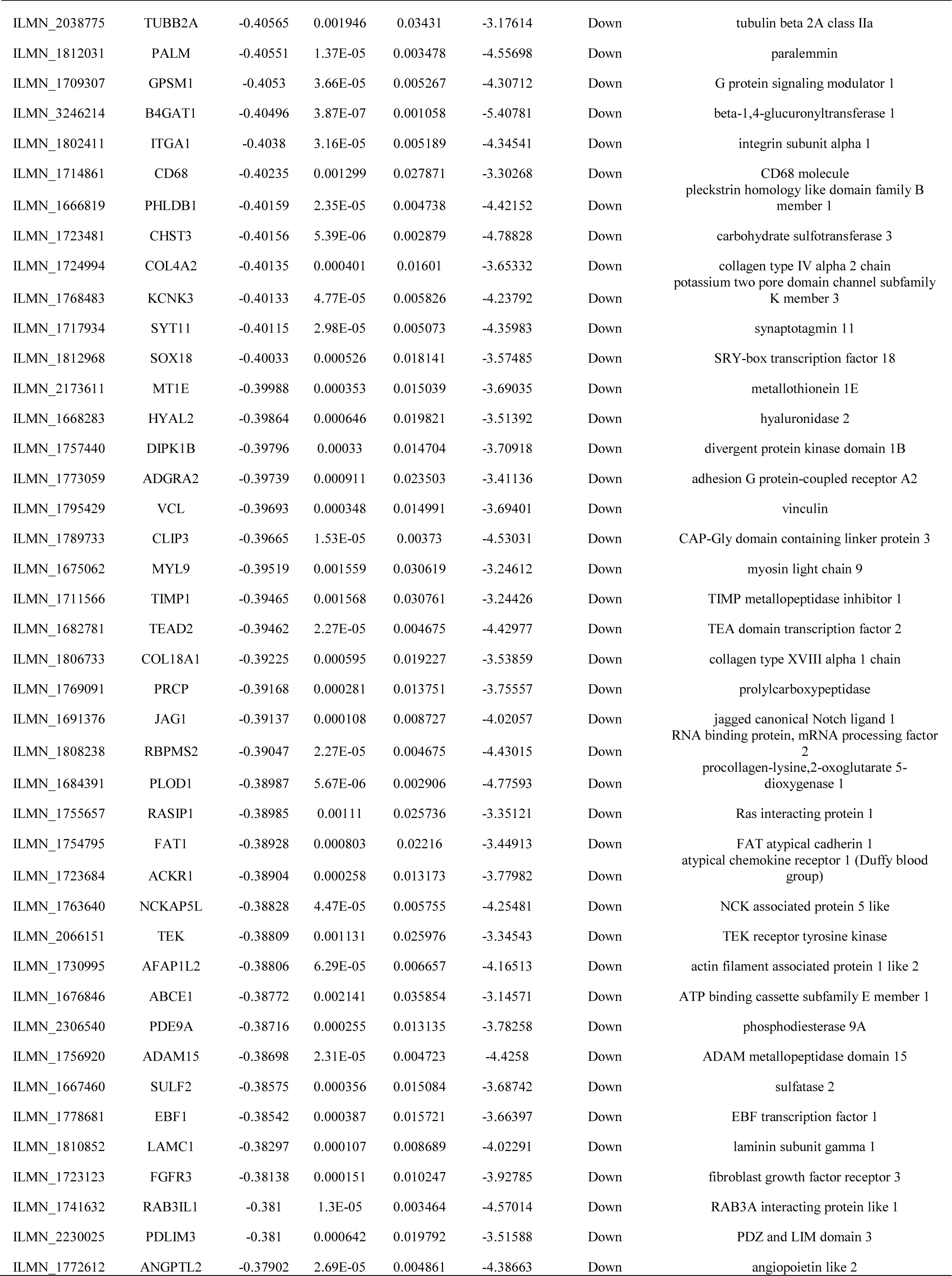
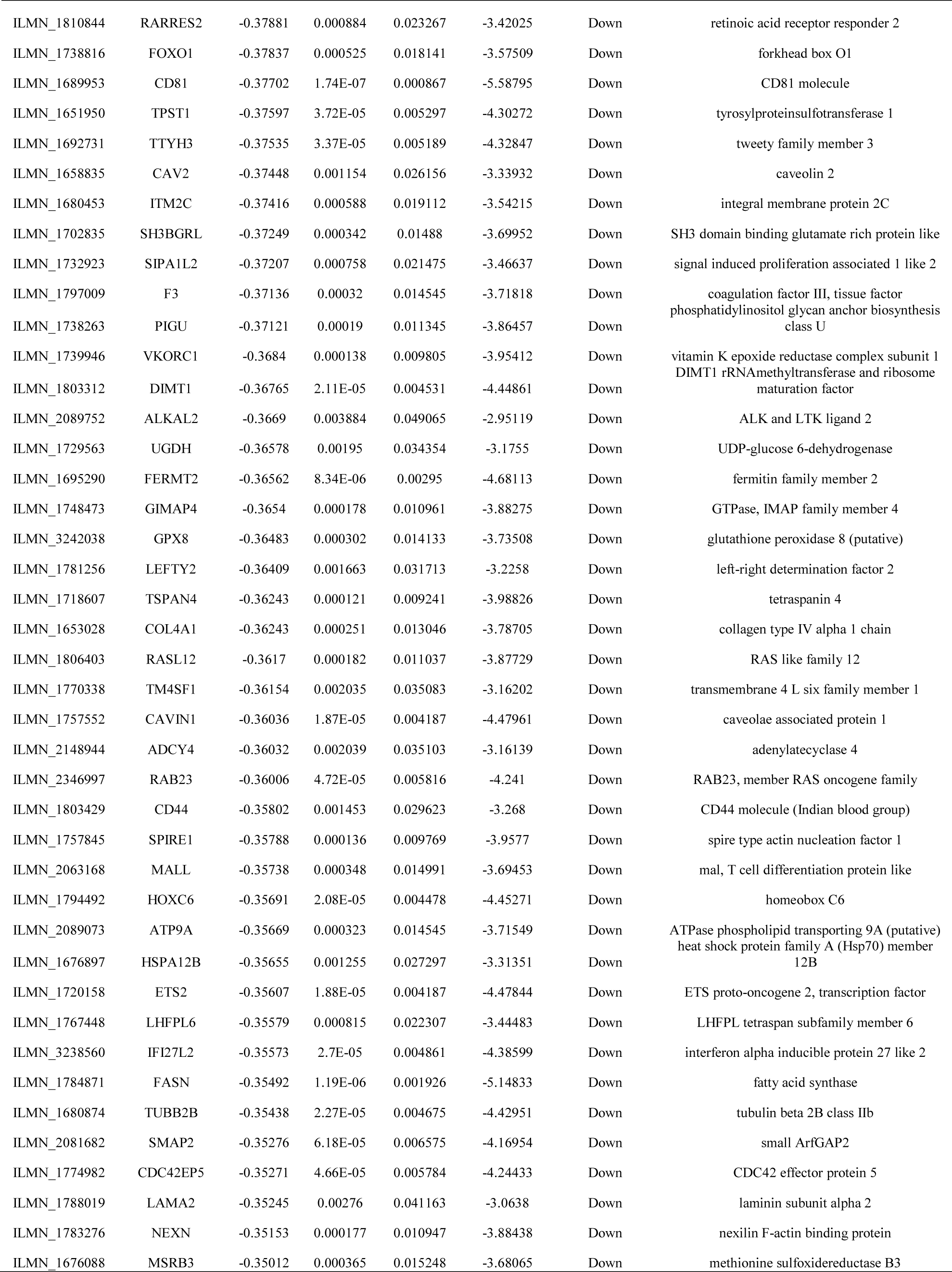
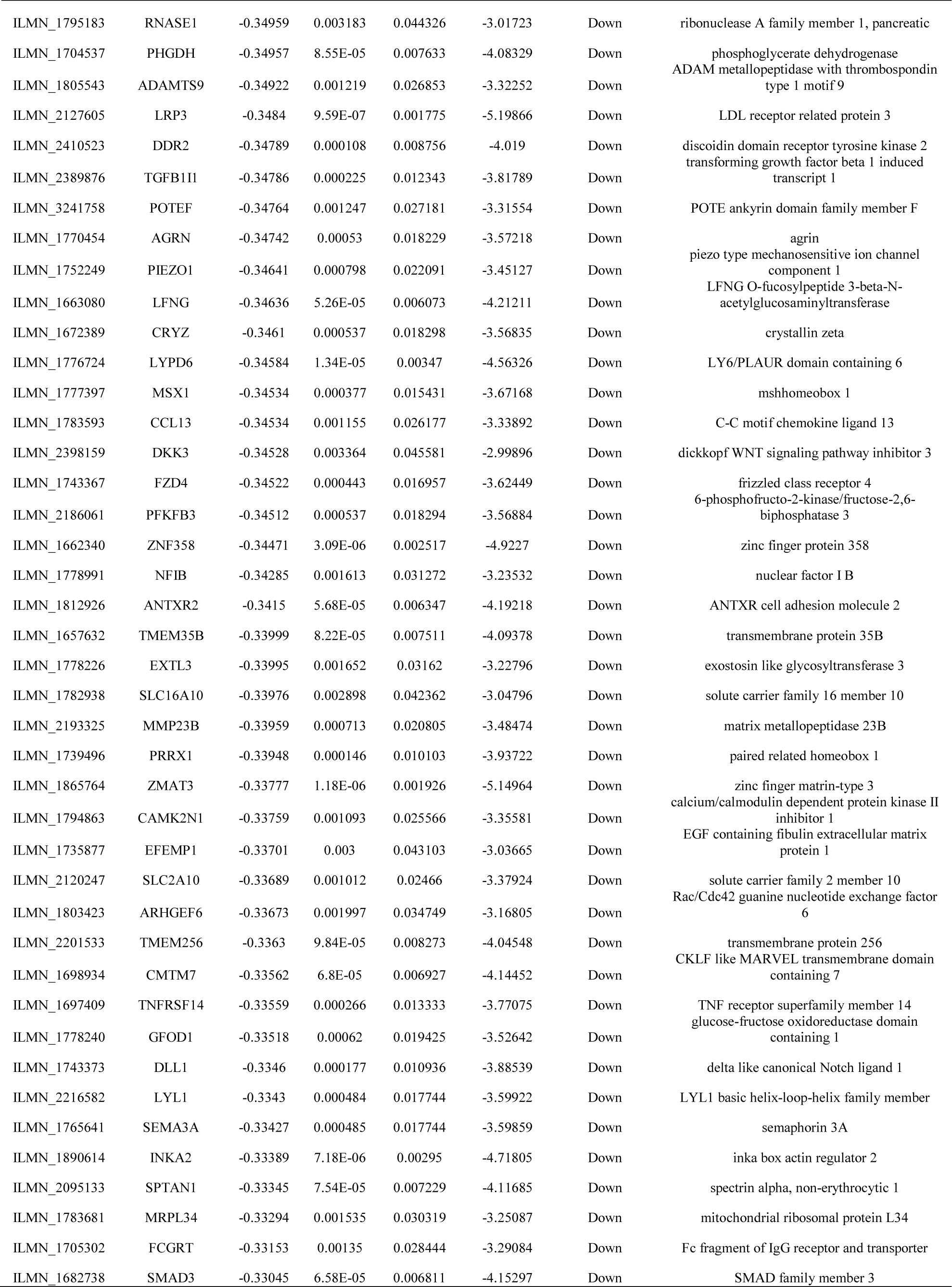
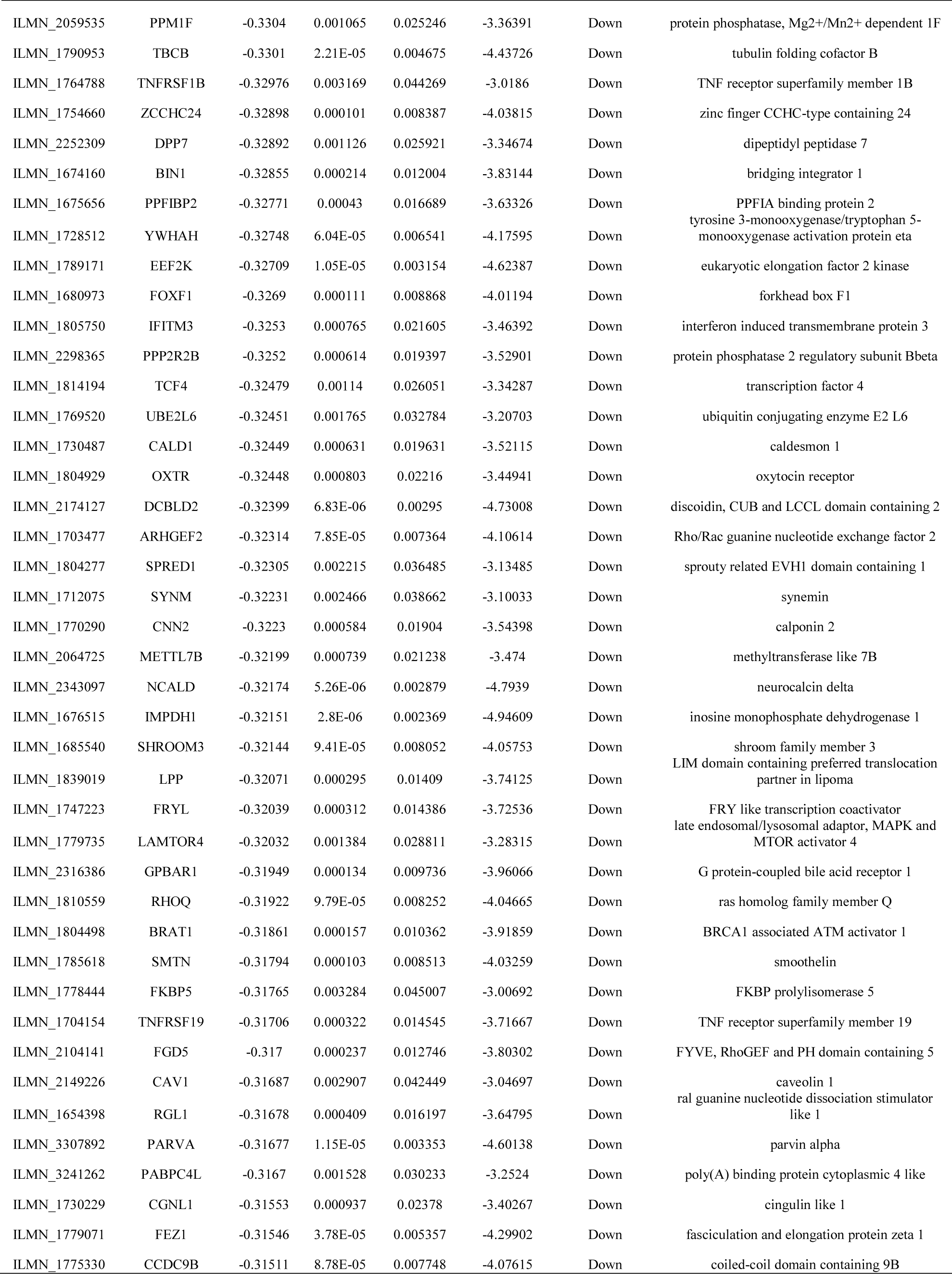
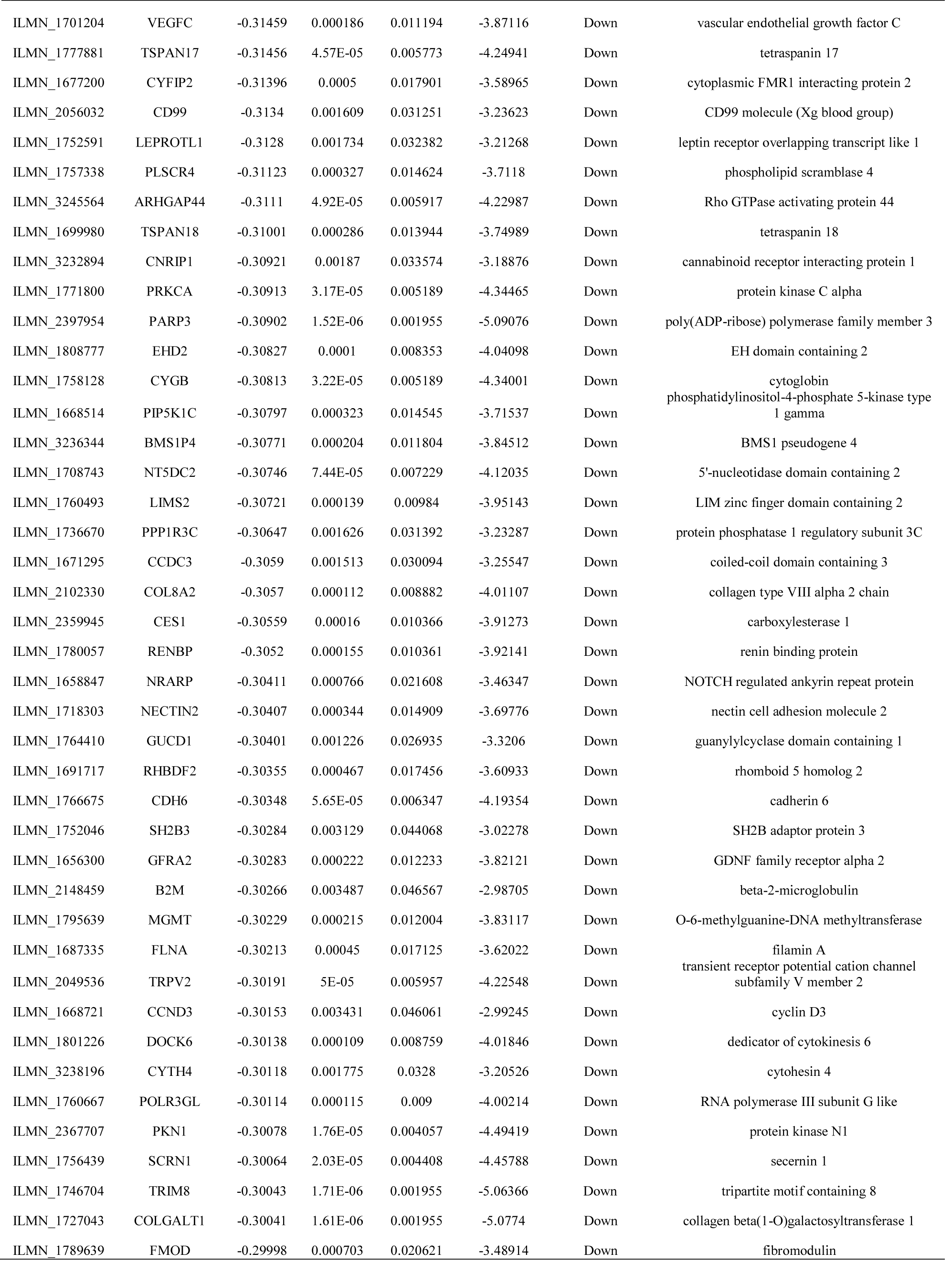
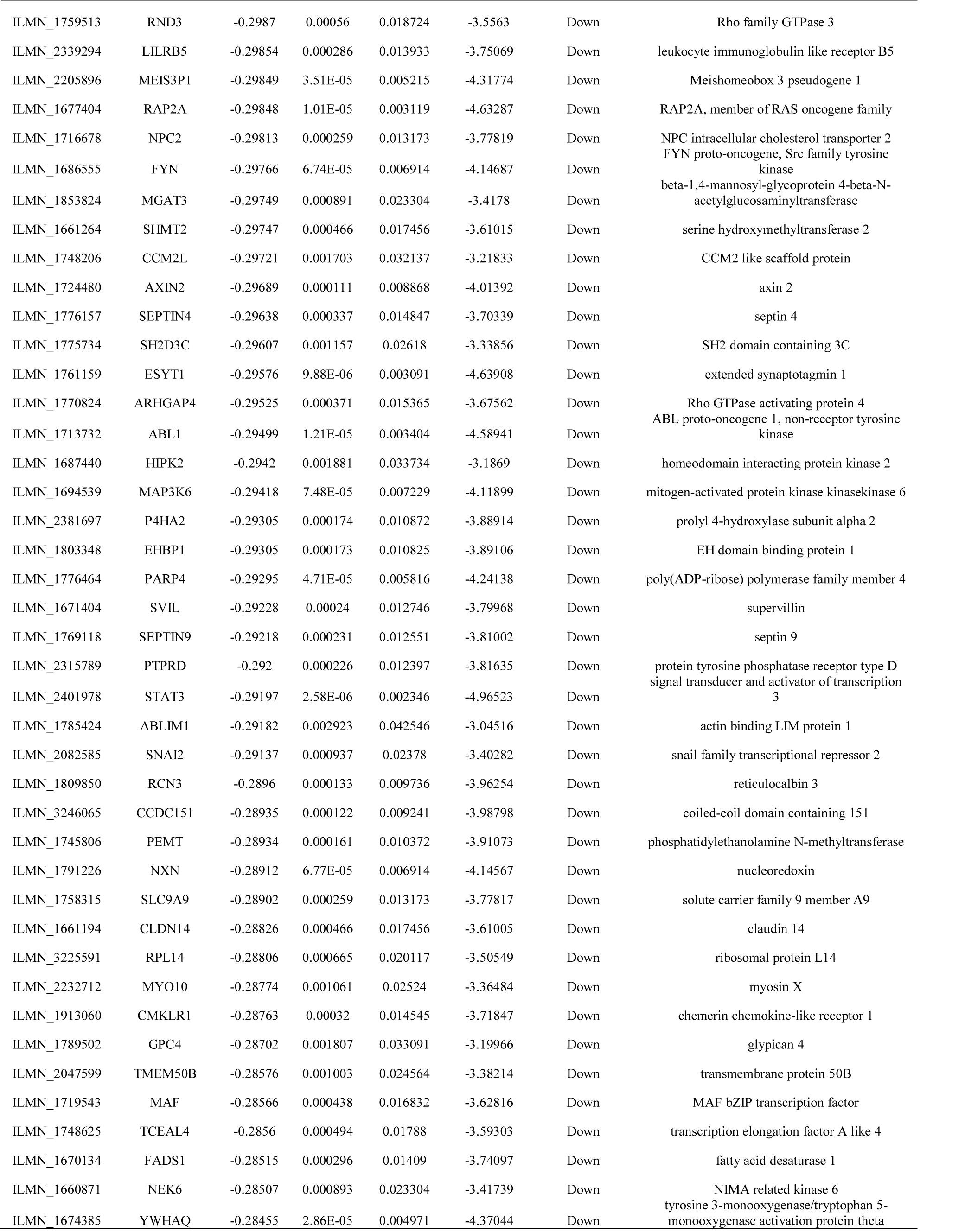
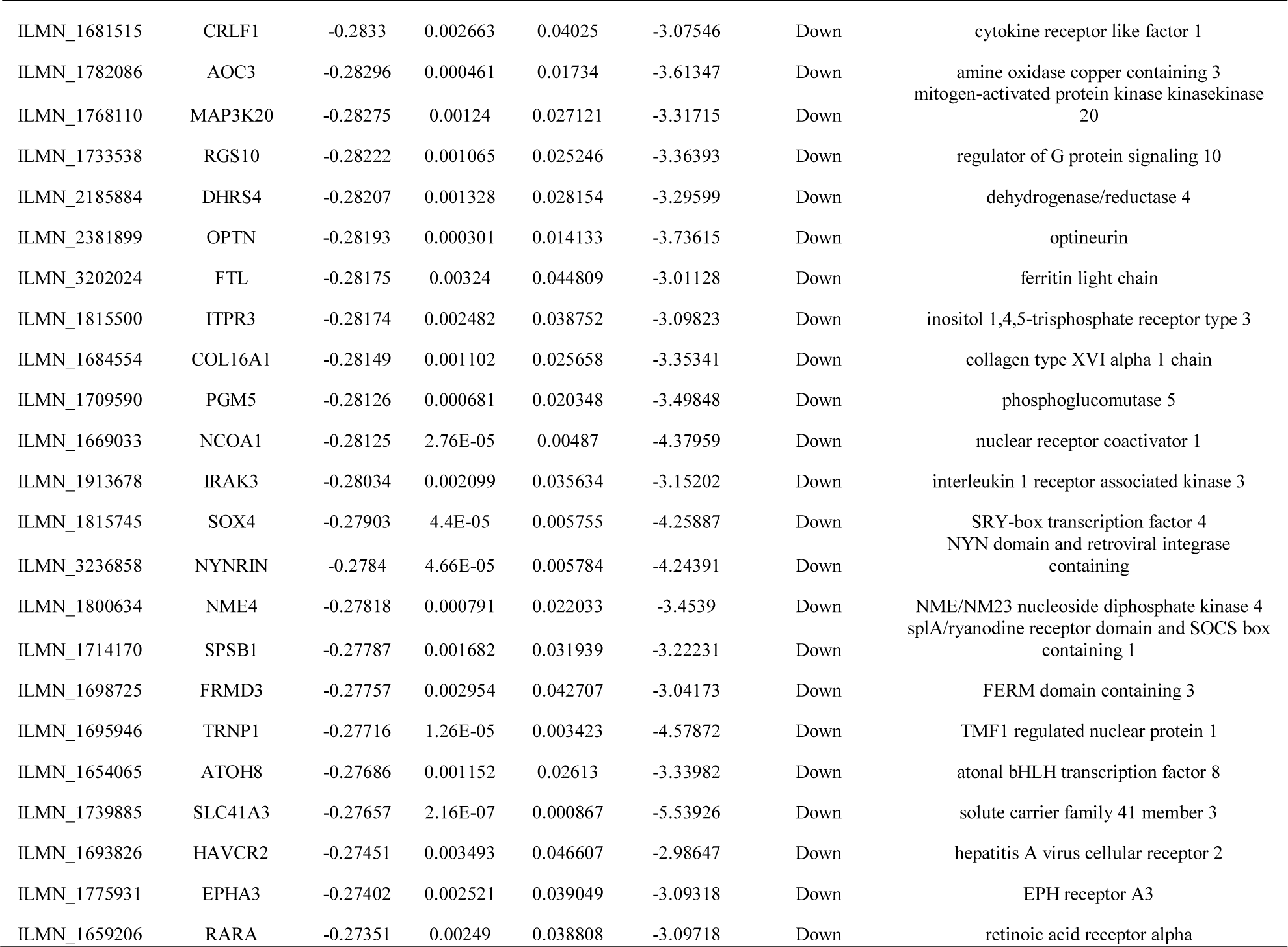
The statistical metrics for key differentially expressed genes (DEGs)

**Table 3.**
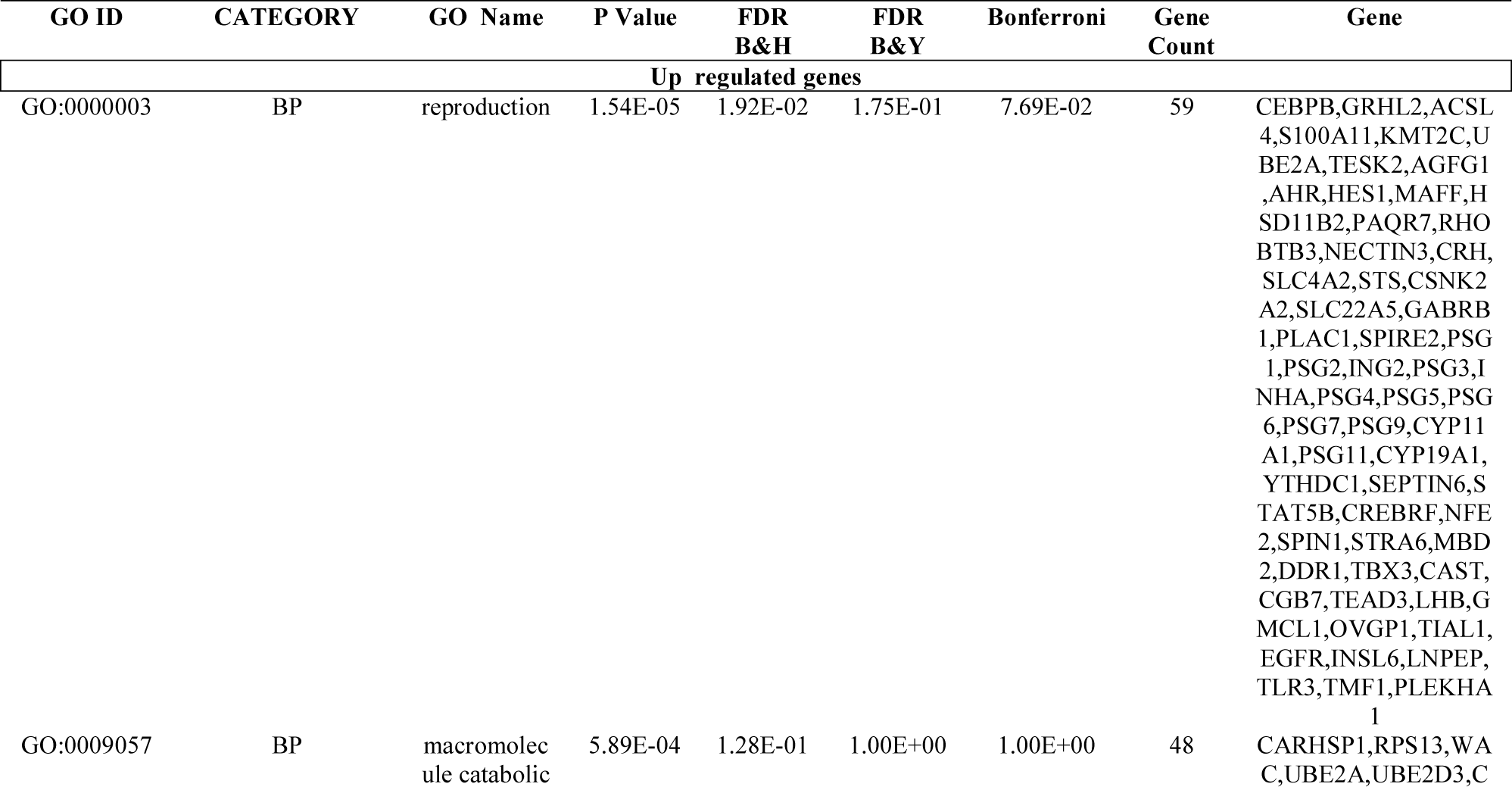
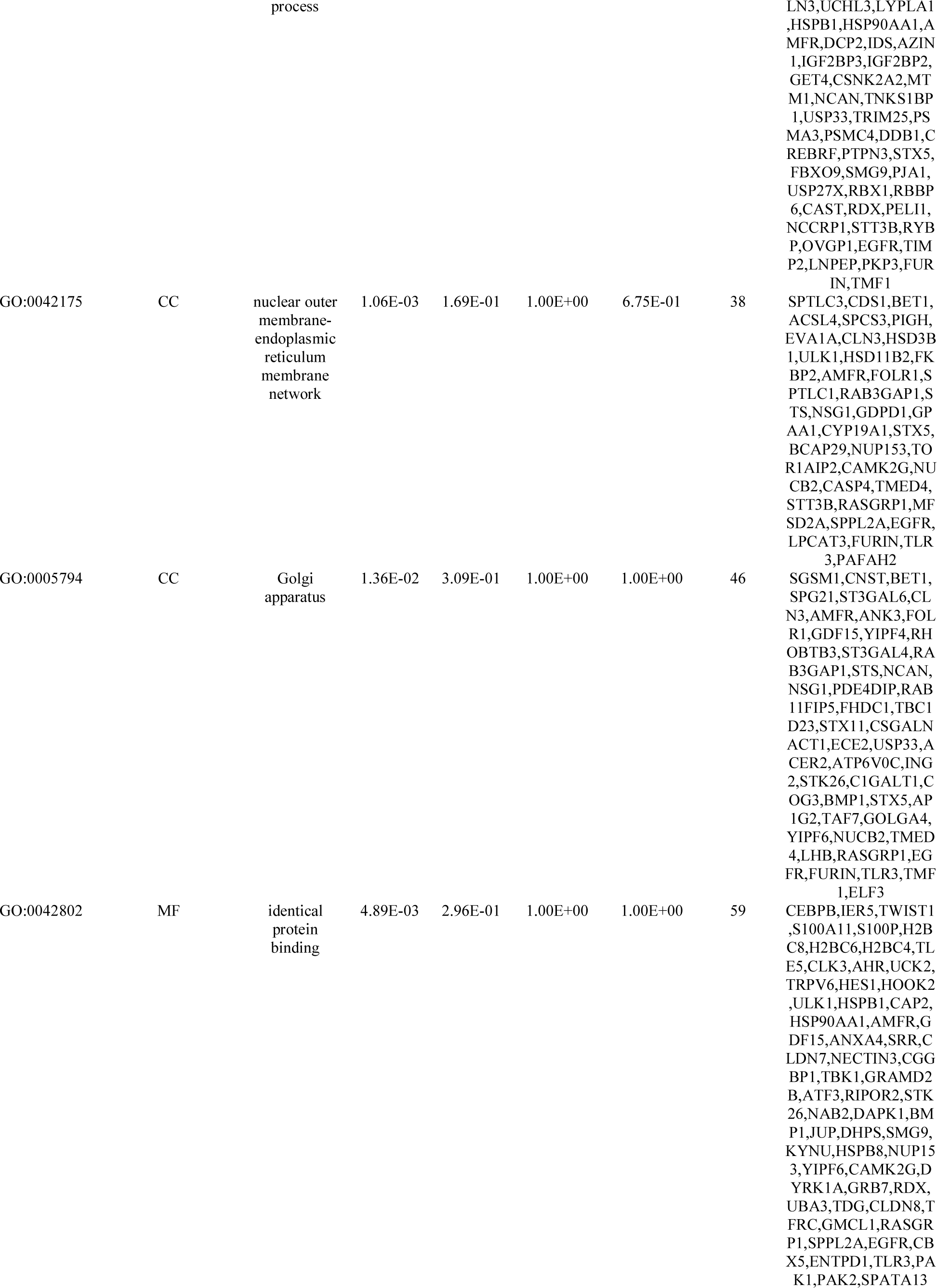
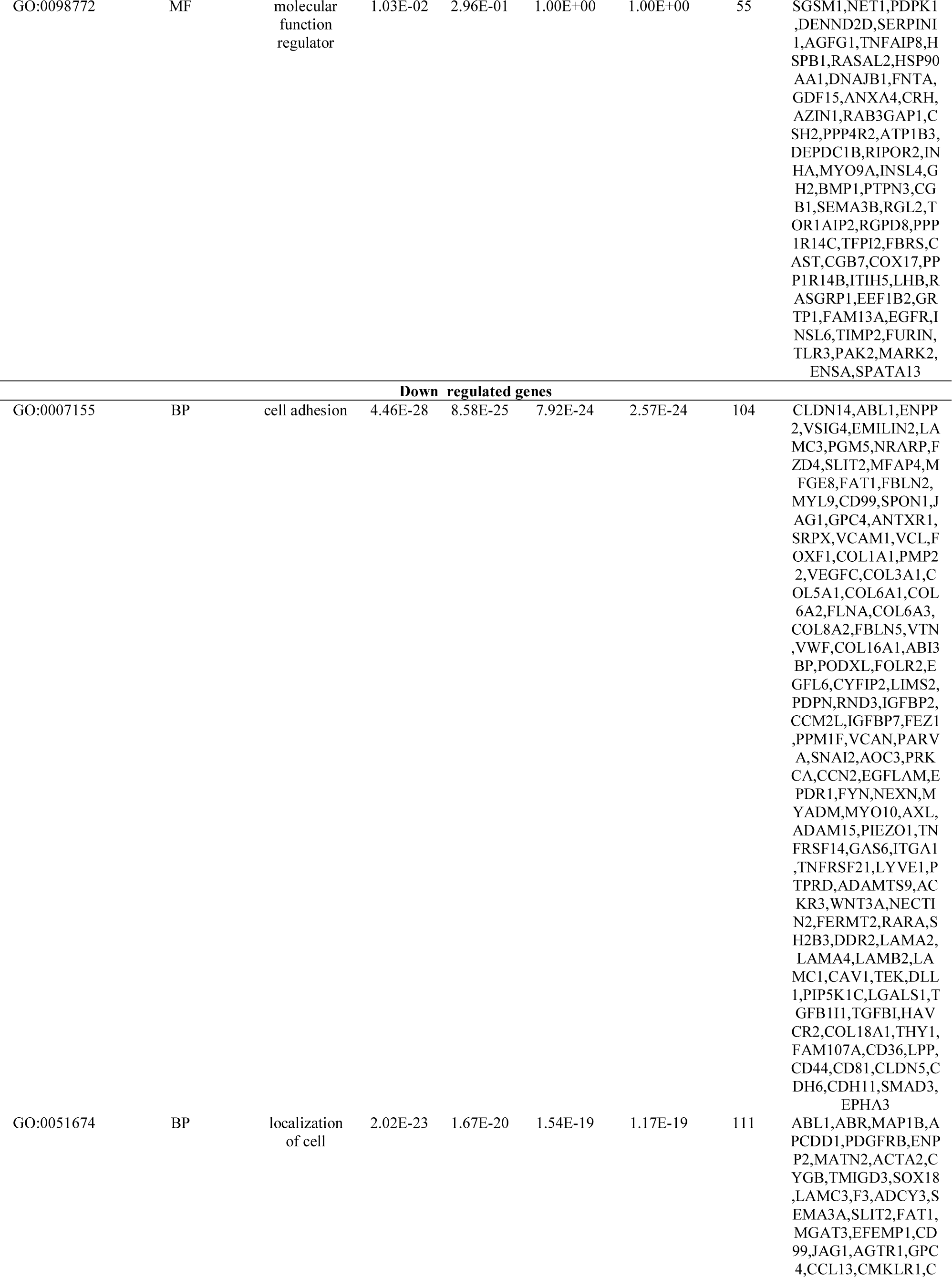
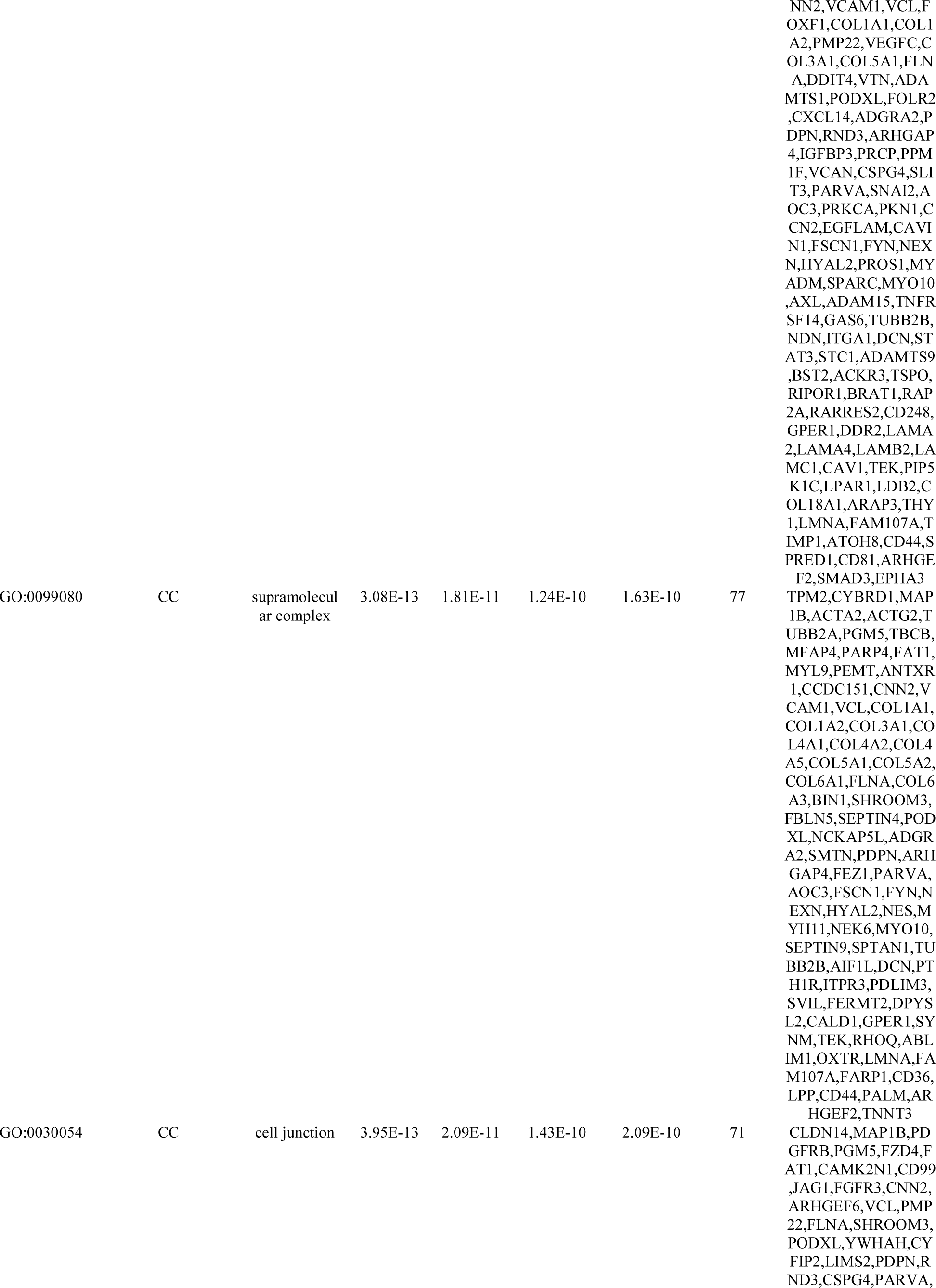
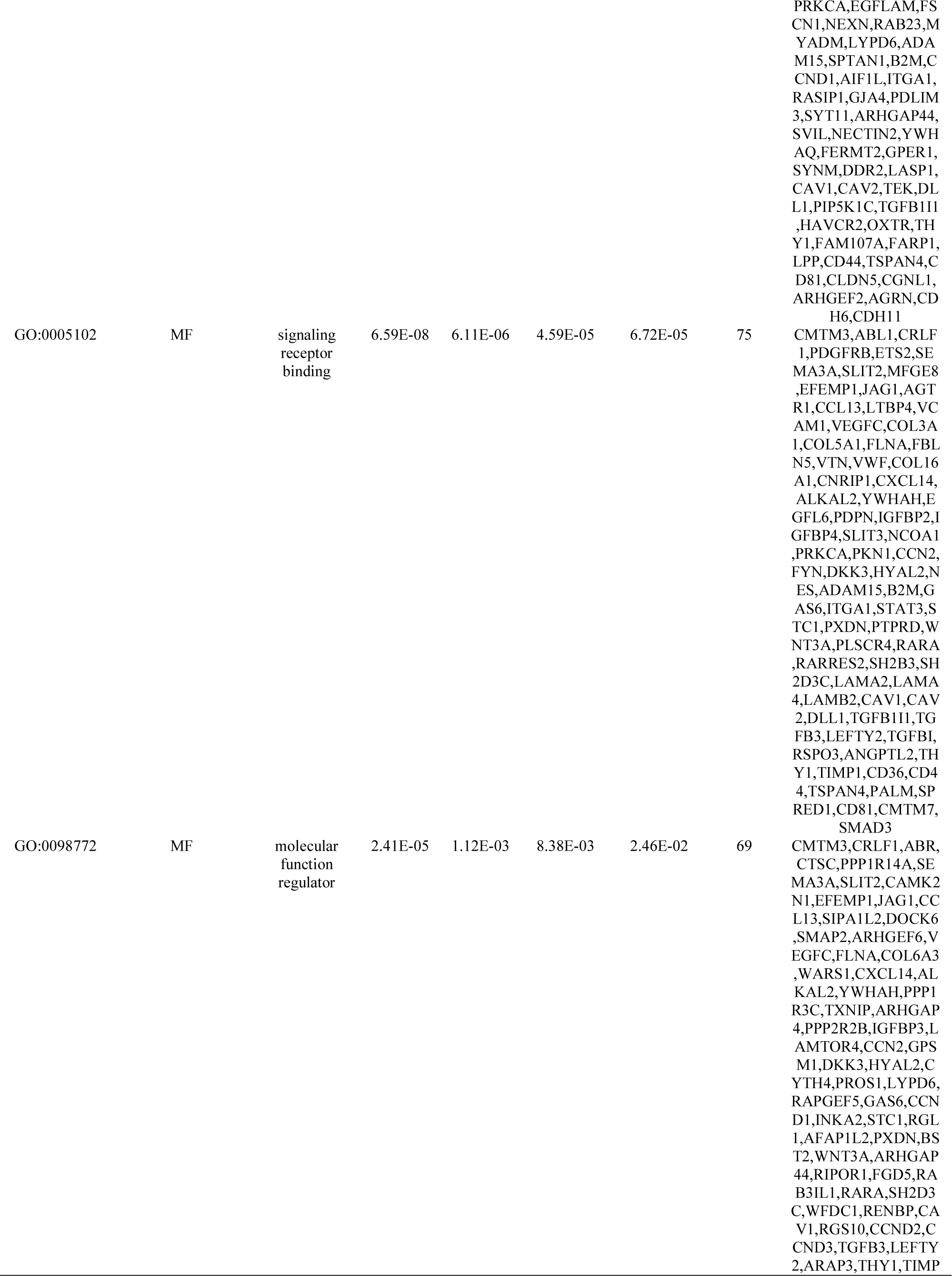
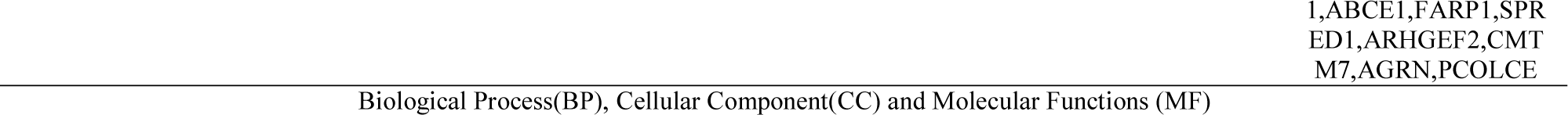
The enriched GO terms of the up and down regulated differentially expressed genes

### PPI network establishment and modules selection

By using the STRING database, the PPI network of DEGs was established and consisted of 4687 nodes and 11236 edges (Fig.3). A total of 10 hub genes were selected for key biomarker identification and are listed in Table 3. They consisted of 5 up regulated genes (HSP90AA1, EGFR, RPS13, RBX1 and PAK1) and 5 down regulated genes (FYN, ABL1, SMAD3, STAT3 and PRKCA). Then PEWCC1 was used to find clusters in the network. Four modules were calculated 2. Among them, module 1 contained 16 nodes and 32 edges, with the highest score (Fig.4A) and module 2 contained 16 nodes and 34 edges (Fig.4B). We performed the functional analysis for the top 2 modules. In functional enrichment analysis, the DEGs of module 1 were mostly enriched in post-translational protein modification, developmental biology and macromolecule catabolic process; the DEGs of module 2 in supramolecular complex and localization of cell.

**Fig. 3.**
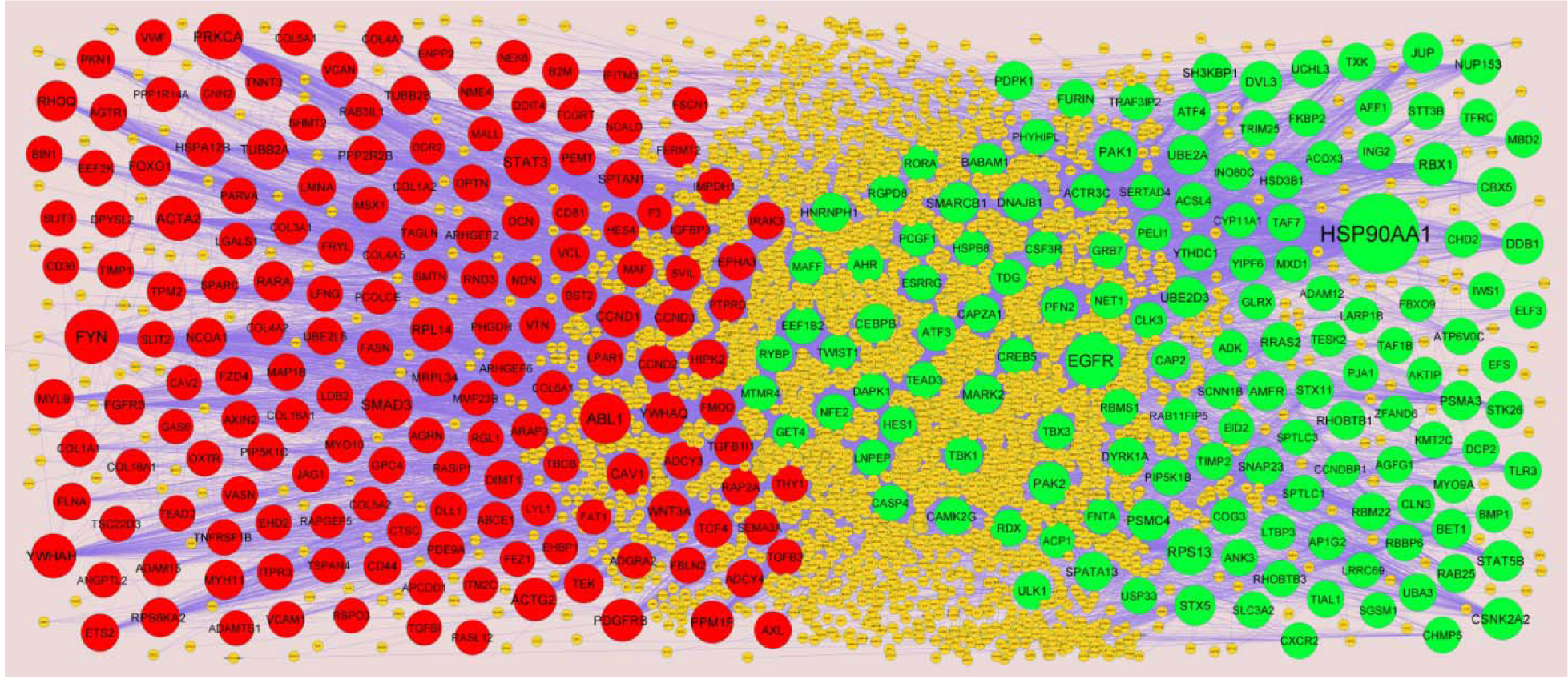
PPI network of DEGs. The PPI network of DEGs was constructed using Cytoscap. Up regulated genes are marked in green; down regulated genes are marked in red

**Fig. 4.**
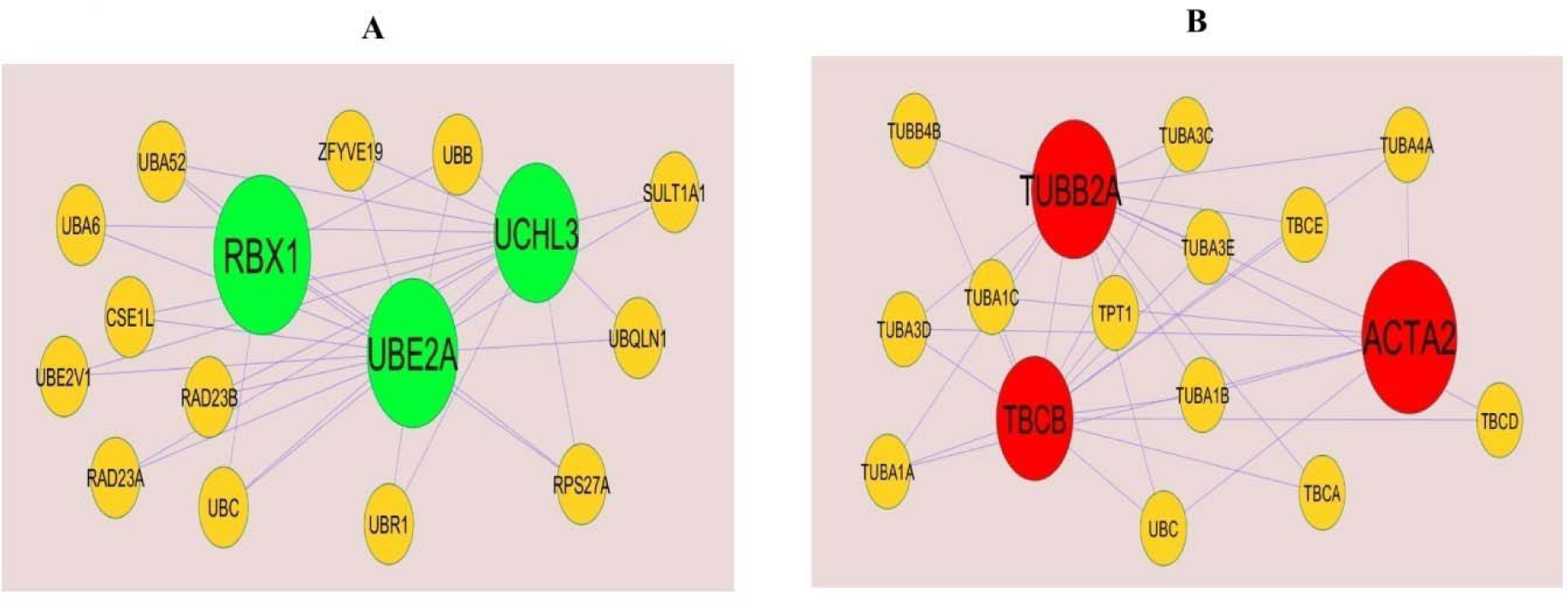
Modules of isolated form PPI of DEGs. (A) The most significant module was obtained from PPI network with 16 nodes and 32 edges for up regulated genes (B) The most significant module was obtained from PPI network with 16 nodes and 34 edges for down regulated genes. Up regulated genes are marked in green; down regulated genes are marked in red.

### miRNA-hub gene regulatory network construction

miRNet database was applied to screen the targeted miRNAs of the hub genes. Cytoscape software was used to construct the miRNA-hub gene network. As illustrated in Fig. 5, the interaction network consists of 307 hub genes and 2280 miRNAs. According to the hub genes and miRNAs in the network ranked by their degree of connectivity using Network Analyzer and are listed in Table 4. Based on the expression trend of hub genes in GDM, we found that UBE2D3 was the predicted target of hsa-mir-6127, HSP90AA1 was the predicted target of hsa-let-7d-5p, PAK2 was the predicted target of hsa-mir-8063, DDB1 was the predicted target of hsa-mir-329-3p, DVL3 was the predicted target of hsa-mir-1207-5p, FYN was the predicted target of hsa-mir-4651, ABL1 was the predicted target of hsa-mir-410-5p, SMAD3 was the predicted target of hsa-mir-222-3p, STAT3 was the predicted target of hsa-mir-29c-3p and PRKCA was the predicted target of hsa-mir-663a.

**Fig. 5.**
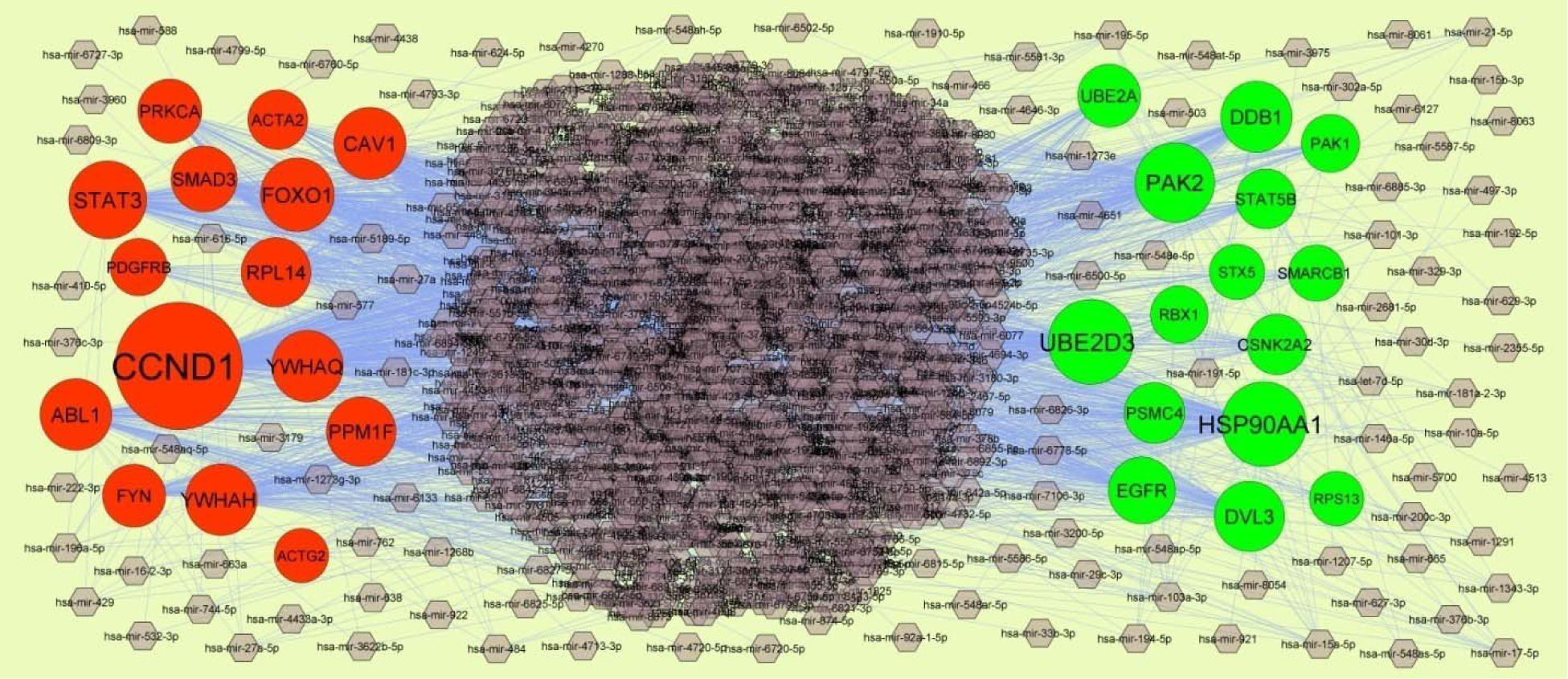
MiRNA - hub gene regulatory network. The light purple color diamond nodes represent the key miRNAs; up regulated genes are marked in green; down regulated genes are marked in red.

**Table 4.**
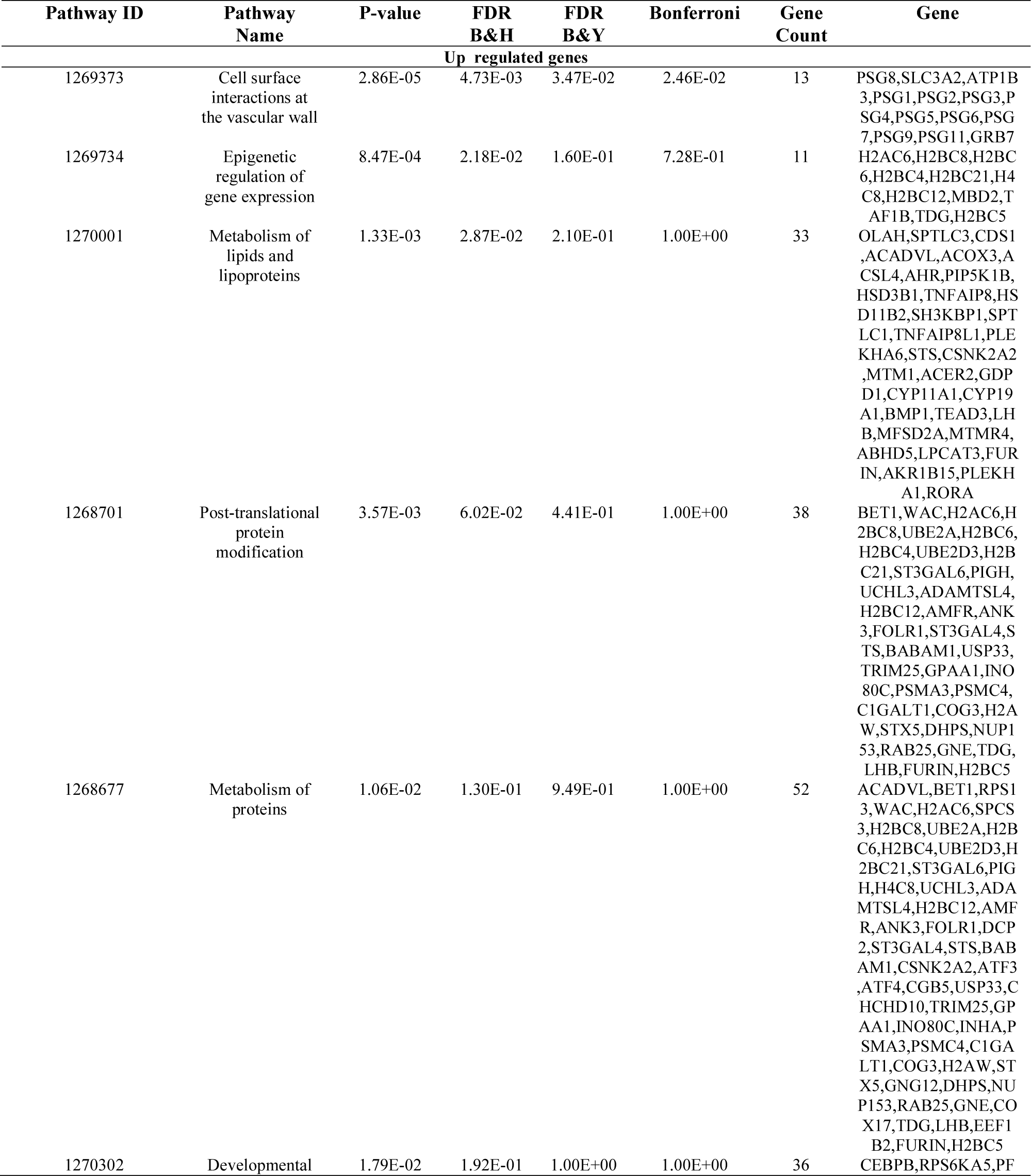
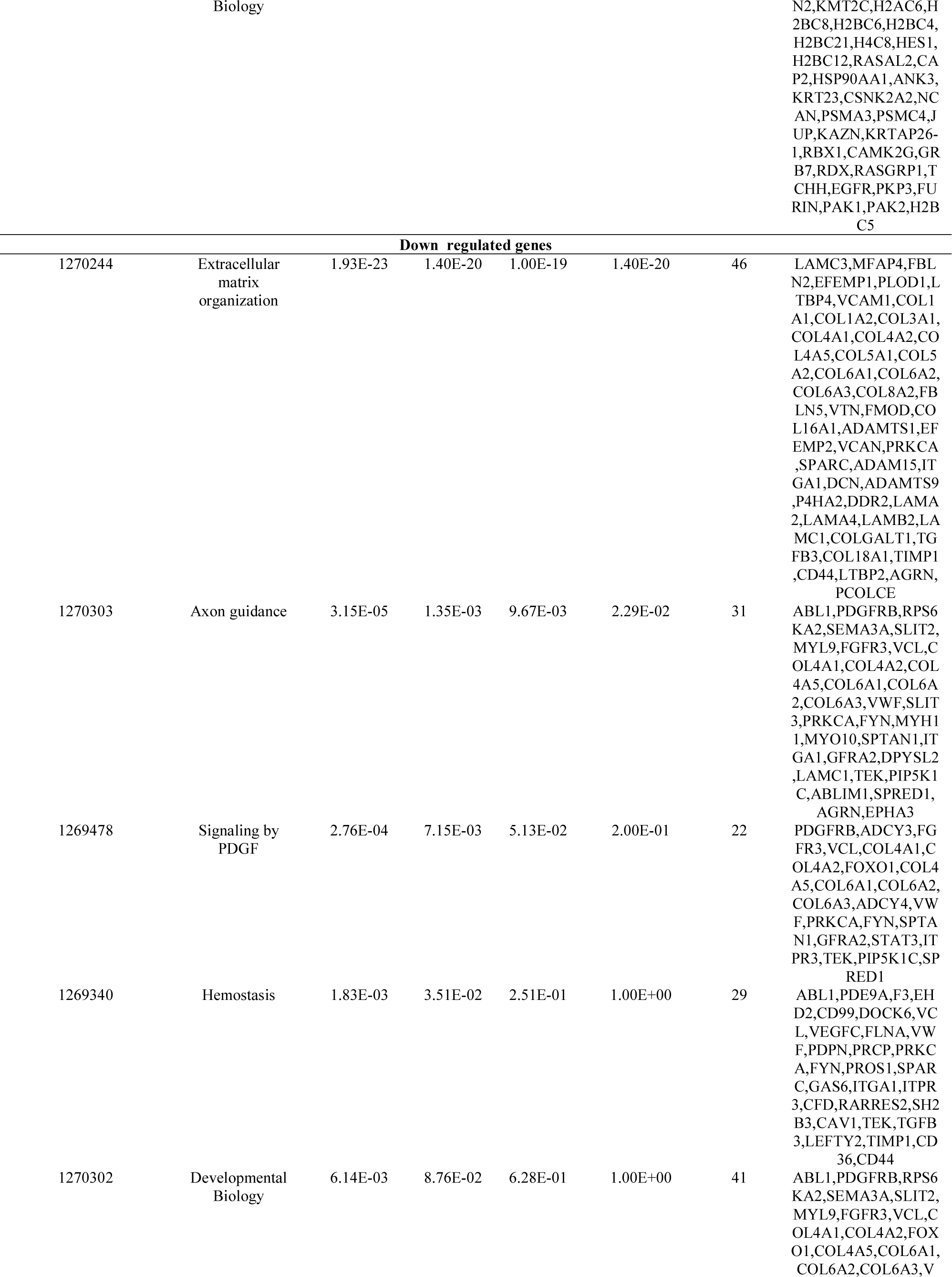
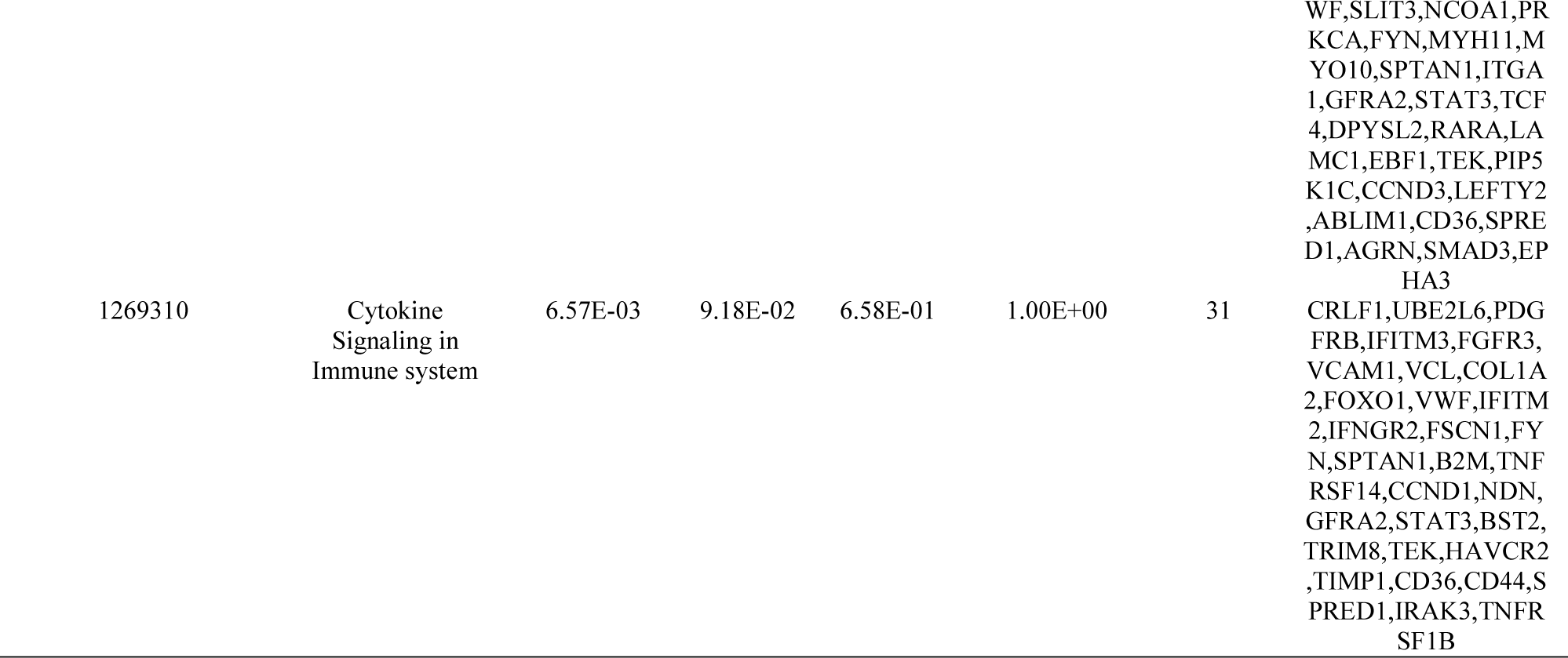
The enriched pathway terms of the up and down regulated differentially expressed genes

**Table 5.** Topology table for up and down regulated genes.

| Regulation | Node | Degree | Betweenness | Stress | Closeness |
| --- | --- | --- | --- | --- | --- |
| Up | HSP90AA1 | 655 | 0.22721 | 81351654 | 0.412863 |
| Up | EGFR | 324 | 0.081831 | 21358048 | 0.396682 |
| Up | RPS13 | 176 | 0.040553 | 21024400 | 0.332742 |
| Up | RBX1 | 132 | 0.02978 | 9542636 | 0.3408 |
| Up | PAK1 | 115 | 0.016755 | 4561194 | 0.37158 |
| Up | CSNK2A2 | 112 | 0.026157 | 6172590 | 0.354758 |
| Up | PAK2 | 107 | 0.012051 | 4334042 | 0.349858 |
| Up | DDB1 | 105 | 0.024917 | 6240978 | 0.340602 |
| Up | PSMC4 | 101 | 0.019368 | 3810802 | 0.348739 |
| Up | DVL3 | 99 | 0.017256 | 5360158 | 0.344762 |
| Up | UBE2D3 | 96 | 0.018158 | 6872298 | 0.333975 |
| Up | SMARCB1 | 90 | 0.021042 | 7085456 | 0.333215 |
| Up | STAT5B | 89 | 0.005633 | 1782886 | 0.349937 |
| Up | STX5 | 86 | 0.021729 | 5783250 | 0.327692 |
| Up | UBE2A | 86 | 0.01665 | 6800442 | 0.330652 |
| Up | NUP153 | 81 | 0.019493 | 3839268 | 0.33388 |
| Up | JUP | 79 | 0.012614 | 3875882 | 0.320148 |
| Up | PSMA3 | 78 | 0.009276 | 2735940 | 0.330535 |
| Up | SH3KBP1 | 76 | 0.011315 | 2648130 | 0.345321 |
| Up | HSPB1 | 69 | 0.012593 | 3676246 | 0.34167 |
| Up | BET1 | 67 | 0.013941 | 4543480 | 0.325711 |
| Up | AMFR | 66 | 0.018899 | 2411936 | 0.345907 |
| Up | RRAS2 | 64 | 0.00438 | 2166928 | 0.301914 |
| Up | MARK2 | 63 | 0.012107 | 3287228 | 0.338315 |
| Up | CBX5 | 62 | 0.015481 | 3171408 | 0.327807 |
| Up | CEBPB | 62 | 0.007601 | 2245984 | 0.350172 |
| Up | PDPK1 | 62 | 0.005919 | 1612918 | 0.342169 |
| Up | HNRNPH1 | 59 | 0.013608 | 2423406 | 0.335625 |
| Up | DNAJB1 | 58 | 0.00611 | 1362926 | 0.344205 |
| Up | ATF3 | 57 | 0.005226 | 1555506 | 0.352755 |
| Up | SPATA13 | 56 | 0.00717 | 2217896 | 0.295759 |
| Up | FURIN | 55 | 0.017306 | 2333572 | 0.332317 |
| Up | RHOBTB1 | 54 | 0.005265 | 891404 | 0.309798 |
| Up | SNAP23 | 52 | 0.010301 | 2133980 | 0.326551 |
| Up | STX11 | 49 | 0.006356 | 1012388 | 0.276999 |
| Up | TBK1 | 48 | 0.00946 | 1612462 | 0.346521 |
| Up | RAB25 | 48 | 0.00796 | 4209932 | 0.282579 |
| Up | ING2 | 42 | 0.006848 | 2338492 | 0.325236 |
| Up | ULK1 | 42 | 0.007643 | 7120856 | 0.274292 |
| Up | UCHL3 | 42 | 0.007629 | 1994146 | 0.325779 |
| Up | PFN2 | 41 | 0.004889 | 1200798 | 0.329235 |
| Up | CAP2 | 40 | 0.002011 | 661156 | 0.288156 |
| Up | UBA3 | 40 | 0.005749 | 1625200 | 0.326187 |
| Up | DYRK1A | 40 | 0.004394 | 1121650 | 0.309634 |
| Up | RBM22 | 39 | 0.01146 | 1346380 | 0.322616 |
| Up | TXK | 38 | 0.001513 | 482866 | 0.300558 |
| Up | TAF7 | 38 | 0.010147 | 1476134 | 0.322972 |
| Up | CAMK2G | 38 | 0.007152 | 1050770 | 0.341969 |
| Up | ATF4 | 37 | 0.006742 | 1255258 | 0.328036 |
| Up | TLR3 | 36 | 0.004686 | 1054636 | 0.305217 |
| Up | MYO9A | 36 | 8.05E-04 | 406980 | 0.28752 |
| Up | BABAM1 | 36 | 0.008431 | 1423346 | 0.324044 |
| Up | STK26 | 36 | 0.008804 | 1319200 | 0.323842 |
| Up | ACTR3C | 35 | 0.002617 | 659206 | 0.282732 |
| Up | CREB5 | 35 | 0.001154 | 566084 | 0.298035 |
| Up | NET1 | 33 | 0.003076 | 787876 | 0.327189 |
| Up | EEF1B2 | 32 | 0.00465 | 1484558 | 0.32501 |
| Up | AHR | 32 | 0.003156 | 847220 | 0.347858 |
| Up | PIP5K1B | 31 | 0.00183 | 957308 | 0.326391 |
| Up | CLN3 | 31 | 0.007159 | 992822 | 0.323485 |
| Up | HES1 | 30 | 0.0035 | 796872 | 0.300115 |
| Up | RBBP6 | 29 | 0.008094 | 1087968 | 0.324089 |
| Up | AP1G2 | 29 | 0.008892 | 933344 | 0.321377 |
| Up | RDX | 29 | 0.003378 | 953914 | 0.325146 |
| Up | RGPD8 | 28 | 0.002967 | 626564 | 0.32295 |
| Up | HSD3B1 | 28 | 0.009418 | 6545696 | 0.230497 |
| Up | COG3 | 27 | 0.00261 | 458374 | 0.24988 |
| Up | ATP6V0C | 27 | 0.008472 | 953284 | 0.320959 |
| Up | TFRC | 27 | 0.004398 | 847110 | 0.329004 |
| Up | TIAL1 | 27 | 0.005954 | 1083682 | 0.323485 |
| Up | DCP2 | 27 | 0.007451 | 4215608 | 0.243872 |
| Up | CAPZA1 | 26 | 0.002869 | 626878 | 0.325191 |
| Up | FKBP2 | 25 | 0.003414 | 683462 | 0.282255 |
| Up | TWIST1 | 24 | 0.001372 | 488874 | 0.298643 |
| Up | TIMP2 | 24 | 0.001662 | 741652 | 0.250924 |
| Up | SPTLC1 | 24 | 0.008272 | 878220 | 0.320827 |
| Up | DAPK1 | 24 | 0.003354 | 582304 | 0.3437 |
| Up | STT3B | 23 | 0.005888 | 688678 | 0.320805 |
| Up | RBMS1 | 23 | 0.005223 | 803708 | 0.322772 |
| Up | ANK3 | 21 | 0.004965 | 565126 | 0.321819 |
| Up | TDG | 21 | 0.00124 | 515588 | 0.326346 |
| Up | CHMP5 | 20 | 0.00503 | 681262 | 0.320915 |
| Up | MBD2 | 20 | 0.003168 | 3223460 | 0.265195 |
| Up | TRIM25 | 20 | 0.002013 | 463938 | 0.326073 |
| Up | TEAD3 | 20 | 0.001225 | 501618 | 0.323954 |
| Up | SLC3A2 | 20 | 0.005187 | 569152 | 0.332435 |
| Up | BMP1 | 19 | 0.00311 | 565522 | 0.259095 |
| Up | RHOBTB3 | 19 | 0.002152 | 346198 | 0.322439 |
| Up | CHD2 | 19 | 0.00121 | 998828 | 0.256894 |
| Up | RYBP | 19 | 0.002415 | 541940 | 0.322572 |
| Up | GLRX | 18 | 0.00459 | 583470 | 0.320959 |
| Up | KMT2C | 18 | 0.003676 | 553422 | 0.285019 |
| Up | YTHDC1 | 18 | 0.003349 | 530960 | 0.33407 |
| Up | GRB7 | 18 | 5.17E-04 | 151894 | 0.294958 |
| Up | IWS1 | 17 | 0.004258 | 1715034 | 0.236798 |
| Up | TRAF3IP2 | 17 | 0.001901 | 398464 | 0.324246 |
| Up | CSF3R | 17 | 0.001832 | 324530 | 0.329027 |
| Up | AFF1 | 17 | 0.003482 | 1512904 | 0.247805 |
| Up | INO80C | 17 | 0.003498 | 441732 | 0.321245 |
| Up | CASP4 | 17 | 0.002253 | 931980 | 0.263822 |
| Up | YIPF6 | 17 | 0.003441 | 5013996 | 0.201367 |
| Up | PCGF1 | 16 | 0.003259 | 409788 | 0.320827 |
| Up | CXCR2 | 16 | 0.003899 | 2017434 | 0.253558 |
| Up | ELF3 | 16 | 6.72E-04 | 203260 | 0.333452 |
| Up | USP33 | 16 | 0.002932 | 430028 | 0.322927 |
| Up | AGFG1 | 15 | 0.003227 | 435804 | 0.321443 |
| Up | LNPEP | 14 | 7.10E-04 | 211470 | 0.322483 |
| Up | RORA | 14 | 0.001493 | 299584 | 0.282187 |
| Up | LARP1B | 14 | 0.004683 | 546700 | 0.321708 |
| Up | ACSL4 | 14 | 0.005208 | 423594 | 0.320388 |
| Up | ESRRG | 14 | 0.001317 | 573474 | 0.244687 |
| Up | MXD1 | 13 | 6.21E-04 | 548286 | 0.273427 |
| Up | GET4 | 13 | 0.001988 | 297146 | 0.321113 |
| Up | NFE2 | 13 | 0.001758 | 361494 | 0.277475 |
| Up | SCNN1B | 13 | 0.001794 | 293352 | 0.321863 |
| Up | TBX3 | 11 | 0.00107 | 660104 | 0.254826 |
| Up | TAF1B | 11 | 0.00167 | 339720 | 0.323106 |
| Up | ADK | 11 | 0.003862 | 318638 | 0.320257 |
| Up | CLK3 | 11 | 0.001752 | 276054 | 0.272997 |
| Up | PELI1 | 10 | 0.001025 | 246998 | 0.322195 |
| Up | PHYHIPL | 2 | 1.23E-05 | 3410 | 0.238801 |
| Up | MAFF | 2 | 8.27E-06 | 5370 | 0.260986 |
| Up | TESK2 | 2 | 1.97E-05 | 2606 | 0.259425 |
| Up | MTMR4 | 1 | 0 | 0 | 0.259138 |
| Up | ZFAND6 | 1 | 0 | 0 | 0.230519 |
| Up | ACP1 | 1 | 0 | 0 | 0.27858 |
| Up | EFS | 1 | 0 | 0 | 0.27858 |
| Up | RAB11FIP5 | 1 | 0 | 0 | 0.220331 |
| Up | CYP11A1 | 1 | 0 | 0 | 0.187328 |
| Up | ADAM12 | 1 | 0 | 0 | 0.230951 |
| Up | FBXO9 | 1 | 0 | 0 | 0.25419 |
| Up | HSPB8 | 1 | 0 | 0 | 0.254674 |
| Up | CCNDBP1 | 1 | 0 | 0 | 0.256234 |
| Up | LTBP3 | 1 | 0 | 0 | 0.217731 |
| Up | FNTA | 1 | 0 | 0 | 0.246424 |
| Up | ACOX3 | 1 | 0 | 0 | 0.24266 |
| Up | PJA1 | 1 | 0 | 0 | 0.24372 |
| Up | SERTAD4 | 1 | 0 | 0 | 0.247308 |
| Up | EID2 | 1 | 0 | 0 | 0.259138 |
| Up | AKTIP | 1 | 0 | 0 | 0.254951 |
| Up | SPTLC3 | 1 | 0 | 0 | 0.242911 |
| Up | SGSM1 | 1 | 0 | 0 | 0.246424 |
| Up | LRRC69 | 1 | 0 | 0 | 0.292236 |
| Down | FYN | 305 | 0.060673 | 15146282 | 0.386124 |
| Down | ABL1 | 256 | 0.044808 | 14298188 | 0.385806 |
| Down | SMAD3 | 209 | 0.042609 | 22810928 | 0.349754 |
| Down | STAT3 | 202 | 0.037341 | 10226766 | 0.378422 |
| Down | PRKCA | 182 | 0.041662 | 11865040 | 0.377234 |
| Down | ACTA2 | 161 | 0.02569 | 7462512 | 0.366351 |
| Down | YWHAH | 149 | 0.028069 | 10474112 | 0.350015 |
| Down | ACTG2 | 131 | 0.011051 | 3779892 | 0.354141 |
| Down | RPL14 | 127 | 0.022838 | 13966136 | 0.331377 |
| Down | PDGFRB | 123 | 0.013902 | 4225738 | 0.358778 |
| Down | CCND1 | 119 | 0.017659 | 8841110 | 0.344483 |
| Down | PPM1F | 119 | 0.016265 | 6535610 | 0.32041 |
| Down | CAV1 | 113 | 0.018938 | 4593828 | 0.368628 |
| Down | YWHAQ | 101 | 0.013637 | 4845932 | 0.355674 |
| Down | FOXO1 | 96 | 0.015104 | 5652522 | 0.349128 |
| Down | WNT3A | 95 | 0.013629 | 4052094 | 0.299674 |
| Down | RHOQ | 92 | 0.016733 | 3678142 | 0.331471 |
| Down | TUBB2A | 91 | 0.012587 | 3886552 | 0.334213 |
| Down | FGFR3 | 83 | 0.010949 | 3642512 | 0.326755 |
| Down | RARA | 81 | 0.013223 | 4465300 | 0.341869 |
| Down | MYH11 | 72 | 0.01194 | 3062450 | 0.35249 |
| Down | HIPK2 | 72 | 0.01135 | 3565432 | 0.339639 |
| Down | HSPA12B | 68 | 0.003037 | 744544 | 0.301991 |
| Down | RPS6KA2 | 68 | 0.005549 | 2522498 | 0.317738 |
| Down | TEK | 66 | 0.004665 | 1653094 | 0.338168 |
| Down | NCOA1 | 65 | 0.008395 | 3153458 | 0.31475 |
| Down | VCL | 63 | 0.009681 | 2780132 | 0.338217 |
| Down | TUBB2B | 62 | 0.00676 | 1694194 | 0.343977 |
| Down | PPP2R2B | 62 | 0.010479 | 4191976 | 0.328542 |
| Down | TPM2 | 62 | 0.006776 | 1763750 | 0.334428 |
| Down | SPTAN1 | 62 | 0.011399 | 2596256 | 0.334046 |
| Down | MYL9 | 60 | 0.008098 | 2479906 | 0.335361 |
| Down | DIMT1 | 58 | 0.008481 | 2242228 | 0.327326 |
| Down | TCF4 | 54 | 0.009445 | 2760706 | 0.330629 |
| Down | MRPL34 | 52 | 0.006656 | 17810170 | 0.236106 |
| Down | CCND3 | 51 | 0.003239 | 1601134 | 0.295927 |
| Down | LMNA | 47 | 0.00911 | 2401846 | 0.338682 |
| Down | PKN1 | 46 | 0.003592 | 1349794 | 0.299559 |
| Down | CD44 | 46 | 0.010187 | 2270300 | 0.349493 |
| Down | DCN | 45 | 0.007594 | 1930616 | 0.322283 |
| Down | ADCY3 | 45 | 0.008618 | 1694556 | 0.33133 |
| Down | TGFB1I1 | 45 | 0.003137 | 931752 | 0.325349 |
| Down | EPHA3 | 44 | 0.002379 | 661600 | 0.318884 |
| Down | FZD4 | 43 | 0.005096 | 1738748 | 0.327006 |
| Down | RND3 | 42 | 0.006152 | 1123788 | 0.333832 |
| Down | ADCY4 | 41 | 0.001819 | 672196 | 0.287785 |
| Down | ETS2 | 41 | 0.004246 | 1834106 | 0.329559 |
| Down | RAP2A | 40 | 0.007351 | 1433240 | 0.326983 |
| Down | CCND2 | 40 | 0.002601 | 1155154 | 0.334046 |
| Down | AXIN2 | 39 | 0.00394 | 1203884 | 0.333618 |
| Down | OPTN | 37 | 0.00695 | 1707428 | 0.328289 |
| Down | UBE2L6 | 37 | 0.005601 | 1462120 | 0.327189 |
| Down | PIP5K1C | 37 | 0.002815 | 1263578 | 0.334714 |
| Down | MAP1B | 34 | 0.007029 | 1488054 | 0.325371 |
| Down | AXL | 34 | 0.002374 | 729250 | 0.345881 |
| Down | IRAK3 | 33 | 8.12E-04 | 865022 | 0.277229 |
| Down | FLNA | 33 | 0.004649 | 1087030 | 0.310146 |
| Down | COL1A1 | 32 | 0.008312 | 1395294 | 0.32942 |
| Down | DLL1 | 31 | 0.006383 | 1006300 | 0.324583 |
| Down | JAG1 | 29 | 0.005281 | 874336 | 0.325033 |
| Down | PDE9A | 29 | 0.003164 | 644058 | 0.281577 |
| Down | AGTR1 | 28 | 0.004101 | 926250 | 0.304088 |
| Down | TNNT3 | 27 | 1.64E-04 | 89708 | 0.268585 |
| Down | MYO10 | 27 | 0.002183 | 551478 | 0.325892 |
| Down | GPC4 | 26 | 0.001618 | 1207760 | 0.258652 |
| Down | IGFBP3 | 26 | 0.004163 | 894182 | 0.300288 |
| Down | ABCE1 | 25 | 0.007191 | 831802 | 0.321973 |
| Down | TIMP1 | 23 | 0.006574 | 950444 | 0.322905 |
| Down | VTN | 23 | 0.005502 | 970948 | 0.284138 |
| Down | ARAP3 | 23 | 8.48E-04 | 298776 | 0.326642 |
| Down | COL1A2 | 23 | 0.001462 | 373484 | 0.28554 |
| Down | B2M | 23 | 0.00758 | 619198 | 0.322505 |
| Down | TBCB | 22 | 0.002726 | 428028 | 0.322705 |
| Down | VASN | 22 | 0.003962 | 582444 | 0.343574 |
| Down | LGALS1 | 21 | 0.004371 | 578082 | 0.333903 |
| Down | CD81 | 21 | 0.005039 | 710796 | 0.323708 |
| Down | VCAM1 | 21 | 0.003333 | 2717522 | 0.2699 |
| Down | THY1 | 21 | 0.003795 | 675432 | 0.288956 |
| Down | PHGDH | 21 | 0.004117 | 689964 | 0.323999 |
| Down | ITPR3 | 21 | 0.003472 | 727920 | 0.327326 |
| Down | TGFB3 | 20 | 0.002631 | 725004 | 0.278316 |
| Down | BIN1 | 20 | 0.002083 | 351704 | 0.291835 |
| Down | LPAR1 | 20 | 0.001469 | 1296676 | 0.242258 |
| Down | MSX1 | 20 | 0.003633 | 648486 | 0.289134 |
| Down | SLIT2 | 20 | 0.001084 | 281232 | 0.291273 |
| Down | SHMT2 | 19 | 0.005637 | 547666 | 0.32215 |
| Down | COL4A1 | 19 | 0.001742 | 983934 | 0.257813 |
| Down | RAB3IL1 | 19 | 0.002314 | 1857444 | 0.231054 |
| Down | SPARC | 17 | 0.002023 | 197820 | 0.275923 |
| Down | VCAN | 17 | 0.003487 | 561304 | 0.267405 |
| Down | LDB2 | 17 | 0.003396 | 1291388 | 0.254812 |
| Down | DPYSL2 | 17 | 0.003321 | 506826 | 0.325869 |
| Down | MAF | 17 | 0.001088 | 288082 | 0.289456 |
| Down | PARVA | 16 | 0.003041 | 421786 | 0.323731 |
| Down | TNFRSF1B | 16 | 0.001419 | 325188 | 0.32382 |
| Down | FBLN2 | 16 | 0.001835 | 589690 | 0.249987 |
| Down | EEF2K | 15 | 0.001411 | 364288 | 0.323083 |
| Down | TEAD2 | 15 | 7.99E-04 | 352390 | 0.324066 |
| Down | TAGLN | 15 | 0.001241 | 205764 | 0.280381 |
| Down | FASN | 15 | 0.001757 | 367956 | 0.322084 |
| Down | CD36 | 15 | 0.002116 | 386412 | 0.294661 |
| Down | NDN | 15 | 0.003831 | 473694 | 0.322239 |
| Down | ADAM15 | 15 | 0.001982 | 294010 | 0.331283 |
| Down | ARHGEF2 | 15 | 9.41E-04 | 262426 | 0.330396 |
| Down | PEMT | 14 | 0.005116 | 629532 | 0.293664 |
| Down | VWF | 14 | 0.001828 | 345586 | 0.275097 |
| Down | OXTR | 14 | 0.001359 | 1221820 | 0.23325 |
| Down | FRYL | 14 | 0.001994 | 332012 | 0.321598 |
| Down | AGRN | 14 | 0.001733 | 263834 | 0.326482 |
| Down | LFNG | 13 | 0.001654 | 331212 | 0.264357 |
| Down | ARHGEF6 | 13 | 0.001032 | 179796 | 0.326573 |
| Down | TSC22D3 | 13 | 1.34E-04 | 119696 | 0.32391 |
| Down | IMPDH1 | 13 | 0.002712 | 390200 | 0.321554 |
| Down | NME4 | 12 | 0.003215 | 901430 | 0.228485 |
| Down | EHD2 | 12 | 0.003128 | 309602 | 0.320717 |
| Down | RGL1 | 12 | 5.53E-04 | 670384 | 0.247308 |
| Down | COL18A1 | 12 | 0.001098 | 309928 | 0.266008 |
| Down | ADGRA2 | 12 | 8.13E-04 | 196338 | 0.301583 |
| Down | FMOD | 11 | 2.68E-04 | 98450 | 0.300385 |
| Down | SLIT3 | 10 | 6.95E-05 | 22762 | 0.24207 |
| Down | COL5A1 | 3 | 4.31E-05 | 14682 | 0.244636 |
| Down | TGFBI | 3 | 8.62E-06 | 14986 | 0.248607 |
| Down | RASIP1 | 2 | 1.64E-06 | 1660 | 0.251732 |
| Down | CAV2 | 2 | 0 | 0 | 0.285836 |
| Down | PCOLCE | 2 | 3.25E-05 | 6584 | 0.246761 |
| Down | COL4A2 | 2 | 0 | 0 | 0.208517 |
| Down | SVIL | 2 | 0 | 0 | 0.262801 |
| Down | SMTN | 2 | 7.00E-06 | 2536 | 0.261073 |
| Down | COL4A5 | 2 | 7.45E-06 | 6672 | 0.244776 |
| Down | FSCN1 | 2 | 2.41E-06 | 2090 | 0.268585 |
| Down | ADAMTS1 | 2 | 5.19E-05 | 3268 | 0.250857 |
| Down | CTSC | 1 | 0 | 0 | 0.244559 |
| Down | FEZ1 | 1 | 0 | 0 | 0.21526 |
| Down | MALL | 1 | 0 | 0 | 0.269357 |
| Down | LYL1 | 1 | 0 | 0 | 0.248489 |
| Down | SEMA3A | 1 | 0 | 0 | 0.27858 |
| Down | ENPP2 | 1 | 0 | 0 | 0.195022 |
| Down | BST2 | 1 | 0 | 0 | 0.24757 |
| Down | F3 | 1 | 0 | 0 | 0.236738 |
| Down | COL16A1 | 1 | 0 | 0 | 0.204978 |
| Down | ANGPTL2 | 1 | 0 | 0 | 0.252724 |
| Down | NCALD | 1 | 0 | 0 | 0.250508 |
| Down | RASL12 | 1 | 0 | 0 | 0.228262 |
| Down | PPP1R14A | 1 | 0 | 0 | 0.273923 |
| Down | FCGRT | 1 | 0 | 0 | 0.243872 |
| Down | CNN2 | 1 | 0 | 0 | 0.268139 |
| Down | EHBP1 | 1 | 0 | 0 | 0.242848 |
| Down | COL3A1 | 1 | 0 | 0 | 0.216264 |
| Down | DDIT4 | 1 | 0 | 0 | 0.25408 |
| Down | ITM2C | 1 | 0 | 0 | 0.249441 |
| Down | GAS6 | 1 | 0 | 0 | 0.257007 |
| Down | FERMT2 | 1 | 0 | 0 | 0.252751 |
| Down | RAPGEF5 | 1 | 0 | 0 | 0.246424 |
| Down | APCDD1 | 1 | 0 | 0 | 0.230588 |
| Down | MMP23B | 1 | 0 | 0 | 0.200599 |
| Down | PTPRD | 1 | 0 | 0 | 0.274549 |
| Down | RSPO3 | 1 | 0 | 0 | 0.249441 |
| Down | COL6A1 | 1 | 0 | 0 | 0.243745 |
| Down | DDR2 | 1 | 0 | 0 | 0.247805 |
| Down | NEK6 | 1 | 0 | 0 | 0.244776 |
| Down | COL5A2 | 1 | 0 | 0 | 0.205788 |
| Down | TSPAN4 | 1 | 0 | 0 | 0.244559 |
| Down | IFITM3 | 1 | 0 | 0 | 0.27858 |
| Down | HES4 | 1 | 0 | 0 | 0.261875 |
| Down | FAT1 | 1 | 0 | 0 | 0.242521 |

**Table 6.**
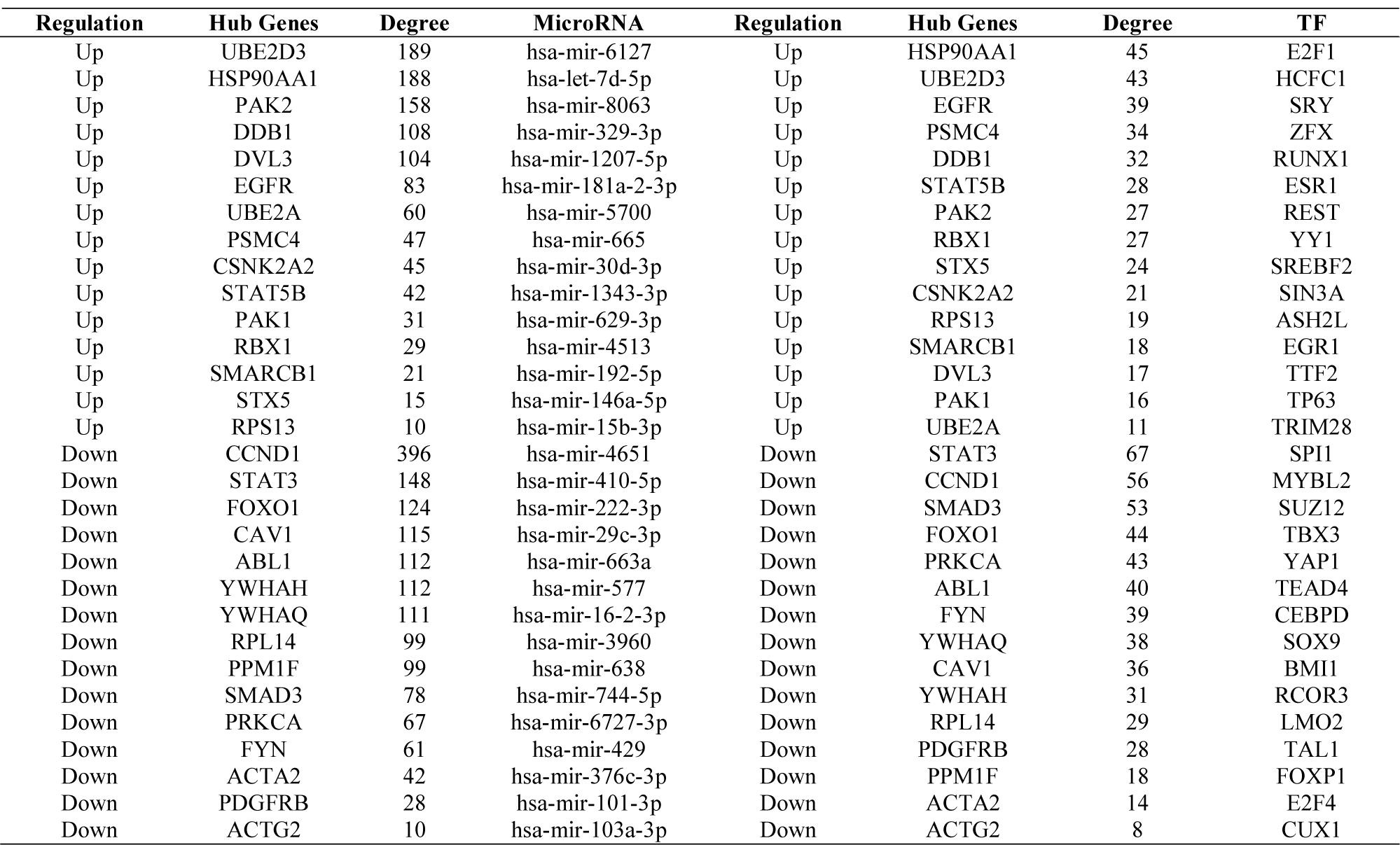
miRNA - hub gene and TF – hub gene interaction

### TF-hub gene regulatory network construction

NetworkAnalyst database was applied to screen the targeted TFs of the hub genes. Cytoscape software was used to construct the TF-hub gene network. As illustrated in Fig. 6, the interaction network consists of 306 hub genes and 195 TFs. According to the hub genes and TFs in the network ranked by their degree of connectivity using Network Analyzer and are listed in Table 4. Based on the expression trend of hub genes in GDM, we found that HSP90AA1 was the predicted target of E2F1, UBE2D3 was the predicted target of HCFC1, EGFR was the predicted target of SRY, PSMC4 was the predicted target of ZFX, DDB1 was the predicted target of RUNX1, STAT3 was the predicted target of SPI1, CCND1 was the predicted target of MYBL2, SMAD3 was the predicted target of SUZ12, FOXO1 was the predicted target of TBX3 and PRKCA was the predicted target of YAP1.

**Fig. 6.**
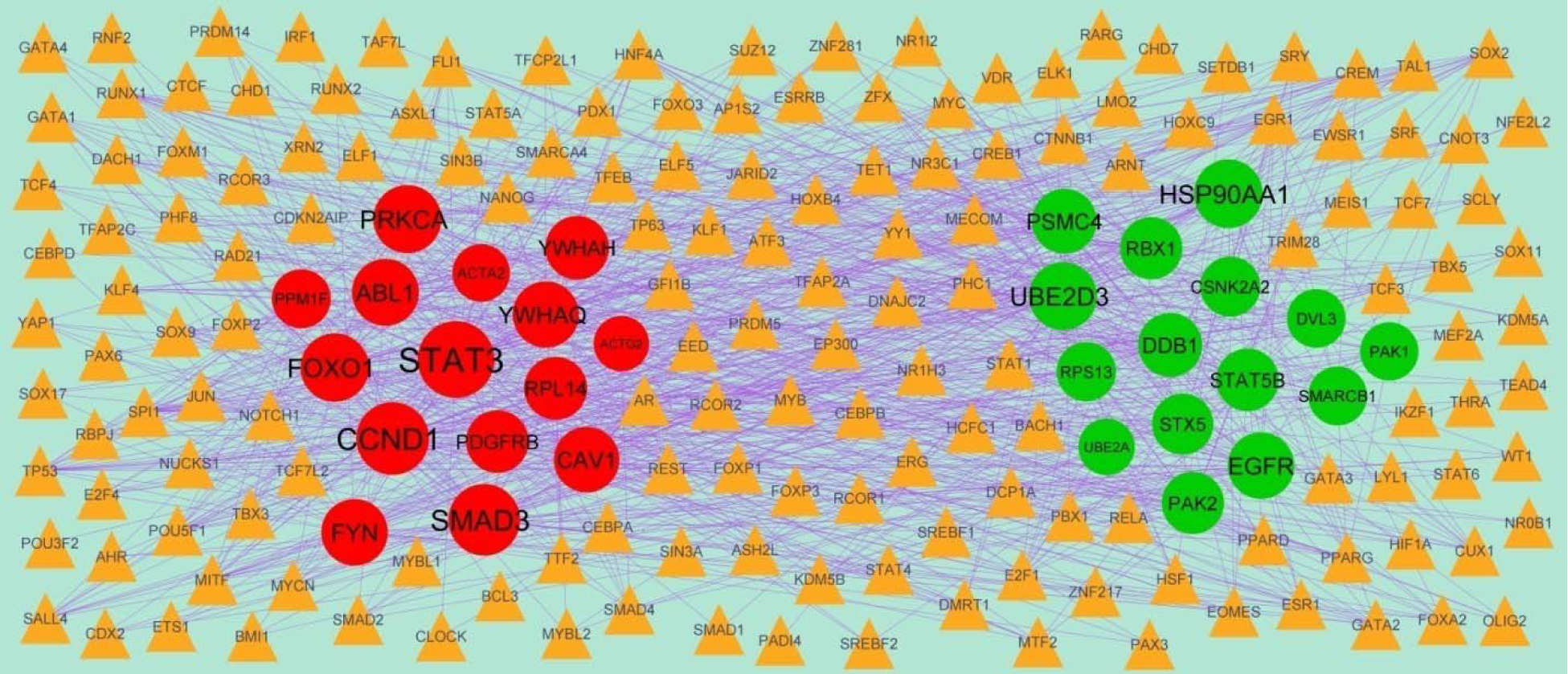
TF - hub gene regulatory network. The yellow color triangle nodes represent the key TFs; up regulated genes are marked in green; down regulated genes are marked in red.

### Receiver operating characteristic (ROC) curve analysis

ROC curve analysis was implemented to evaluate the capacity of hub genes to distinguish GDM and non GDM in E-MTAB-6418, HSP90AA1, EGFR, RPS13, RBX1, PAK1, FYN, ABL1, SMAD3, STAT3 and PRKCA, exhibiting better diagnostic efficiency for GDM and non GDM, and the combined diagnosis of these ten hub genes was more effective. The AUC index for the 10 hub gene scores were 0.906, 0.838, 0.825, 0.897, 0.863, 0.876, 0.855, 0.880, 0.932 and 0.872, and are shown Fig.7.

**Fig. 7.**
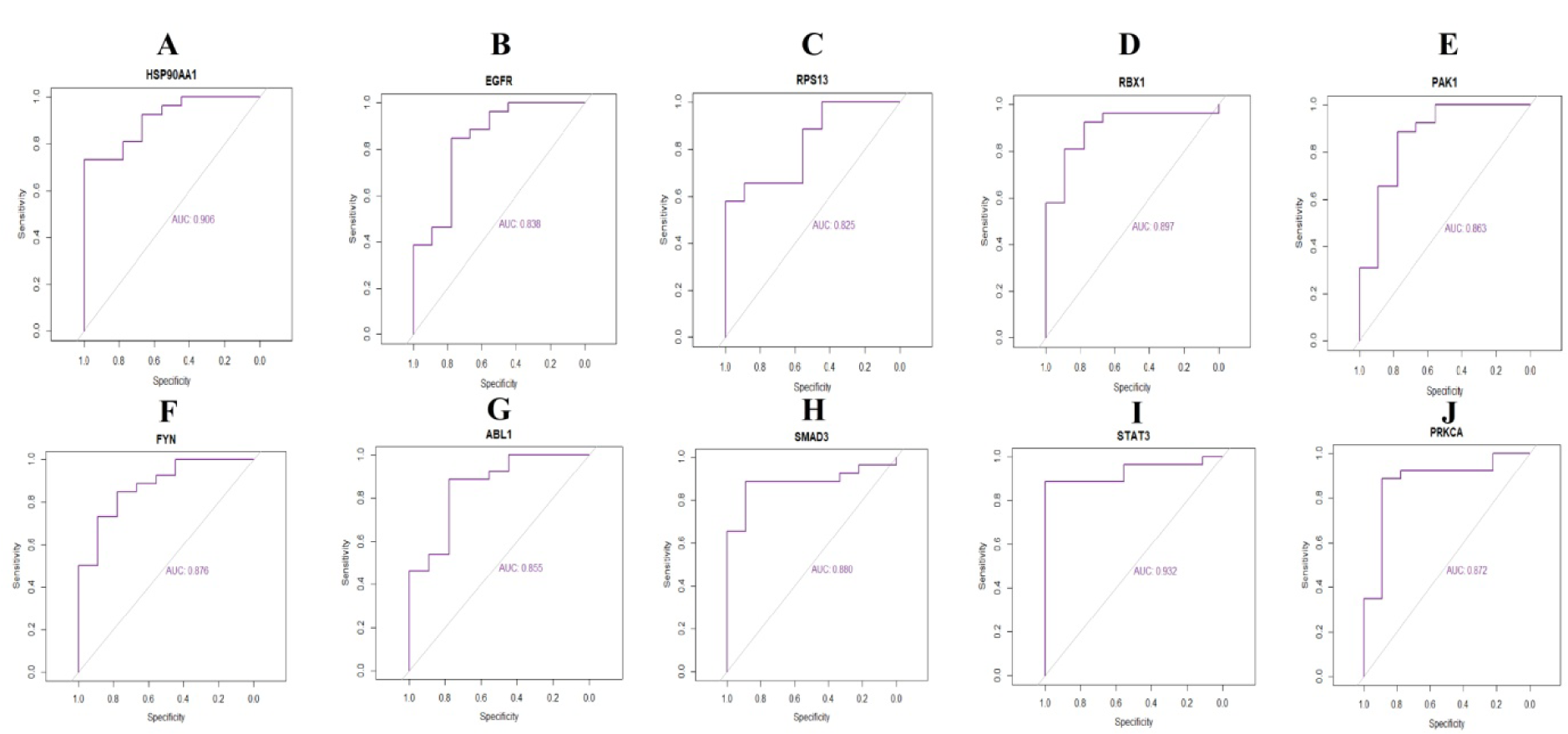
ROC curve validated the sensitivity, specificity of hub genes as a predictive biomarker for GDM prognosis. A) HSP90AA1 B) EGFR C) RPS13 D) RBX1 E) PAK1 F) FYN G) ABL1 H) SMAD3 I) STAT3 J) PRKCA

### RT-PCR Analysis

To further verify the expression level of hub genes in GDM, RT-PCR was performed to calculate the mRNA levels of the ten hub genes identified in the present study (HSP90AA1, EGFR, RPS13, RBX1, PAK1, FYN, ABL1, SMAD3, STAT3 and PRKCA) in GDM. As illustrated in Fig. 8, the expression of HSP90AA1, EGFR, RPS13, RBX1, PAK1 were significantly up regulated in GDM samples compared with normal, while FYN, ABL1, SMAD3, STAT3 and PRKCA were significantly down regulated in GDM samples compared with normal. The present RT-PCR results were in line with the aforementioned bioinformatics analysis, suggesting that these hub genes might be linked to the molecular mechanism underlying GDM.

**Fig. 8.**
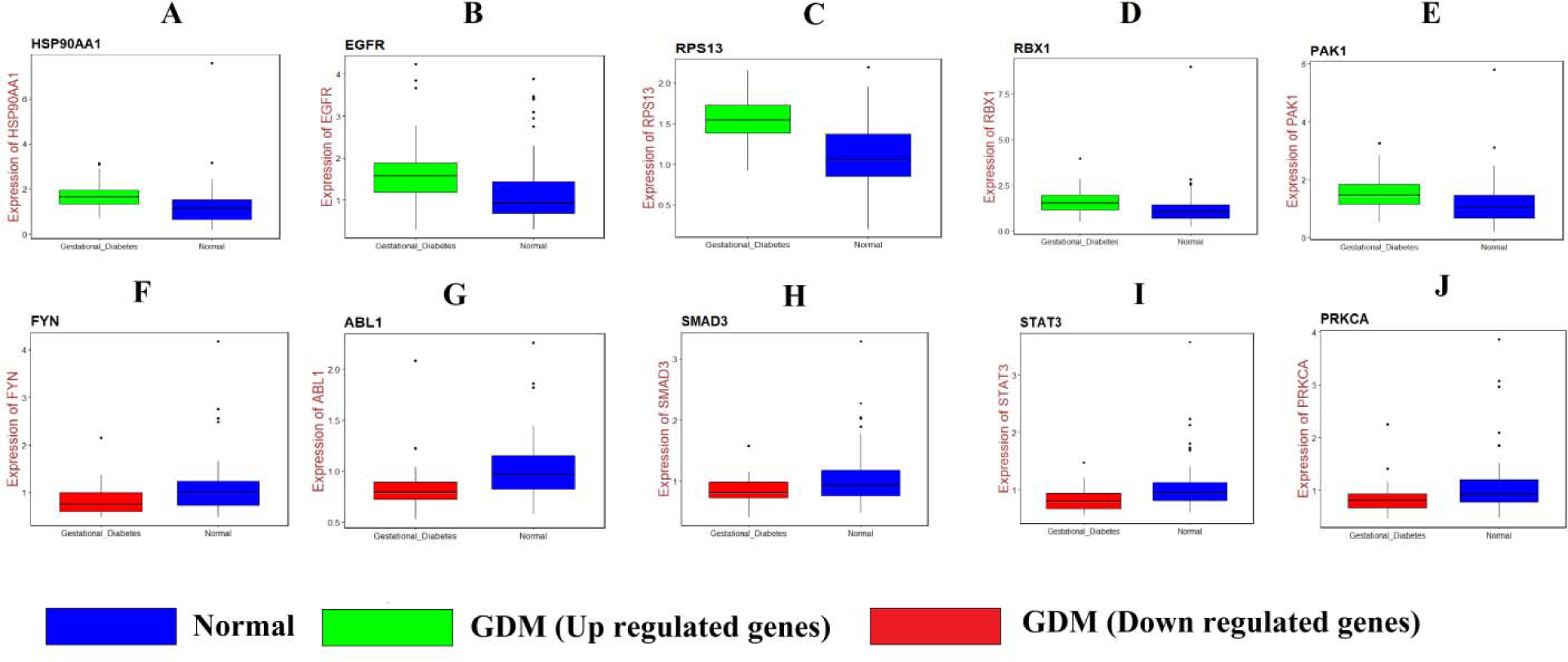
Validation of hub genes by RT-PCR. A) HSP90AA1 B) EGFR C) RPS13 D) RBX1 E) PAK1 F) FYN G) ABL1 H) SMAD3 I) STAT3 J) PRKCA

### Molecular docking experiments

In the recent findings, the docking study was performed using Biovia Discovery Studio perpetual software to analyse the binding pattern of the natural plants products such as herbs have the ability to lower blood glucose levels and ameliorate diabetes with decreased adverse side effects. The natural well known phytoconstituents which decreases the blood sugar level are Malvidin 3-laminaribioside (MLR), Ferulic acid (FRA), Inosporone (INO), Allicin (ALL), Liriodenin (LIR), Azadirachitin (AZA), Sulforaphane, Cajanin (CAJ), Carvone (CAR), Capsaicin (CAP), Terpineol (TER), Phellandrene (PHE), Terpene (TPN), Ellagic acid (ELA), Leucodelphinidin, O-methyltylophorinidine (OMT), Gymnemic acid, beta-Carotene (BCR), Leucocyanidin (LEC), Syringin (SYR), Ginsenoside (GNS), Phyllanthin (PHY), Punicalagin (PUC), Punicalin (PUN), Arjunic acid (AJA), Arjunetin (ARJ), Arabic acid (ARA), Arjungenin (ARG), Gingerol (GIN), Shogaol, Aloe emodin (ALE), Arabic acid (ARA), Aloin (ALO), Charantin (CHR), Cinnamic acid (CIN), Curcumin (CUR), Euginol (EUG), Gymnemagenin (GMG), Gymnestrogenin (GYM), Hydroxylucin (HYD), Methoxy hydroxyl chalcoli (MHC), Myricetin (MYR), Nimbine (NIM), Quercetin (QUE), Vicine (VIC) and Shagoal (SHA) are shown in Fig. 9. The molecules were constructed based on the natural plant products containing these chemical constituents which play vital role in reducing type 2 diabetes mellitus. The traditional plant products are used in conjunction with allopathic drug to reduce the dose of the allopathic drugs and or to increase the efficacy of allopathic drugs. Some common and most prominent antidiabetic plants and active principles were selected from their phytochemicals for docking studies in the present research to identify the active natural molecule to avoid the use of allopathic drugs in gestational diabetes and the blood sugar level is controlled by altering the diet. For docking experiments well known and most commonly used two allopathic drugs such as Glyburide (GLY), Metformin (MET) in gestational diabetes are used as standard and to compare the binding interaction of natural phyto constituents with allopathic drugs. A total of common 44 in that 42 natural active constituents few from each of flavonoids, saponins, tannins and glycosides etc., present in plant extracts responsible for antidiabetic function and 2 allopathic drugs were chosen for docking studies on over expressed proteins and the structures are depicted in figure 1 respectively. The one protein from each over expressed genes in gestational diabetes 2 diabetes mellitus such as EGFR (epidermal growth factor receptor), HSP90AA1 (heat shock protein 90 alpha family class A member 1), PAK1 (p21 (RAC1) activated kinase 1), and RBX1 (ring-box 1) and their X-RAY crystallographic structure and co-crystallized PDB code and their PDB code of 4UV7, 5NJX, 3Q4Z and 3FNI respectively were constructed for docking. The docking on natural active constituents was conducted to classify the potential molecule and their binding affinity to proteins. A higher number of negative number -CDOCKER energy and binding energy indicates a stronger binding interactions with proteins, few constituents obtained with a greater -CDOCKER energy and binding energy respectively with particular proteins. Docking experiments were carried out on a total of 42 constituents from plant products, few constituents obtained excellent -CDOCKER energy and binding energy. Out of 44 molecules few of the molecules obtained -CDOCKER interaction energy of more than 40 and majority with more than 30 and less than 40, few molecules obtained optimum -CDOCKER interaction energy of less than 30 respectively. the molecules with -CDOCKER interaction energy of 40 and above are said to have good interaction with proteins and stable. The natural constituents of the molecules GLY, GNS, GYM, MLR, PUC and ALO, GLY, MLRand ALE, ALO, BCR, CAP, CHR, ELA, LUR, GIN, GLY, GMG, GNS, GYM, LEC, LIR, MLR, MYR, NIM, OMP, PHY, PUC, PUN, QUE, SHE,VI C obtained a -CDOCKER interaction energy of more than 40 with protein of PDB code 5NJX and 3FNI and 3Q4Z respectively. The natural constituents obtained -CDOCKER interaction energy of less than 40 and more than 30 are ALO, ARJ, BCR, CHR, CUR, PHY, PUN and BCR, CAJ, CAP, CUR, GIN, LEC, MYR, OMP, QUE, VIC and AJA, ARA, ARG, CAJ, FRA, HYD, MHC and GNS, PHY, PUC, PUN with 5NJX and 3FNI and 3Q4Z and 4UV7. The constituents obtained less than 30 and more than 20 are AJA, ALE, ARG, CAJ, CAP, GIN, GMG, GYM, HYD, LEC, MHC, MLR, MYR, NIM, OMP, QUE, VIC and AJA, ALE, ALL, ARG, AJA,CHR, CIN, EUG, FRA, GMG, GNS, GYM, LIR, MHC, NIM, PUC and ALL, CIN, EUG, MET, TER and ALA, ALE, ALO, ARJ, BCR, CAJ, CHR, ELA, FRA, GIN, GMG, LEC, MLR, MYR, OMP, QUE, SHA with 5NJX and 3FNI and 3Q4Z and 4UV7. Following the molecules obtained less than 20 -CDOCKER interaction energy are ALL, ARA, CAR, CHR, CIN, EUG, FRA, LIR, MET, PHE, TER, TPN and ARJ, ARA, CAR, HYD, MET, PHE, TPN and CAR, PHE, TPN and AJA, ALL, ARG, CAR, CIN, EUG, GYM, HYD, LIR, MET, MHC, NIM, PHE, TER, TPN, VIC with protein 5NJX and 3FNI and 3Q4Z and 4UV7 respectively the biding energy,-CDOCKER energy and -CDOCKER interaction energy are depicted in Table 7. The two molecules such as ALO and MAL Fig. 10 and Fig. 11, their interaction with amino acids of proteins with 3D strictures for 3FN1 Fig.12 and 3Q4Z Fig.13, while 2D strictures for 3FN1 Fig.14 and 3Q4Z Fig.15.

**Fig 9.**
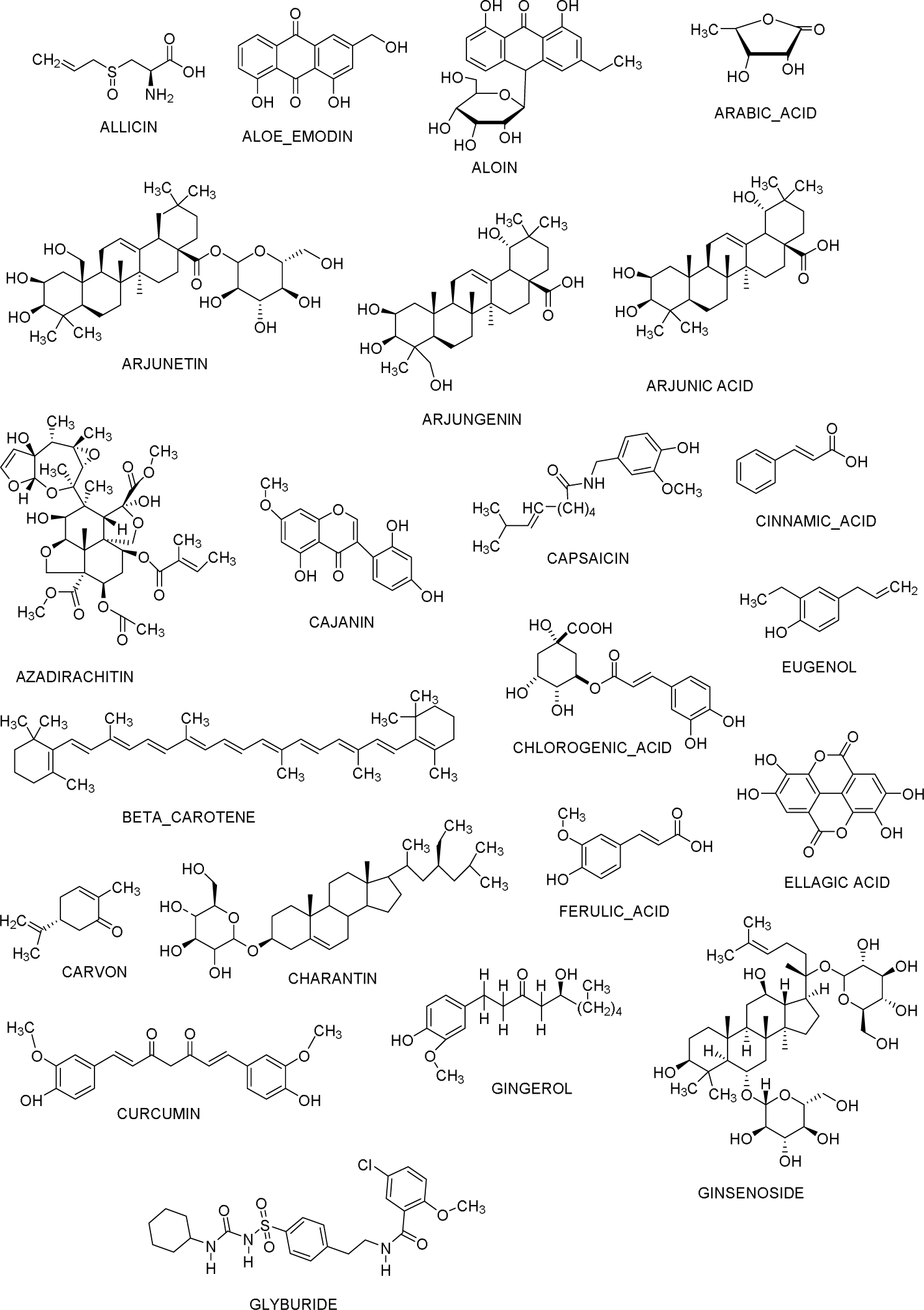
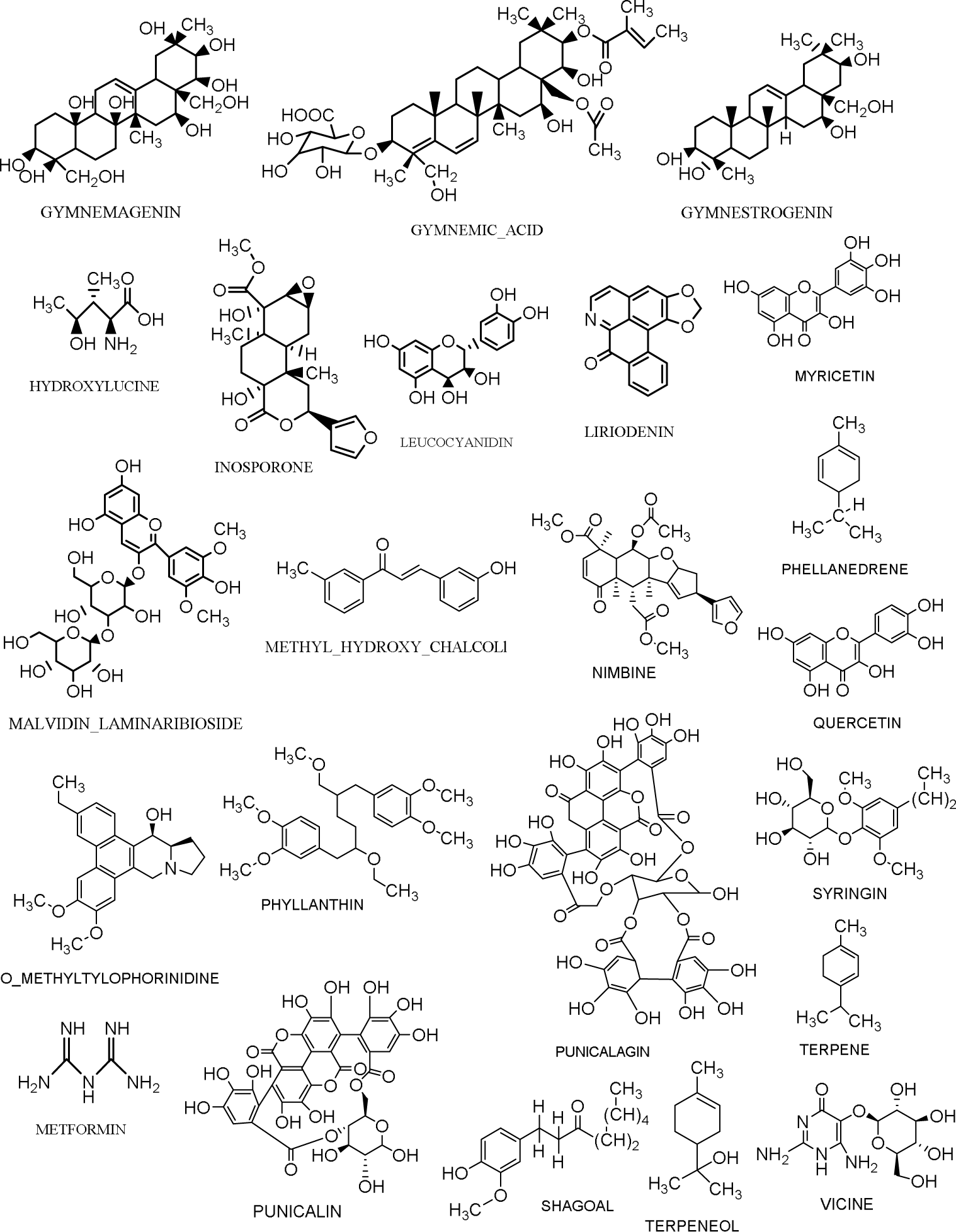
Chemical Structures of Phytoconstituents

**Fig. 10.**
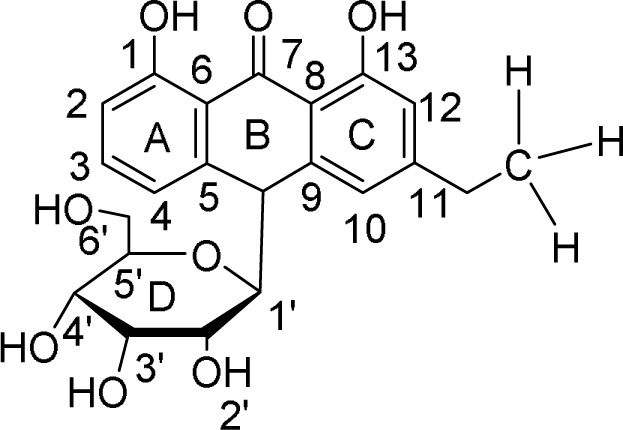
Structure of ALO

**Fig. 11.**
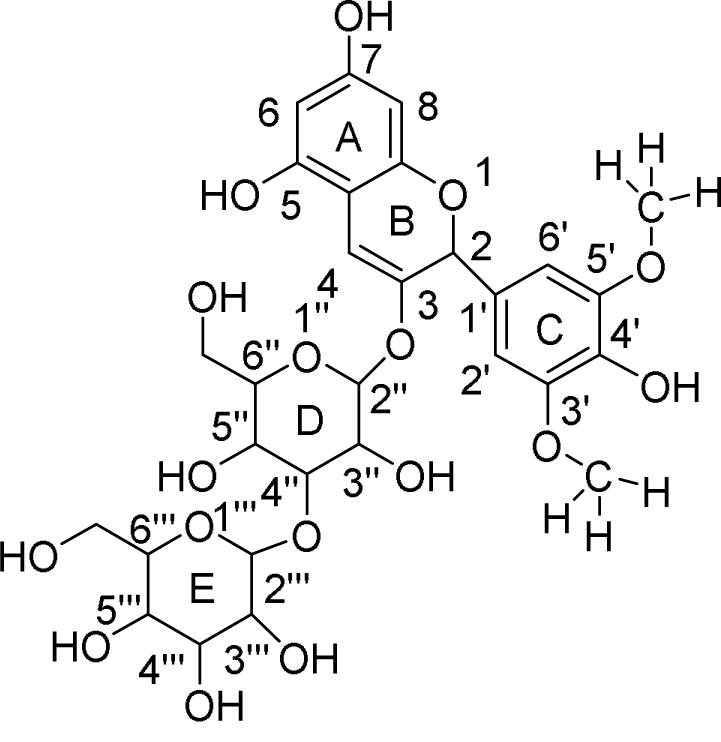
Structure of MAL

**Fig. 12.**
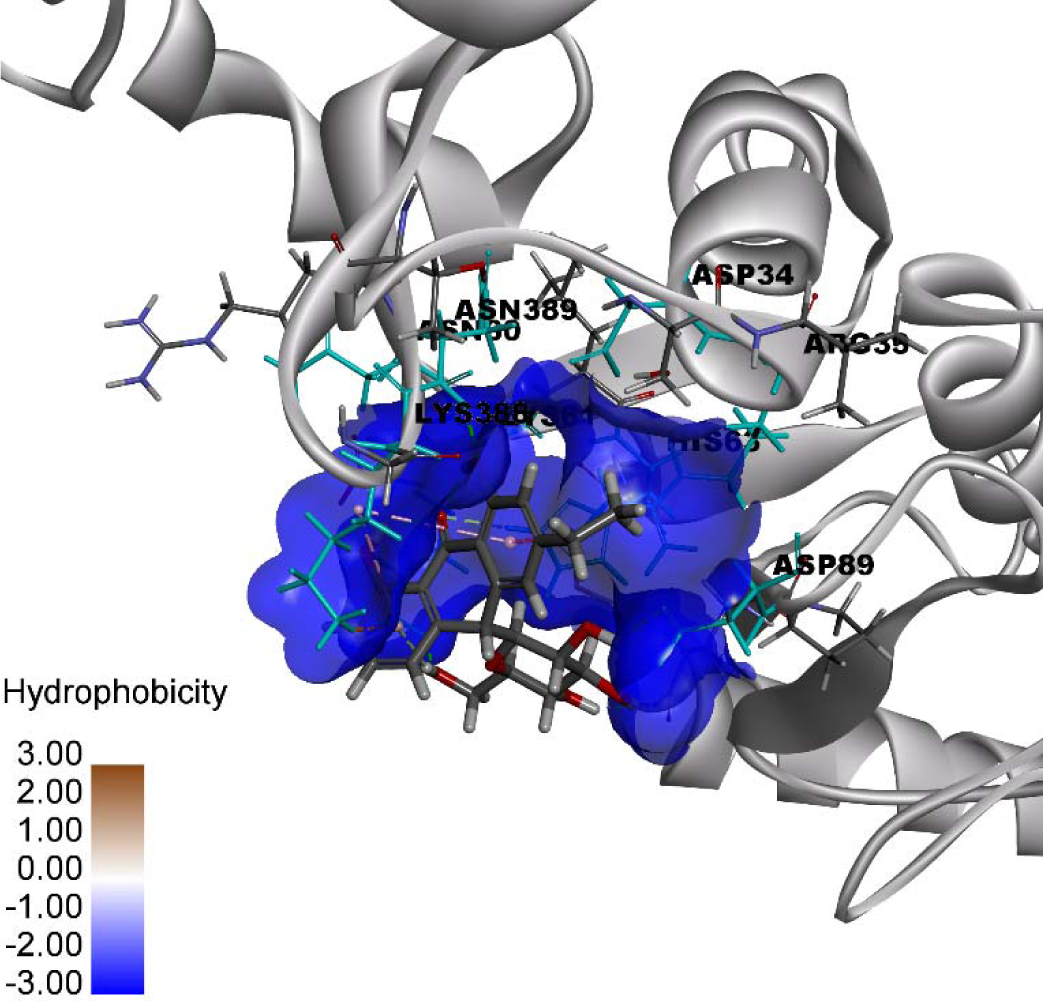
3D Binding of ALO with 3FN1

**Fig. 13.**
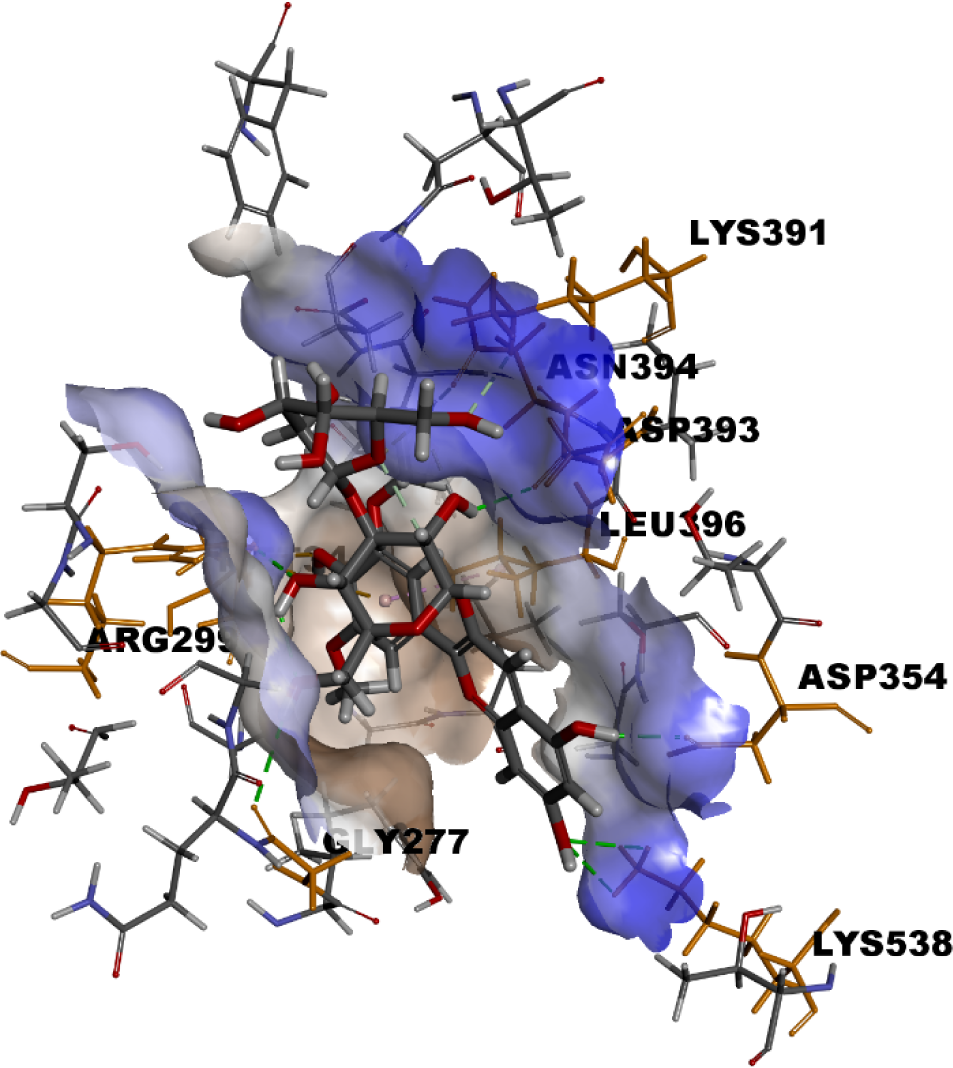
3D Binding of MAL with 3Q4Z

**Fig. 14.**
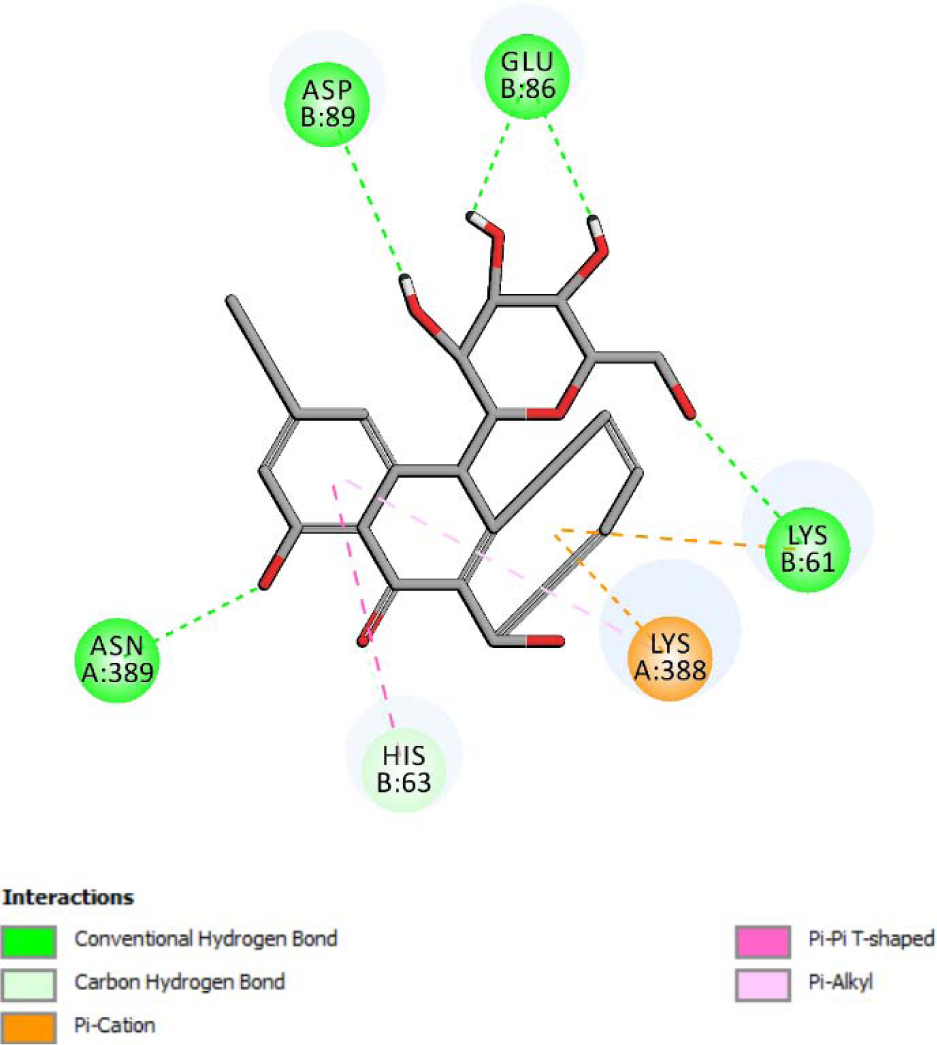
2D Binding of ALO with 3FN1

**Fig. 15.**
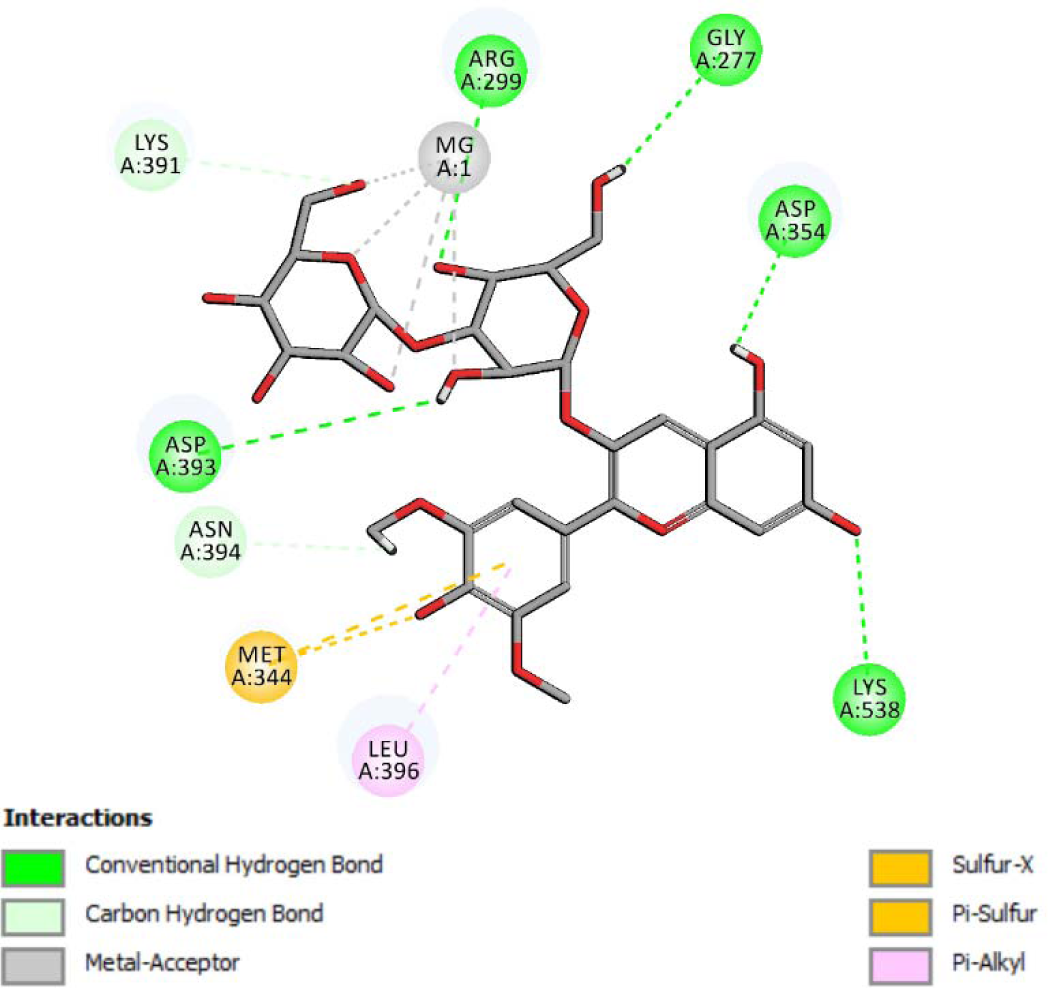
2D Binding of MAL with 3Q4Z

**Table 7.**
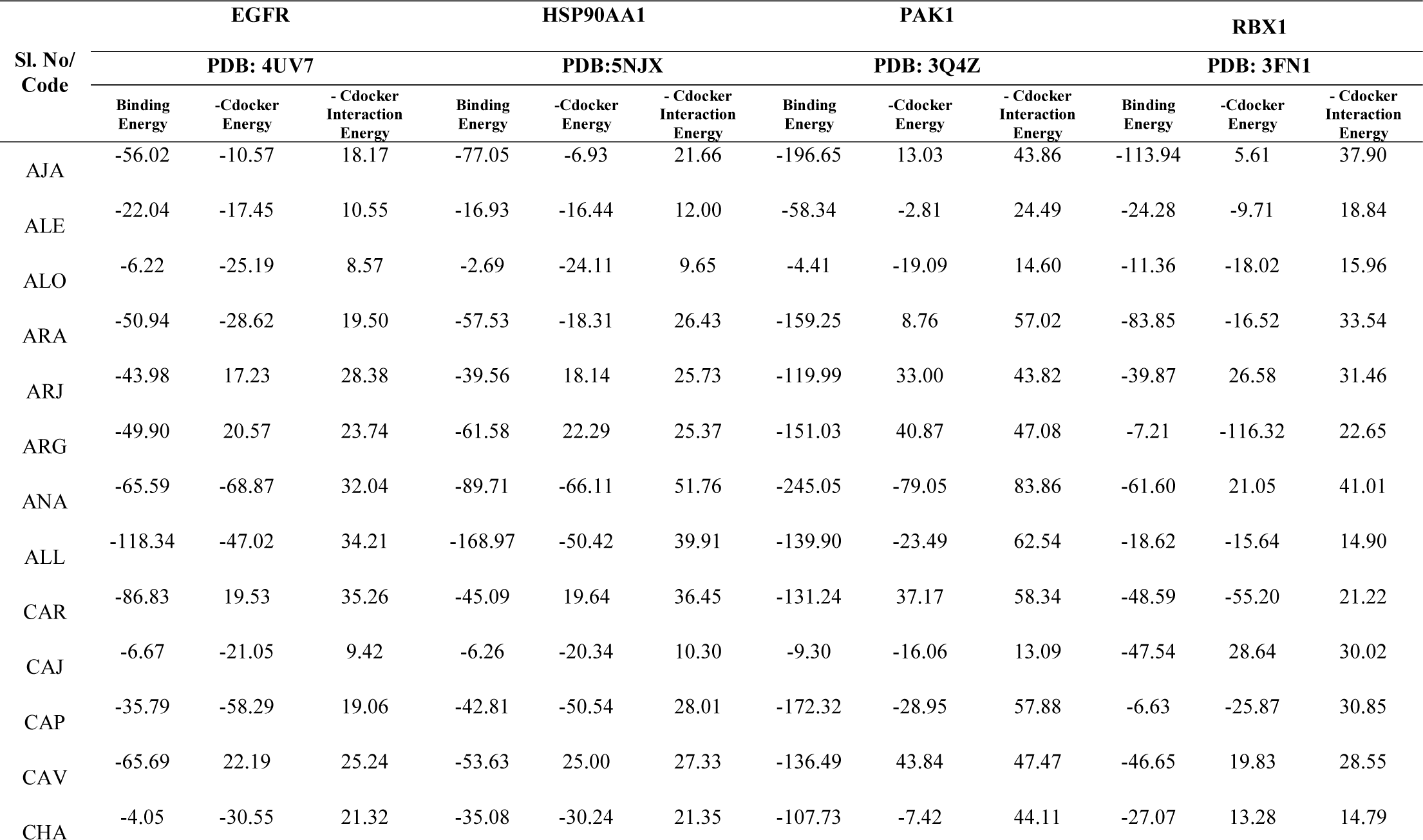
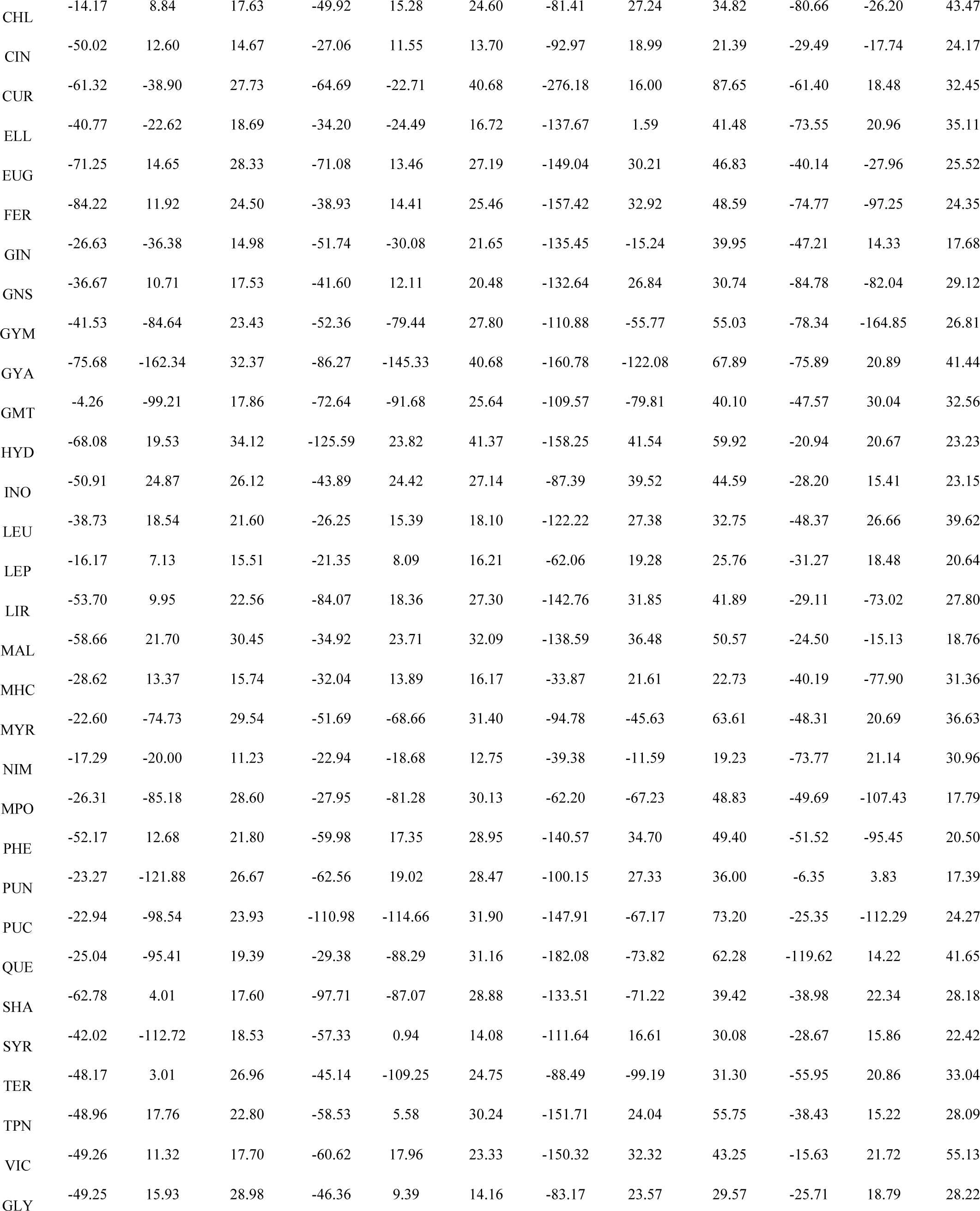
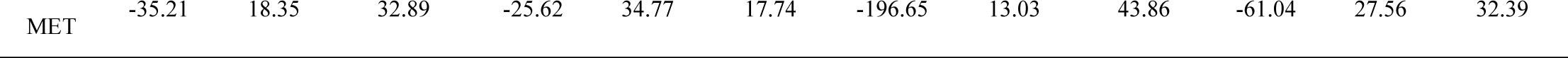
Docking results of designed molecules on over expressed proteins

## Discussion

Although people have continuously investigated GDM, the early diagnosis and treatment of GDM is still a huge problem due to the inadequacy of understanding of the molecular mechanisms that drive the occurrence and progression of GDM.

Therefore, in-depth investigation into the factors and mechanisms of GDM advancement are necessary for GDM diagnosis and treatment. Due to well-developed transcription profiling by array technology, it is accessible to resolve the general genetic modification in the development of diseases, which can allow for the recognition of gene targets for diagnosis, therapy, and prognosis of GDM.

In our study, a total of 869 DEGs were screened, including 439 up regulated genes and 430 down regulated genes. Several studies have reported that expression of CGB5 was essential for pregnancy success [30]. Aberrations of CRH (corticotropin releasing hormone) [31] and PSG1 [32] contribute to preeclampsia occurrence. The expression of CYP19A1 was significantly up regulated in hypertensive disorders of pregnancy [33]. Based on previous studies, CD248 is generally associated with progression of hypertension [34], but this gene might be linked with development of GDM. Lin et al [35] reported that expression of COL1A1 was essential for type 2 diabetes mellitus progression, but this gene might be involved in the development of GDM. Delfín et al [36] found that ABI3BP was responsible for progression of cardiovascular diseases, but this gene might be linked with development of GDM. MFAP4 was reported to cause type 1diabetes mellitus [37], but this gene might be responsible for progression of GDM.

DEGs were found to be enriched in reproduction, nuclear outer membrane-endoplasmic reticulum membrane network, identical protein binding, cell surface interactions at the vascular wall, cell adhesion, supramolecular complex, signaling receptor binding and extracellular matrix organization. CEBPB (CCAAT enhancer binding protein beta) [38], ACSL4 [39], MBD2 [40], ULK1 [41], NUCB2 [42], TWIST1 [43], HOOK2 [44], CLDN7 [45], TBK1 [46], YIPF6 [47], TFRC (transferrin receptor) [48], ENPP2 [49], SLIT2 [50], MFGE8 [51], FAT1 [52], GPC4 [53], COL6A3 [54], EGFL6 [55], AOC3 [56], CCN2 [57], LYVE1 [58], RARA (retinoic acid receptor alpha) [59], COL18A1 [60], THY1 [61], CD36 [62], PEMT (phosphatidylethanolamine N-methyltransferase) [63], AIF1L [64], OXTR (oxytocin receptor) [65], LMNA (lamin A/C) [66], CXCL14 [67], DKK3 [68], ANGPTL2 [69] and CMTM7 [70] were reported to be associated with obesity, but these genes might be linked with progression of GDM. AHR (aryl hydrocarbon receptor) [71], STS (steroid sulfatase) [72], PLAC1 [73], CYP11A1 [74], PSG11 [75], STAT5B [76], TLR3 [77], FOLR1 [78], HSPB1 [79], HSP90AA1 [80], ANXA4 [81], ATF3 [82], DAPK1 [83], ENTPD1 [84], ABL1 [85], VSIG4 [86], CD99 [87], VWF (von Willebrand factor) [88], PODXL (podocalyxin like) [89], PDPN (podoplanin) [90], RND3 [91], VCAN (versican) [92], AXL (AXL receptor tyrosine kinase) [93], PIEZO1 [94], GAS6 [93], LAMA4 [95], CAV1 [96], DLL1 [97], CD44 [98], CD81 [99], SMAD3 [100], NES (nestin) [101], DCN (decorin) [102], AGTR1 [103], SLIT3 [104], B2M [105], STAT3 [106], STC1 [107] and ADAMTS1 [108] were shown to participate in facilitating the preeclampsia. Majchrzak-Celiń ka et al [109] and Shimodaira et al [110] reported that HSD11B2 and HSD3B1 are responsible for hypertensive disorders of pregnancy. Altered expression of CSNK2A2 [111], NFE2 [112], CAMK2G [113], RASGRP1 [114], S100P [115], SRR (serine racemase) [116], DHPS (deoxyhypusine synthase) [117], DYRK1A [118], JAG1 [119], COL3A1 [120], VTN (vitronectin) [121], WNT3A [122], ACTA2 [123], SEMA3A [124], RARRES2 [125], CAV2 [126] and SPRED1 [127] were observed to be associated with the progression of type 2 diabetes mellitus, but these genes might be liable for advancement of GDM. In a previous report, Santiago et al [128], Auburger et al [129], Qu et al [130], Śnit et al [131] and Hjortebjerg et al [132] reported that SLC22A5, SH2B3, ITPR3, CALD1 and IGFBP4 expression might be regarded as an indicator of susceptibility to type 1 diabetes mellitus, but these genes might be associated with progression of GDM. Krishnan et al [133], Hu et al [134], Martins et al [135], Prieto-Sánchez et al [136], Sugulle et al [137], Zhao et al [138], Siddiqui et al [139], Han et al [140], Lappas et al [141], Wang et al [142], Artunc-Ulkumen et al [143], Blois et al [144], Vacínová et al [145] and Vilmi-Kerälä et al [146] demonstrated that the expression of CREBRF (CREB3 regulatory factor), STRA6, EGFR (epidermal growth factor receptor), MFSD2A, GDF15, PAK1, VCAM1, IGFBP2, IGFBP7, PRKCA (protein kinase C alpha), ADAMTS9, LGALS1, BIN1, TIMP1 and are associated with progression of GDM. Aquila et al, [147],Chen et al. [148], Xie et al. [149], Zhang et al. [150], Aspit et al. [151], Akadam-Teker et al. [152], Jiang et al. [153], Cetinkaya et al. [154], Grond-Ginsbach et al [155], Dong et al. [156], Chardon et al. [157], Chen et al. [158], Yamada et al. [159], Hu et al. [160], Bobik and Kalinina [161], Schwanekamp et al. [162], Liu et al. [163], Schroer et al. [164], Raza et al. [165], Yang et al. [166], Azuaje et al. [167], Durbin et al. [168], Chowdhury et al. [169], Wang et al. [170], Li et al. [171], Lv et al. [172], Bertoli-Avella et al. [173], Grossman et al. [174], Andenæs et al. [175] and Chen et al. [176] demonstrated that HES1, SPIN1, TBX3, EVA1A, CAP2, BMP1, HSPB8, RDX (radixin), COL5A1, LIMS2, PARVA (parvin alpha), EGFLAM (EGF like, fibronectin type III and laminin G domains), NEXN (nexilin F-actin binding protein), TNFRSF14, TGFBI (transforming growth factor beta induced), HAVCR2, CDH11, COL4A1, COL4A2, COL5A2, SHROOM3, HYAL2, PDLIM3, ETS2, PLSCR4, TGFB3, COL6A2 and LTBP2 could induce cardiovascular diseases, but these genes might be essential for progression of GDM. Flamant et al [177], Wan et al [178], Zhang et al [179], Vallvé et al [180], Heximer and Husain [181], Selvarajah et al [182], Jain et al [183], Sun et al [184], Satomi-Kobayashi et al [185], Jiang et al [186], Waghulde et al [187] and Dahal et al [188] reported that DDR1, CAST (calpastatin), KYNU (kynureninase), FBLN2, SPON1, VEGFC (vascular endothelial growth factor C), FLNA (filamin A), SNAI2, MYADM (myeloid associated differentiation marker), NECTIN2 and SMTN (smoothelin), GPER1, PDGFRB (platelet derived growth factor receptor beta) crucially contribute to the development of hypertension, but these genes might be linked with advancement of GDM.

From the PPI network and modules diagram, it can be observed that HSP90AA1, EGFR, RPS13, RBX1, PAK1, FYN (FYN proto-oncogene, Src family tyrosine kinase), ABL1, SMAD3, STAT3, PRKCA, UBE2A, UCHL3, TUBB2A, ACTA2 and TBCB (tubulin folding cofactor B) were the key nodes of the PPI network and modules, with the highest node degree, betweenness, stress and closeness value. RPS13, RBX1, FYN, UBE2A, TUBB2A and TBCB were the novel biomarkers for the progression of GDM.

From the miRNA-hub gene network construction and TF-hub gene network diagram, it can be observed that UBE2D3, HSP90AA1, PAK2, DDB1, DVL3, FYN, ABL1, SMAD3, STAT3, PRKCA, EGFR, PSMC4, CCND1, FOXO1, hsa-mir-6127, hsa-let-7d-5p, hsa-mir-8063, hsa-mir-329-3p, hsa-mir-1207-5p, hsa-mir-4651, hsa-mir-410-5p, hsa-mir-222-3p, hsa-mir-29c-3p, hsa-mir-663a, E2F1, HCFC1, SRY, ZFX, RUNX1, SPI1, MYBL2, SUZ12, TBX3 and YAP1 were the key nodes of the miRNA-hub gene network construction and TF-hub gene network, with the highest node degree value. Expression of the CCND1 gene plays a role in the development of obesity [189], but this gene might be associated with progression of GDM. FOXO1 [190], hsa-mir-1207-5p [191], hsa-mir-4651 [191], hsa-mir-222-3p [192] and E2F1 [193] are essential for the progression of GDM.

Hsa-let-7d-5p [194], hsa-mir-29c-3p [195] and SRY (Sex-determining Region Y) [196] have been shown to have an important role in type 2 diabetes mellitus, but these genes might be responsible for progression of GDM. Hsa-mir-663a [197] and TBX3 [198] have been shown as a promising biomarker in cardiovascular diseases, but this gene might be involved in progression of GDM. RUNX1 [199] and YAP1 [200] have been demonstrated to function in preeclampsia. UBE2D3, PAK2, DDB1, DVL3, PSMC4, hsa-mir-6127, hsa-mir-8063, hsa-mir-329-3p, hsa-mir-410-5p, HCFC1, ZFX (zinc finger protein, X-linked), SPI1, MYBL2 and SUZ12 were the novel biomarkers for the progression of GDM.

The molecule GLY, MLR obtained a good -CDOCKER interaction energy with 5NJX, 3FNI and 3Q4Z the -CDOCKER interaction energy of GLY is 41.37, 59.92, 41.44 and for MLR is 40.68, 87.65, 43.47 with 5NJX, 3FNI and 3Q4Z respectively. The two molecules such as ALO and MAL its interaction with amino acids are 2’ hydroxyl group formed hydrogen bond interaction with ASP-89 and 3’, 4’ hydroxyl groups formed hydrogen bond interaction with GLU-86. Following 6’ hydroxyl group formed hydrogen bond interaction with LYS-61. The C-13 hydroxyl formed hydrogen bond interaction with ASP-389 and ring C electrons formed pi-pi t-shaped interactions with HIS-63 and pi-alkyl interaction with LYS-388. Ring A electrons formed pi-carbon interaction with LYS-388 and LYS61 respectively. The ring C electrons and 4’ hydroxyl group of molecule MLR formed sulphur oxygen interaction with MET-344 and ring C electrons formed pi-alkyl interaction with LEU-396. The ring A 5 & 6 hydroxyl group formed hydrogen bond interaction with ASP-354 & LYS-538. Ring D 3’’ & 6’’ hydroxyl group formed hydrogen bond interaction with ASP-393 & GLY-277, 3’’ hydroxyl group formed pi-alkyl interaction with Mg ion. Ring D 5’’hydroxyl group formed hydrogen bond interaction with ARG-299. Ring E 6’’’ alkyl hydroxyl formed Carbon hydrogen interaction with LYS-391 and ring E oxygen, 3’’’ hydroxyl group and 6’’’ alkyl hydroxyl formed pi-alkyl interaction with Mg ions respectively.

In conclusion, the results from the current investigation not only identify a series of DEGs, but also analyze the significant modules, hub and target genes identification, and screening of small therapeutic molecules. In addition, in order to further verify the bioinformatics analysis data, the current investigation detected the expression levels of hub genes (HSP90AA1, EGFR, RPS13, RBX1, PAK1, FYN, ABL1, SMAD3, STAT3 and PRKCA) in a GDM. These hub genes might serve as potential diagnostic and prognostic biomarkers, and novel therapeutic targets in GDM.

## Acknowledgement

I thank Marian C Aldhous, Tommy’s Centre for Fetal and Maternal Health, Medical Research Council Centre for Reproductive Health, Queen’s Medical Research Institute, University of Edinburgh, Edinburgh, UK, very much, the author who deposited their profiling by high throughput sequencing dataset E-MTAB-6418, into the public ArrayExpress database.

## Ethical approval

This article does not contain any studies with human participants or animals performed by any of the authors.

## Informed consent

No informed consent because this study does not contain human or animals participants.

## Availability of data and materials

The datasets supporting the conclusions of this article are available in the ArrayExpress database (https://www.ebi.ac.uk/arrayexpress/) repository. [(E-MTAB-6418) (https://www.ebi.ac.uk/arrayexpress/experiments/E-MTAB-6418/?array=A-MEXP-2072]

## Consent for publication

Not applicable.

## Author Contributions

B. V - Writing original draft, and review and editing

C. V - Software and investigation

A. T - Formal analysis and validation

## Authors

Basavaraj Vastrad ORCID ID: <u>0000-0003-2202-7637</u> Anandkumar Tengli ORCID ID: 0000-0001-8076-928X Chanabasayya Vastrad ORCID ID: <u>0000-0003-3615-4450</u>

## References

1. Alfadhli EM. Gestational diabetes mellitus. Saudi Med J. 2015;36(4):399–406. doi:10.15537/smj.2015.4.10307

2. Lambrinoudaki I, Vlachou SA, Creatsas G. Genetics in gestational diabetes mellitus: association with incidence, severity, pregnancy outcome and response to treatment. Curr Diabetes Rev. 2010;6(6):393–399. doi:10.2174/157339910793499155

3. Chen P, Wang S, Ji J, Ge A, Chen C, Zhu Y, Xie N, Wang Y. Risk factors and management of gestational diabetes. Cell Biochem Biophys. 2015;71(2):689–694. doi:10.1007/s12013-014-0248-2

4. Tieu J, McPhee AJ, Crowther CA, Middleton P, Shepherd E. Screening for gestational diabetes mellitus based on different risk profiles and settings for improving maternal and infant health. Cochrane Database Syst Rev. 2017;8(8):CD007222. doi:10.1002/14651858.CD007222.pub4

5. Farrar D, Simmonds M, Bryant M, Sheldon TA, Tuffnell D, Golder S, Dunne F, Lawlor DA. Hyperglycaemia and risk of adverse perinatal outcomes: systematic review and meta-analysis. BMJ. 2016;354:i4694. doi:10.1136/bmj.i4694

6. Zhu N, Hou J, Wu Y, Li G, Liu J, Ma G, Chen B, Song Y.Identification of key genes in rheumatoid arthritis and osteoarthritis based on bioinformatics analysis. Medicine (Baltimore). 2018;97(22):e10997. doi:10.1097/MD.0000000000010997

7. Liu Y, Wang Y, Wang Y, Lv Y, Zhang Y, Wang H. Gene expression changes in arterial and venous endothelial cells exposed to gestational diabetes mellitus. Gynecol Endocrinol. 2020;36(9):791–795. doi:10.1080/09513590.2020.1712696

8. Zhang Q, He M, Wang J, Liu S, Cheng H, Cheng Y. Predicting of disease genes for gestational diabetes mellitus based on network and functional consistency. Eur J Obstet Gynecol Reprod Biol. 2015;186:91–96. doi:10.1016/j.ejogrb.2014.12.016

9. Chiswick CA, Reynolds RM, Denison FC, Drake AJ, Forbes S, Newby DE, Walker BR, Quenby S, Wray S, Weeks A, et al. Does metformin reduce excess birthweight in offspring of obese pregnant women? A randomised controlled trial of efficacy, exploration of mechanisms and evaluation of other pregnancy complications. Southampton (UK): NIHR Journals Library; August 2016.

10. Kolesnikov N, Hastings E, Keays M, Melnichuk O, Tang YA, Williams E, Dylag M, Kurbatova N, Brandizi M, Burdett T, et al. ArrayExpress update--simplifying data submissions. Nucleic Acids Res. 2015;43(Database issue):D1113-D1116. doi:10.1093/nar/gku1057

11. Ritchie ME, Phipson B, Wu D, Hu Y, Law CW, Shi W, Smyth GK. limma powers differential expression analyses for RNA-sequencing and microarray studies. Nucleic Acids Res. 2015;43(7):e47. doi:10.1093/nar/gkv007

12. Thomas PD. The Gene Ontology and the Meaning of Biological Function. Methods Mol Biol. 2017;1446:15L doi:10.1007/978-1-4939-3743-1_2

13. Fabregat A, Jupe S, Matthews L, Sidiropoulos K, Gillespie M, Garapati P, Haw R, Jassal B, Korninger F, May B et al The Reactome Pathway Knowledgebase. Nucleic Acids Res. 2018;46(D1):D649–D655. doi:10.1093/nar/gkx1132

14. Chen J, Bardes EE, Aronow BJ, Jegga AG. ToppGene Suite for gene list enrichment analysis and candidate gene prioritization. Nucleic Acids Res. 2009;37(Web Server issue):W305–W311. doi:10.1093/nar/gkp427

15. Szklarczyk D, Morris JH, Cook H, Kuhn M, Wyder S, Simonovic M, Santos A, Doncheva NT, Roth A, Bork P, et al. The STRING database in 2017: quality-controlled protein-protein association networks, made broadly accessible. Nucleic Acids Res. 2017;45(D1):D362–D368. doi:10.1093/nar/gkw937

16. Shannon P, Markiel A, Ozier O, Baliga NS, Wang JT, Ramage D, Amin N, Schwikowski B, Ideker T Cytoscape: a software environment for integrated models of biomolecular interaction networks. Genome Res 2003;13(11):2498–2504. doi:10.1101/gr.1239303

17. Przulj N, Wigle DA, Jurisica I. Functional topology in a network of protein interactions. Bioinformatics. 2004;20(3):340–348. doi:10.1093/bioinformatics/btg415

18. Nguyen TP, Liu WC, Jordán F. Inferring pleiotropy by network analysis: linked diseases in the human PPI network. BMC Syst Biol. 2011;5:179. doi:10.1186/1752-0509-5-179

19. Shi Z, Zhang B. Fast network centrality analysis using GPUs. BMC Bioinformatics. 2011;12:149. doi:10.1186/1471-2105-12-149

20. Fadhal E, Gamieldien J, Mwambene EC. Protein interaction networks as metric spaces: a novel perspective on distribution of hubs. BMC Syst Biol. 2014;8:6. doi:10.1186/1752-0509-8-6

21. Zaki N, Efimov D, Berengueres J. Protein complex detection using interaction reliability assessment and weighted clustering coefficient. BMC Bioinformatics. 2013;14:163. doi:10.1186/1471-2105-14

22. Fan Y, Xia J (2018) miRNet-Functional Analysis and Visual Exploration of miRNA-Target Interactions in a Network Context. Methods Mol Biol 1819:215–233. doi:10.1007/978-1-4939-8618-7_10

23. Zhou G, Soufan O, Ewald J, Hancock REW, Basu N, Xia J (2019) NetworkAnalyst 3.0: a visual analytics platform for comprehensive gene expression profiling and meta-analysis. Nucleic Acids Res 47:W234–W241. doi:10.1093/nar/gkz240

24. Robin X, Turck N, Hainard A, Tiberti N, Lisacek F, Sanchez JC, Müller M. pROC: an open-source package for R and S+ to analyze and compare ROC curves. BMC Bioinformatics 2011;12:77. doi:10.1186/1471-2105-12-77

25. Livak KJ, Schmittgen TD Analysis of relative gene expression data using real-time quantitative PCR and the 2(-Delta Delta C(T)) Method. Methods 2001;25:402–408. doi:10.1006/meth.2001.1262

26. Wierzchowski M, Dutkiewicz Z, Gielara-Korzańska A, Korzań methylthio derivatives as cytochromes P450 family 1 inhibitors. Chem Biol Drug Des. 2017;90(6):1226–1236. doi:10.1111/cbdd.13042

27. O’Boyle NM, Banck M, James CA, Morley C, Vandermeersch T, Hutchison GR. Open Babel: An open chemical toolbox. J Cheminform. 2011;3:33. doi:10.1186/1758-2946-3-33

28. Petchi RR, Vijaya C, Parasuraman S. Antidiabetic activity of polyherbal formulation in streptozotocin - nicotinamide induced diabetic wistar rats. J Tradit Complement Med. 2014;4(2):108–117. doi:10.4103/2225- 4110.126174

29. Gupta RC, Chang D, Nammi S, Bensoussan A, Bilinski K, Roufogalis BD. Interactions between antidiabetic drugs and herbs: an overview of mechanisms of action and clinical implications. Diabetol Metab Syndr. 2017;9:59. doi:10.1186/s13098-017-0254-9

30. Uusküla L, Rull K, Nagirnaja L, Laan M. Methylation allelic polymorphism (MAP) in chorionic gonadotropin beta5 (CGB5) and its association with pregnancy success. J Clin Endocrinol Metab. 2011;96(1):E199–E207. doi:10.1210/jc.2010-1647

31. Purwosunu Y, Sekizawa A, Farina A, Wibowo N, Okazaki S, Nakamura M, Samura O, Fujito N, Okai T.Cell-free mRNA concentrations of CRH, PLAC1, and selectin-P are increased in the plasma of pregnant women with preeclampsia. Prenat Diagn. 2007;27(8):772–777. doi:10.1002/pd.1780

32. Temur M, Serpim G, Tuzluoğlu S, Taş z FN, Ş E, Üstünyurt E. Comparison of serum human pregnancy-specific beta-1-glycoprotein 1 levels in pregnant women with or without preeclampsia. J Obstet Gynaecol. 2020;40(8):1074–1078. doi:10.1080/01443615.2019.1679734

33. Shimodaira M, Nakayama T, Sato I, Sato N, Izawa N, Mizutani Y, Furuya K, Yamamoto T. Estrogen synthesis genes CYP19A1, HSD3B1, and HSD3B2 in hypertensive disorders of pregnancy. Endocrine. 2012;42(3):700–707. doi:10.1007/s12020-012-9699-7

34. Xu T, Shao L, Wang A, Liang R, Lin Y, Wang G, Zhao Y, Hu J, Liu S. CD248 as a novel therapeutic target in pulmonary arterial hypertension. Clin Transl Med. 2020;10(5):e175. doi:10.1002/ctm2.175

35. Lin G, Wan X, Liu D, Wen Y, Yang C, Zhao C. COL1A1 as a potential new biomarker and therapeutic target for type 2 diabetes. Pharmacol Res. 2021;105436. doi:10.1016/j.phrs.2021.105436

36. Delfín DA, DeAguero JL, McKown EN. The Extracellular Matrix Protein ABI3BP in Cardiovascular Health and Disease. Front Cardiovasc Med. 2019;6:23. doi:10.3389/fcvm.2019.00023

37. Blindbæk SL, Schlosser A, Green A, Holmskov U, Sorensen GL, Grauslund J. Association between microfibrillar-associated protein 4 (MFAP4) and micro- and macrovascular complications in long-term type 1 diabetes mellitus. Acta Diabetol. 2017;54(4):367–372. doi:10.1007/s00592-016-0953-y

38. Bennett CE, Nsengimana J, Bostock JA, Cymbalista C, Futers TS, Knight BL, McCormack LJ, Prasad UK, Riches K, Rolton D, et al. CCAAT/enhancer binding protein alpha, beta and delta gene variants: associations with obesity related phenotypes in the Leeds Family Study. Diab Vasc Dis Res. 2010;7(3):195–203. doi:10.1177/1479164110366274

39. Killion EA, Reeves AR, El Azzouny MA, Yan QW, Surujon D, Griffin JD, Bowman TA, Wang C, Matthan NR, Klett EL, et al. A role for long-chain acyl-CoA synthetase-4 (ACSL4) in diet-induced phospholipid remodeling and obesity-associated adipocyte dysfunction. Mol Metab. 2018;9:43–56. doi:10.1016/j.molmet.2018.01.012

40. Cheng J, Song J, He X, Zhang M, Hu S, Zhang S, Yu Q, Yang P, Xiong F, Wang DW, et al. Loss of Mbd2 Protects Mice Against High-Fat Diet-Induced Obesity and Insulin Resistance by Regulating the Homeostasis of Energy Storage and Expenditure. Diabetes. 2016;65(11):3384–3395. doi:10.2337/db16-0151

41. An M, Ryu DR, Won Park J, Ha Choi J, Park EM, Eun Lee K, Woo M, Kim M. ULK1 prevents cardiac dysfunction in obesity through autophagy-meditated regulation of lipid metabolism. Cardiovasc Res. 2017;113(10):1137–1147. doi:10.1093/cvr/cvx064

42. Hofmann T, Weibert E, Ahnis A, Obbarius A, Elbelt U, Rose M, Klapp BF, Stengel A. Alterations of circulating NUCB2/nesfatin-1 during short term therapeutic improvement of anxiety in obese inpatients. Psychoneuroendocrinology. 2017;79:107–115. doi:10.1016/j.psyneuen.2017.02.021

43. Ma W, Lu S, Sun T, Wang X, Ma Y, Zhang X, Zhao R, Wang Y. Twist 1 regulates the expression of PPARγ during hormone-induced 3T3-L1 preadipocyte differentiation: a possible role in obesity and associated diseases. Lipids Health Dis. 2014;13:132. doi:10.1186/1476-511X-13-132

44. Rodríguez-Rodero S, Menéndez-Torre E, Fernández-Bayón G, Morales-Sánchez P, Sanz L, Turienzo E, González JJ, Martinez-Faedo C, Suarez-Gutiérrez L, Ares J, et al. Altered intragenic DNA methylation of HOOK2 gene in adipose tissue from individuals with obesity and type 2 diabetes. PLoS One. 2017;12(12):e0189153. doi:10.1371/journal.pone.0189153

45. Belalcazar LM, Papandonatos GD, McCaffery JM, Peter I, Pajewski NM, Erar B, Allred ND, Balasubramanyam A, Bowden DW, Brautbar A, et al. A common variant in the CLDN7/ELP5 locus predicts adiponectin change with lifestyle intervention and improved fitness in obese individuals with diabetes. Physiol Genomics. 2015;47(6):215–224. doi:10.1152/physiolgenomics.00109.2014

46. Reilly SM, Chiang SH, Decker SJ, Chang L, Uhm M, Larsen MJ, Rubin JR, Mowers J, White NM, Hochberg I, et al. An inhibitor of the protein kinases TBK1 and IKK-L improves obesity-related metabolic dysfunctions in mice. Nat Med. 2013;19(3):313–321. doi:10.1038/nm.3082

47. Wang L, Mazagova M, Pan C, Yang S, Brandl K, Liu J, Reilly SM, Wang Y, Miao Z, Loomba R, et al. YIPF6 controls sorting of FGF21 into COPII vesicles and promotes obesity. Proc Natl Acad Sci U S A. 2019;116(30):15184–15193. doi:10.1073/pnas.1904360116

48. Garcia-Valdes L, Campoy C, Hayes H, Florido J, Rusanova I, Miranda MT, McArdle HJ. The impact of maternal obesity on iron status, placental transferrin receptor expression and hepcidin expression in human pregnancy. Int J Obes (Lond). 2015;39(4):571–578. doi:10.1038/ijo.2015.3

49. Reeves VL, Trybula JS, Wills RC, Goodpaster BH, Dubé JJ, Kienesberger PC, Kershaw EE. Serum Autotaxin/ENPP2 correlates with insulin resistance in older humans with obesity. Obesity (Silver Spring). 2015;23(12):2371–2376. doi:10.1002/oby.21232

50. Lim R, Lappas M. Slit2 exerts anti-inflammatory actions in human placenta and is decreased with maternal obesity. Am J Reprod Immunol. 2015;73(1):66–78. doi:10.1111/aji.12334

51. Khalifeh-Soltani A, McKleroy W, Sakuma S, Cheung YY, Tharp K, Qiu Y, Turner SM, Chawla A, Stahl A, Atabai K. Mfge8 promotes obesity by mediating the uptake of dietary fats and serum fatty acids. Nat Med. 2014;20(2):175–183. doi:10.1038/nm.3450

52. Bidu C, Escoula Q, Bellenger S, Spor A, Galan M, Geissler A, Bouchot A, Dardevet D, Morio B, Cani PD, et al. The Transplantation of ω PUFA-Altered Gut Microbiota of fat-1 Mice to Wild-Type Littermates Prevents Obesity and Associated Metabolic Disorders. Diabetes. 2018;67(8):1512–1523. doi:10.2337/db17-1488

53. Leelalertlauw C, Korwutthikulrangsri M, Mahachoklertwattana P, Chanprasertyothin S, Khlairit P, Pongratanakul S, Poomthavorn P. Serum glypican 4 level in obese children and its relation to degree of obesity. Clin Endocrinol (Oxf). 2017;87(6):689–695. doi:10.1111/cen.13435

54. McCulloch LJ, Rawling TJ, Sjöholm K, Franck N, Dankel SN, Price EJ, Knight B, Liversedge NH, Mellgren G, Nystrom F, et al. COL6A3 is regulated by leptin in human adipose tissue and reduced in obesity. Endocrinology. 2015;156(1):134–146. doi:10.1210/en.2014-1042

55. Oberauer R, Rist W, Lenter MC, Hamilton BS, Neubauer H. EGFL6 is increasingly expressed in human obesity and promotes proliferation of adipose tissue-derived stromal vascular cells. Mol Cell Biochem. 2010;343(1-2):257–269. doi:10.1007/s11010-010-0521-7

56. Jargaud V, Bour S, Tercé F, Collet X, Valet P, Bouloumié A, Guillemot JC, Mauriège P, Jalkanen S, Stolen C, et al. Obesity of mice lacking VAP-1/SSAO by Aoc3 gene deletion is reproduced in mice expressing a mutated vascular adhesion protein-1 (VAP-1) devoid of amine oxidase activity. J Physiol Biochem. 2020;10.1007/s13105-020-00756-y. doi:10.1007/s13105-020-00756-y

57. Tan JT, McLennan SV, Williams PF, Rezaeizadeh A, Lo LW, Bonner JG, Twigg SM. Connective tissue growth factor/CCN-2 is upregulated in epididymal and subcutaneous fat depots in a dietary-induced obesity model. Am J Physiol Endocrinol Metab. 2013;304(12):E1291–E1302. doi:10.1152/ajpendo.00654.2012

58. Michurina SV, Ishchenko IY, Arkhipov SA, Klimontov VV, Rachkovskaya LN, Konenkov VI, Zavyalov EL. Effects of Melatonin, Aluminum Oxide, and Polymethylsiloxane Complex on the Expression of LYVE-1 in the Liver of Mice with Obesity and Type 2 Diabetes Mellitus. Bull Exp Biol Med. 2016;162(2):269–272. doi:10.1007/s10517-016-3592-y

59. Redonnet A, Bonilla S, Noël-Suberville C, Pallet V, Dabadie H, Gin H, Higueret P. Relationship between peroxisome proliferator-activated receptor gamma and retinoic acid receptor alpha gene expression in obese human adipose tissue. Int J Obes Relat Metab Disord. 2002;26(7):920–927. doi:10.1038/sj.ijo.0802025

60. Errera FI, Canani LH, Yeh E, Kague E, Armelin-Corrêa LM, Suzuki OT, Tschiedel B, Silva ME, Sertié AL, Passos-Bueno MR. COL18A1 is highly expressed during human adipocyte differentiation and the SNP c.1136C > T in its “frizzled” motif is associated with obesity in diabetes type 2 patients. An Acad Bras Cienc. 2008;80(1):167–177. doi:10.1590/s0001-37652008000100012

61. Paine A, Woeller CF, Zhang H, de la Luz Garcia-Hernandez M, Huertas N, Xing L, Phipps RP, Ritchlin CT. Thy1 is a positive regulator of osteoblast differentiation and modulates bone homeostasis in obese mice. FASEB J. 2018;32(6):3174–3183. doi:10.1096/fj.201701379R

62. Wu RX, Dong YY, Yang PW, Wang L, Deng YH, Zhang HW, Huang XY. CD36- and obesity-associated granulosa cells dysfunction. Reprod Fertil Dev. 2019;31(5):993–1001. doi:10.1071/RD18292

63. Gao X, van der Veen JN, Zhu L, Chaba T, Ordoñez M, Lingrell S, Koonen DP, Dyck JR, Gomez-Muñoz A, Vance DE, et al. Vagus nerve contributes to the development of steatohepatitis and obesity in phosphatidylethanolamine N-methyltransferase deficient mice. J Hepatol. 2015;62(4):913–920. doi:10.1016/j.jhep.2014.11.026

64. Parikh D, Riascos-Bernal DF, Egaña-Gorroño L, Jayakumar S, Almonte V, Chinnasamy P, Sibinga NES. Allograft inflammatory factor-1-like is not essential for age dependent weight gain or HFD-induced obesity and glucose insensitivity. Sci Rep. 2020;10(1):3594. doi:10.1038/s41598-020-60433-4

65. Bush NR, Allison AL, Miller AL, Deardorff J, Adler NE, Boyce WT. Socioeconomic Disparities in Childhood Obesity Risk: Association With an Oxytocin Receptor Polymorphism. JAMA Pediatr. 2017;171(1):61–67. doi:10.1001/jamapediatrics.2016.2332

66. Kim Y, Bayona PW, Kim M, Chang J, Hong S, Park Y, Budiman A, Kim YJ, Choi CY, Kim WS, et al. Macrophage Lamin A/C Regulates Inflammation and the Development of Obesity-Induced Insulin Resistance. Front Immunol. 2018;9:696. doi:10.3389/fimmu.2018.00696

67. Cereijo R, Quesada-López T, Gavaldà-Navarro A, Tarascó J, Pellitero S, Reyes M, Puig-Domingo M, Giralt M, Sánchez-Infantes D, Villarroya F. The chemokine CXCL14 is negatively associated with obesity and concomitant type-2 diabetes in humans. Int J Obes (Lond). 2021;10.1038/s41366-020-00732-y. doi:10.1038/s41366-020-00732-y

68. Xie L, Wang PX, Zhang P, Zhang XJ, Zhao GN, Wang A, Guo J, Zhu X, Zhang Q, Li H. DKK3 expression in hepatocytes defines susceptibility to liver steatosis and obesity. J Hepatol. 2016;65(1):113–124. doi:10.1016/j.jhep.2016.03.008

69. Tabata M, Kadomatsu T, Fukuhara S, Miyata K, Ito Y, Endo M, Urano T, Zhu HJ, Tsukano H, Tazume H, et al. Angiopoietin-like protein 2 promotes chronic adipose tissue inflammation and obesity-related systemic insulin resistance. Cell Metab. 2009;10(3):178–188. doi:10.1016/j.cmet.2009.08.003

70. 70. 70.Zhu Q, Xue K, Guo HW, Deng FF, Yang YH. Interaction of the CMTM7 rs347134 Polymorphism with Dietary Patterns and the Risk of Obesity in Han Chinese Male Children. Int J Environ Res Public Health. 2020;17(5):1515. doi:10.3390/ijerph17051515

71. Wang K, Zhou Q, He Q, Tong G, Zhao Z, Duan T. The possible role of AhR in the protective effects of cigarette smoke on preeclampsia. Med Hypotheses. 2011;77(5):872–874. doi:10.1016/j.mehy.2011.07.061

72. Gratton AM, Ye L, Brownfoot FC, Hannan NJ, Whitehead C, Cannon P, Deo M, Fuller PJ, Tong S, Kaitu’u-Lino TJ. Steroid sulfatase is increased in the placentas and whole blood of women with early-onset preeclampsia. Placenta. 2016;48:72–79. doi:10.1016/j.placenta.2016.10.008

73. Wan L, Sun D, Xie J, Du M, Wang P, Wang M, Lei Y, Wang H, Wang H, Dong M. Declined placental PLAC1 expression is involved in preeclampsia. Medicine (Baltimore). 2019;98(44):e17676. doi:10.1097/MD.0000000000017676

74. Pan T, He G, Chen M, Bao C, Chen Y, Liu G, Zhou M, Li S, Xu W, Liu X. Abnormal CYP11A1 gene expression induces excessive autophagy, contributing to the pathogenesis of preeclampsia. Oncotarget. 2017;8(52):89824–89836. doi:10.18632/oncotarget.21158

75. Zhao L, Triche EW, Walsh KM, Bracken MB, Saftlas AF, Hoh J, Dewan AT. Genome-wide association study identifies a maternal copy-number deletion in PSG11 enriched among preeclampsia patients. BMC Pregnancy Childbirth. 2012;12:61. doi:10.1186/1471-2393-12-61

76. Li W, Geng L, Liu X, Gui W, Qi H. Recombinant adiponectin alleviates abortion in mice by regulating Th17/Treg imbalance via p38MAPK-STAT5 pathway. Biol Reprod. 2019;100(4):1008–1017. doi:10.1093/biolre/ioy251

77. Gierman LM, Silva GB, Pervaiz Z, Rakner JJ, Mundal SB, Thaning AJ, Nervik I, Elschot M, Mathew S, Thomsen LCV, et al. TLR3 expression by maternal and fetal cells at the maternal-fetal interface in normal and preeclamptic pregnancies. J Leukoc Biol. 2020;10.1002/JLB.3MA0620-728RR. doi:10.1002/JLB.3MA0620-728RR

78. Piñuñuri R, Castaño-Moreno E, Llanos MN, Ronco AM. Epigenetic regulation of folate receptor-α (FOLR1) in human placenta of preterm newborns. Placenta. 2020;94:20–25. doi:10.1016/j.placenta.2020.03.009

79. White BG, Williams SJ, Highmore K, Macphee DJ. Small heat shock protein 27 (Hsp27) expression is highly induced in rat myometrium during late pregnancy and labour. Reproduction. 2005;129(1):115–126. doi:10.1530/rep.1.00426

80. Torres-Salazar Q, Martínez-López Y, Reyes-Romero M, Pérez-Morales R, Sifuentes-Álvarez A, Salvador-Moysén J. Differential Methylation in Promoter Regions of the Genes NR3C1 and HSP90AA1, Involved in the Regulation, and Bioavailability of Cortisol in Leukocytes of Women With Preeclampsia. Front Med (Lausanne). 2020;7:206. doi:10.3389/fmed.2020.00206

81. Xu Y, Sui L, Qiu B, Yin X, Liu J, Zhang X. ANXA4 promotes trophoblast invasion via the PI3K/Akt/eNOS pathway in preeclampsia. Am J Physiol Cell Physiol. 2019;316(4):C481–C491. doi:10.1152/ajpcell.00404.2018

82. Kaitu’u-Lino TJ, Brownfoot FC, Hastie R, Chand A, Cannon P, Deo M, Tuohey L, Whitehead C, Hannan NJ, Tong S. Activating Transcription Factor 3 Is Reduced in Preeclamptic Placentas and Negatively Regulates sFlt-1 (Soluble fms-Like Tyrosine Kinase 1), Soluble Endoglin, and Proinflammatory Cytokines in Placenta. Hypertension. 2017;70(5):1014–1024. doi:10.1161/HYPERTENSIONAHA.117.09548

83. Yung C, MacDonald TM, Walker SP, Cannon P, Harper A, Pritchard N, Hannan NJ, Kaitu’u-Lino TJ, Tong S. Death associated protein kinase 1 (DAPK-1) is increased in preeclampsia. Placenta. 2019;88:1–7. doi:10.1016/j.placenta.2019.09.010

84. Zhu L, Lv R, Kong L, Li X. Reduced methylation downregulates CD39/ENTPD1 and ZDHHC14 to suppress trophoblast cell proliferation and invasion: Implications in preeclampsia. Pregnancy Hypertens. 2018;14:59–67. doi:10.1016/j.preghy.2018.03.012

85. Griesshammer M, Sadjadian P, Wille K. Contemporary management of patients with BCR-ABL1-negative myeloproliferative neoplasms during pregnancy. Expert Rev Hematol. 2018;11(9):697–706. doi:10.1080/17474086.2018.1506325

86. Textoris J, Ivorra D, Ben Amara A, Sabatier F, Ménard JP, Heckenroth H, Bretelle F, Mege JL. Evaluation of current and new biomarkers in severe preeclampsia: a microarray approach reveals the VSIG4 gene as a potential blood biomarker. PLoS One. 2013;8(12):e82638. doi:10.1371/journal.pone.0082638

87. Kelemu T, Erlandsson L, Seifu D, Abebe M, Teklu S, Storry JR, Hansson SR. Association of Maternal Regulatory Single Nucleotide Polymorphic CD99 Genotype with Preeclampsia in Pregnancies Carrying Male Fetuses in Ethiopian Women. Int J Mol Sci. 2020;21(16):5837. doi:10.3390/ijms21165837

88. Aref S, Goda H. Increased VWF antigen levels and decreased ADAMTS13 activity in preeclampsia. Hematology. 2013;18(4):237–241. doi:10.1179/1607845412Y.0000000070

89. Martineau T, Boutin M, Côté AM, Maranda B, Bichet DG, Auray-Blais C. Tandem mass spectrometry analysis of urinary podocalyxin and podocin in the investigation of podocyturia in women with preeclampsia and Fabry disease patients. Clin Chim Acta. 2019;495:67–75. doi:10.1016/j.cca.2019.03.1615

90. Onyangunga O, Moodley J, Odun-Ayo F, Naicker T. Immunohistochemical localization of podoplanin in the placentas of HIV-positive preeclamptic women. Turk J Med Sci. 2018;48(5):916–924. doi:10.3906/sag-1706-88

91. Li L, Zhang X, Hong SL, Chen Y, Ren GH. Long non-coding HOTTIP regulates preeclampsia by inhibiting RND3. Eur Rev Med Pharmacol Sci. 2018;22(11):3277–3285. doi:10.26355/eurrev_201806_15146

92. Gogiel T, Galewska Z, Romanowicz L, Jaworski S, Bań eclampsia-associated alterations in decorin, biglycan and versican of the umbilical cord vein wall. Eur J Obstet Gynecol Reprod Biol. 2007;134(1):51–56. doi:10.1016/j.ejogrb.2006.10.003

93. Hirschi KM, Tsai KYF, Davis T, Clark JC, Knowlton MN, Bikman BT, Reynolds PR, Arroyo JA. Growth arrest-specific protein-6/AXL signaling induces preeclampsia in rats†. Biol Reprod. 2020;102(1):199–210. doi:10.1093/biolre/ioz140

94. Arishe OO, Ebeigbe AB, Webb RC. Mechanotransduction and Uterine Blood Flow in Preeclampsia: The Role of Mechanosensing Piezo 1 Ion Channels. Am J Hypertens. 2020;33(1):1–9. doi:10.1093/ajh/hpz158

95. Ji Y, Zhou L, Wang G, Qiao Y, Tian Y, Feng Y. Role of LAMA4 Gene in Regulating Extravillous Trophoblasts in Pathogenesis of Preeclampsia. Med Sci Monit. 2019;25:9630–9636. doi:10.12659/MSM.917402

96. Du F, Zhang Y, Xu Q, Teng Y, Tao M, Chen AF, Jiang R. Preeclampsia serum increases CAV1 expression and cell permeability of human renal glomerular endothelial cells via down-regulating miR-199a-5p, miR-199b-5p, miR-204. Placenta. 2020;99:141–151. doi:10.1016/j.placenta.2020.07.011

97. Shimanuki Y, Mitomi H, Fukumura Y, Makino S, Itakura A, Yao T, Takeda S. Alteration of Delta-like ligand 1 and Notch 1 receptor in various placental disorders with special reference to early onset preeclampsia. Hum Pathol. 2015;46(8):1129–1137. doi:10.1016/j.humpath.2015.03.013

98. Todd N, McNally R, Alqudah A, Jerotic D, Suvakov S, Obradovic D, Hoch D, Hombrebueno JR, Campos GL, Watson CJ, et al. Role of A Novel Angiogenesis FKBPL-CD44 Pathway in Preeclampsia Risk Stratification and Mesenchymal Stem Cell Treatment. J Clin Endocrinol Metab. 2021;106(1):26–41. doi:10.1210/clinem/dgaa403

99. Ding H, Dai Y, Lei Y, Wang Z, Liu D, Li R, Shen L, Gu N, Zheng M, Zhu X, et al. Upregulation of CD81 in trophoblasts induces an imbalance of Treg/Th17 cells by promoting IL-6 expression in preeclampsia. Cell Mol Immunol. 2019;16(1):302–312. doi:10.1038/s41423-018-0186-9

100. Brkić J, Dunk C, Shan Y, O’Brien JA, Lye P, Qayyum S, Yang P, Matthews SG, Lye SJ, Peng C. Differential Role of Smad2 and Smad3 in the Acquisition of an Endovascular Trophoblast-Like Phenotype and Preeclampsia. Front Endocrinol (Lausanne). 2020;11:436. doi:10.3389/fendo.2020.00436

101. Yang X, Ding Y, Yang M, Yu L, Hu Y, Deng Y. Nestin Improves Preeclampsia-Like Symptoms by Inhibiting Activity of Cyclin-Dependent Kinase 5. Kidney Blood Press Res. 2018;43(2):616–627. doi:10.1159/000489146

102. Lala PK, Nandi P. Mechanisms of trophoblast migration, endometrial angiogenesis in preeclampsia: The role of decorin. Cell Adh Migr. 2016;10(1-2):111–125. doi:10.1080/19336918.2015.1106669

103. Zhao L, Dewan AT, Bracken MB. Association of maternal AGTR1 polymorphisms and preeclampsia: a systematic review and meta-analysis. J Matern Fetal Neonatal Med. 2012;25(12):2676–2680. doi:10.3109/14767058.2012.708370

104. Lim R, Barker G, Lappas M. SLIT3 is increased in supracervical human foetal membranes and in labouring myometrium and regulates pro-inflammatory mediators. Am J Reprod Immunol. 2014;71(4):297–311. doi:10.1111/aji.12181

105. Kristensen K, Wide-Swensson D, Schmidt C, Blirup-Jensen S, Lindström V, Strevens H, Grubb A. Cystatin C, beta-2-microglobulin and beta-trace protein in pre-eclampsia. Acta Obstet Gynecol Scand. 2007;86(8):921–926. doi:10.1080/00016340701318133

106. Tang J, Wang D, Lu J, Zhou X. MiR-125b participates in the occurrence of preeclampsia by regulating the migration and invasion of extravillous trophoblastic cells through STAT3 signaling pathway. J Recept Signal Transduct Res. 2020;1–7. doi:10.1080/10799893.2020.1806318

107. Abid N, Embola J, Tryfonos Z, Bercher J, Ashton SV, Khalil A, Thilaganathan B, Cartwright JE, Whitley GS. Regulation of stanniocalcin-1 secretion by BeWo cells and first trimester human placental tissue from normal pregnancies and those at increased risk of developing preeclampsia. FASEB J. 2020;34(5):6086–6098. doi:10.1096/fj.201902426R

108. Namlı Kalem M, Kalem Z, Yüce T, Soylemez F. ADAMTS 1, 4, 12, and 13 levels in maternal blood, cord blood, and placenta in preeclampsia. Hypertens Pregnancy. 2018;37(1):9–17. doi:10.1080/10641955.2017.1397690

109. Majchrzak-Celi ska A, Kosicka K, Paczkowska J, Główka FK, Brę rowicz GH, Krzy cin M, Siemią owska A, Szaumkessel M, Baer-bo ś tk Dubowska W. HSD11B2, RUNX3, and LINE-1 Methylation in Placental DNA of Hypertensive Disorders of Pregnancy Patients. Reprod Sci. 2017;24(11):1520–1531. doi:10.1177/1933719117692043

110. Shimodaira M, Nakayama T, Sato I, Sato N, Izawa N, Mizutani Y, Furuya K, Yamamoto T.Estrogen synthesis genes CYP19A1, HSD3B1, and HSD3B2 in hypertensive disorders of pregnancy. Endocrine. 2012;42(3):700–707. doi:10.1007/s12020-012-9699-7

111. Saxena R, Bjonnes A, Prescott J, Dib P, Natt P, Lane J, Lerner M, Cooper JA, Ye Y, Li KW, et al. Genome-wide association study identifies variants in casein kinase II (CSNK2A2) to be associated with leukocyte telomere length in a Punjabi Sikh diabetic cohort. Circ Cardiovasc Genet. 2014;7(3):287–295. doi:10.1161/CIRCGENETICS.113.000412

112. Yang W, Wang J, Chen Z, Chen J, Meng Y, Chen L, Chang Y, Geng B, Sun L, Dou L, et al. NFE2 Induces miR-423-5p to Promote Gluconeogenesis and Hyperglycemia by Repressing the Hepatic FAM3A-ATP-Akt Pathway. Diabetes. 2017;66(7):1819–1832. doi:10.2337/db16- 1172

113. Gloyn AL, Desai M, Clark A, Levy JC, Holman RR, Frayling TM, Hattersley AT, Ashcroft SJ. Human calcium/calmodulin-dependent protein kinase II gamma gene (CAMK2G): cloning, genomic structure and detection of variants in subjects with type II diabetes. Diabetologia. 2002;45(4):580–583. doi:10.1007/s00125-002-0779-8

114. Li JY, Tao F, Wu XX, Tan YZ, He L, Lu H. Common RASGRP1 Gene Variants That Confer Risk of Type 2 Diabetes. Genet Test Mol Biomarkers. 2015;19(8):439–443. doi:10.1089/gtmb.2015.0005

115. Afarideh M, Zaker Esteghamati V, Ganji M, Heidari B, Esteghamati S, Lavasani S, Ahmadi M, Tafakhori A, Nakhjavani M, Esteghamati A.Associations of Serum S100B and S100P With the Presence and Classification of Diabetic Peripheral Neuropathy in Adults With Type 2 Diabetes: A Case-Cohort Study. Can J Diabetes. 2019;43(5):336–344.e2. doi:10.1016/j.jcjd.2019.01.003

116. Zhang S, Xiao J, Ren Q, Han X, Tang Y, Yang W, Zhou X, Ji L. Association of serine racemase gene variants with type 2 diabetes in the Chinese Han population. J Diabetes Investig. 2014;5(3):286–289. doi:10.1111/jdi.12145

117. Robbins RD, Tersey SA, Ogihara T, Gupta D, Farb TB, Ficorilli J, Bokvist K, Maier B, Mirmira RG. Inhibition of deoxyhypusine synthase enhances islet {beta} cell function and survival in the setting of endoplasmic reticulum stress and type 2 diabetes. J Biol Chem. 2010;285(51):39943–39952. doi:10.1074/jbc.M110.170142

118. Belgardt BF, Lammert E. DYRK1A: A Promising Drug Target for Islet Transplant-Based Diabetes Therapies. Diabetes. 2016;65(6):1496–1498. doi:10.2337/dbi16-0013

119. Yoon CH, Choi YE, Cha YR, Koh SJ, Choi JI, Kim TW, Woo SJ, Park YB, Chae IH, Kim HS. Diabetes-Induced Jagged1 Overexpression in Endothelial Cells Causes Retinal Capillary Regression in a Murine Model of Diabetes Mellitus: Insights Into Diabetic Retinopathy. Circulation. 2016;134(3):233–247. doi:10.1161/CIRCULATIONAHA.116.014411

120. Gaikwad AB, Gupta J, Tikoo K. Epigenetic changes and alteration of Fbn1 and Col3A1 gene expression under hyperglycaemic and hyperinsulinaemic conditions. Biochem J. 2010;432(2):333–341. doi:10.1042/BJ20100414

121. Alessi MC, Nicaud V, Scroyen I, Lange C, Saut N, Fumeron F, Marre M, Lantieri O, Fontaine-Bisson B, Juhan-Vague I, et al. Association of vitronectin and plasminogen activator inhibitor-1 levels with the risk of metabolic syndrome and type 2 diabetes mellitus. Results from the D.E.S.I.R. prospective cohort. Thromb Haemost. 2011;106(3):416–422. doi:10.1160/TH11-03-0179

122. Yang Q, Wang WW, Ma P, Ma ZX, Hao M, Adelusi TI, Lei-Du, Yin XX, Lu Q. Swimming training alleviated insulin resistance through Wnt3a/β-catenin signaling in type 2 diabetic rats. Iran J Basic Med Sci. β 2017;20(11):1220–1226. doi:10.22038/IJBMS.2017.9473

123. Fang H, Luo X, Wang Y, Liu N, Fu C, Wang H, Fang Y, Shi W, Zhang Y, Zeng C, et al. Correlation between single nucleotide polymorphisms of the ACTA2 gene and coronary artery stenosis in patients with type 2 diabetes mellitus. Exp Ther Med. 2014;7(4):970–976. doi:10.3892/etm.2014.1510

124. Xu X, Fang K, Wang L, Liu X, Zhou Y, Song Y. Local Application of Semaphorin 3A Combined with Adipose-Derived Stem Cell Sheet and Anorganic Bovine Bone Granules Enhances Bone Regeneration in Type 2 Diabetes Mellitus Rats. Stem Cells Int. 2019;2019:2506463. doi:10.1155/2019/2506463

125. Zhao K, Ding W, Zhang Y, Ma K, Wang D, Hu C, Liu J, Zhang X. Variants in the RARRES2 gene are associated with serum chemerin and increase the risk of diabetic kidney disease in type 2 diabetes. Int J Biol Macromol. 2020;165(Pt A):1574–1580. doi:10.1016/j.ijbiomac.2020.10.030

126. Fisher E, Schreiber S, Joost HG, Boeing H, Döring F. A two-step association study identifies CAV2 rs2270188 single nucleotide polymorphism interaction with fat intake in type 2 diabetes risk. J Nutr. 2011;141(2):177–181. doi:10.3945/jn.110.124206

127. Meng S, Cao JT, Zhang B, Zhou Q, Shen CX, Wang CQ. Downregulation of microRNA-126 in endothelial progenitor cells from diabetes patients, impairs their functional properties, via target gene Spred-1. J Mol Cell Cardiol. 2012;53(1):64–72. doi:10.1016/j.yjmcc.2012.04.003

128. Santiago JL, Martínez A, de la Calle H, Fernández-Arquero M, Figueredo MA, de la Concha EG, Urcelay E. Evidence for the association of the SLC22A4 and SLC22A5 genes with type 1 diabetes: a case control study. BMC Med Genet. 2006;7:54. doi:10.1186/1471-2350-7-54

129. Auburger G, Gispert S, Lahut S, Omür O, Damrath E, Heck M, Başak N. 12q24 locus association with type 1 diabetes: SH2B3 or ATXN2?. World J Diabetes. 2014;5(3):316–327. doi:10.4239/wjd.v5.i3.316

130. Qu HQ, Marchand L, Szymborski A, Grabs R, Polychronakos C. The association between type 1 diabetes and the ITPR3 gene polymorphism due to linkage disequilibrium with HLA class II. Genes Immun. 2008;9(3):264–266. doi:10.1038/gene.2008.12

131. Śnit M, Nabrdalik K, Długaszek M, Gumprecht J, Trautsolt W, Górczy ska-Kosiorz S, Grzeszczak W. Association of rs 3807337 poly ń of CALD1 gene with diabetic nephropathy occurrence in type Morphism 1 diabetes - preliminary results of a family-based study. Endokrynol Pol. 2017;68(1):13–17. doi:10.5603/EP.2017.0003

132. Hjortebjerg R, Tarnow L, Jorsal A, Parving HH, Rossing P, Bjerre M, Frystyk J. IGFBP-4 Fragments as Markers of Cardiovascular Mortality in Type 1 Diabetes Patients With and Without Nephropathy. J Clin Endocrinol Metab. 2015;100(8):3032–3040. doi:10.1210/jc.2015-2196

133. Krishnan M, Murphy R, Okesene-Gafa KAM, Ji M, Thompson JMD, Taylor RS, Merriman TR, McCowan LME, McKinlay CJD. The Pacific-specific CREBRF rs373863828 allele protects against gestational diabetes mellitus in M ori and Pacific women with obesity. Diabetologia. 2020;63(10)2169–2176. doi:10.1007/s00125-020-05202-8):2169

134. Hu S, Yan J, You Y, Yang G, Zhou H, Li X, Liao X, Tan H. Association of polymorphisms in STRA6 gene with gestational diabetes mellitus in a Chinese Han population. Medicine (Baltimore). 2019;98(11):e14885. doi:10.1097/MD.0000000000014885

135. Martins RS, Ahmed T, Farhat S, Shahid S, Fatima SS. Epidermal growth factor receptor rs17337023 polymorphism in hypertensive gestational diabetic women: A pilot study. World J Diabetes. 2019;10(7):396–402. doi:10.4239/wjd.v10.i7.396

136. Prieto-Sánchez MT, Ruiz-Palacios M, Blanco-Carnero JE, Pagan A, Hellmuth C, Uhl O, Peissner W, Ruiz-Alcaraz AJ, Parrilla JJ, Koletzko B, et al. Placental MFSD2a transporter is related to decreased DHA in cord blood of women with treated gestational diabetes. Clin Nutr. 2017;36(2):513–521. doi:10.1016/j.clnu.2016.01.014

137. Sugulle M, Dechend R, Herse F, Weedon-Fekjaer MS, Johnsen GM, Brosnihan KB, Anton L, Luft FC, Wollert KC, Kempf T, et al. Circulating and placental growth-differentiation factor 15 in preeclampsia and in pregnancy complicated by diabetes mellitus. Hypertension. 2009;54(1):106–112. doi:10.1161/HYPERTENSIONAHA.109.130583

138. Zhao H, Tao S. MiRNA-221 protects islet β cell function in gestational diabetes mellitus by targeting PAK1. Biochem Biophys Res Commun. 2019;520(1):218–224. doi:10.1016/j.bbrc.2019.09.139

139. Siddiqui K, George TP, Nawaz SS, Joy SS. VCAM-1, ICAM-1 and selectins in gestational diabetes mellitus and the risk for vascular disorders. Future Cardiol. 2019;15(5):339–346. doi:10.2217/fca-2018-0042

140. Han N, Fang HY, Jiang JX, Xu Q. Downregulation of microRNA-873 attenuates insulin resistance and myocardial injury in rats with gestational diabetes mellitus by upregulating IGFBP2. Am J Physiol Endocrinol Metab. 2020;318(5):E723–E735. doi:10.1152/ajpendo.00555.2018

141. Lappas M. Insulin-like growth factor-binding protein 1 and 7 concentrations are lower in obese pregnant women, women with gestational diabetes and their fetuses. J Perinatol. 2015;35(1):32–38. doi:10.1038/jp.2014.144

142. Wang F, Xu C, Reece EA, Li X, Wu Y, Harman C, Yu J, Dong D, Wang C, Yang P, et al. Protein kinase C-alpha suppresses autophagy and induces neural tube defects via miR-129-2 in diabetic pregnancy. Nat Commun. 2017;8:15182. doi:10.1038/ncomms15182

143. Artunc-Ulkumen B, Ulucay S, Pala HG, Cam S. Maternal serum ADAMTS-9 levels in gestational diabetes: a pilot study. J Matern Fetal Neonatal Med. 2017;30(12):1442–1445. doi:10.1080/14767058.2016.1219717

144. Blois SM, Gueuvoghlanian-Silva BY, Tirado-González I, Torloni MR, Freitag N, Mattar R, Conrad ML, Unverdorben L, Barrientos G, Knabl J, et al. Getting too sweet: galectin-1 dysregulation in gestational diabetes mellitus. Mol Hum Reprod. 2014;20(7):644–649. doi:10.1093/molehr/gau021

145. Vacínová G, Vejražková D, Lukášová P, Lischková O, Dvo áková K, Rusina R, Holmerová I, Va ková H, V elák J, Bendlová ^ř^, et al. ň č B Associations of polymorphisms in the candidate genes for Alzheimer’s disease BIN1, CLU, CR1 and PICALM with gestational diabetes and impaired glucose tolerance. Mol Biol Rep. 2017;44(2):227–231. doi:10.1007/s11033-017-4100-9

146. Vilmi-Kerälä T, Lauhio A, Tervahartiala T, Palomäki O, Uotila J, Sorsa T, Palomäki A. Subclinical inflammation associated with prolonged TIMP-1 upregulation and arterial stiffness after gestational diabetes mellitus: a hospital-based cohort study. Cardiovasc Diabetol. 2017;16(1):49. doi:10.1186/s12933-017-0530-x

147. Aquila G, Vieceli Dalla Sega F, Marracino L, Pavasini R, Cardelli LS, Piredda A, Scoccia A, Martino V, Fortini F, Bononi I et al Ticagrelor Increases SIRT1 and HES1 mRNA Levels in Peripheral Blood Cells from Patients with Stable Coronary Artery Disease and Chronic Obstructive Pulmonary Disease. Int J Mol Sci. 2020;21(5):1576. doi:10.3390/ijms21051576

148. Chen Y, Wu S, Zhang XS, Wang DM, Qian CY. MicroRNA-489 promotes cardiomyocyte apoptosis induced by myocardial ischemia-reperfusion injury through inhibiting SPIN1. Eur Rev Med Pharmacol Sci. 2019;23(15):6683–6690. doi:10.26355/eurrev_201908_18559

149. Xie H, Zhang E, Hong N, Fu Q, Li F, Chen S, Yu Y, Sun K. Identification of TBX2 and TBX3 variants in patients with conotruncal heart defects by target sequencing. Hum Genomics. 2018;12(1):44. doi:10.1186/s40246-018-0176-0

150. Zhang S, Lin X, Li G, Shen X, Niu D, Lu G, Fu X, Chen Y, Cui M, Bai Y. Knockout of Eva1a leads to rapid development of heart failure by impairing autophagy. Cell Death Dis. 2017;8(2):e2586. doi:10.1038/cddis.2017.17

151. Aspit L, Levitas A, Etzion S, Krymko H, Slanovic L, Zarivach R, Etzion Y, Parvari R. CAP2 mutation leads to impaired actin dynamics and associates with supraventricular tachycardia and dilated cardiomyopathy. J Med Genet. 2019;56(4):228–235. doi:10.1136/jmedgenet-2018-105498

152. Akadam-Teker B, Ozkara G, Kurnaz-Gomleksiz O, Bugra Z, Teker E, Ozturk O, Yilmaz-Aydogan H. BMP1 5’UTR +L T/C gene variation: can be a predictive marker for serum HDL and apoprotein A1 levels in male patients with coronary heart disease. Mol Biol Rep. 2018;45(5):1269–1276. doi:10.1007/s11033-018-4283-8

153. Jiang B, Liu Y, Liang P, Li Y, Liu Z, Tong Z, Lv Q, Liu M, Xiao X. MicroRNA-126a-5p enhances myocardial ischemia-reperfusion injury through suppressing Hspb8 expression. Oncotarget. 2017;8(55):94172–94187. doi:10.18632/oncotarget.21613

154. Cetinkaya A, Berge B, Sen-Hild B, Troidl K, Gajawada P, Kubin N, Valeske K, Schranz D, Akintürk H, Schönburg M, et al. Radixin Relocalization and Nonmuscle α ctinin Expression Are Features of -A Remodeling Cardiomyocytes in Adult Patients with Dilated Cardiomyopathy. Dis Markers. 2020;2020:9356738. doi:10.1155/2020/9356738

155. Grond-Ginsbach C, Weber R, Haas J, Orberk E, Kunz S, Busse O, Hausser I, Brandt T, Wildemann B. Mutations in the COL5A1 coding sequence are not common in patients with spontaneous cervical artery dissections. Stroke. 1999;30(9):1887–1890. doi:10.1161/01.str.30.9.1887

156. Dong Y, Zhai W, Xu Y. Bioinformatic gene analysis for potential biomarkers and therapeutic targets of diabetic nephropathy associated renal cell carcinoma. Transl Androl Urol. 2020;9(6):2555–2571. doi:10.21037/tau- 19-911

157. Chardon JW, Smith AC, Woulfe J, Pena E, Rakhra K, Dennie C, Beaulieu C, Huang L, Schwartzentruber J, Hawkins C, et al. LIMS2 mutations are associated with a novel muscular dystrophy, severe cardiomyopathy and triangular tongues. Clin Genet. 2015;88(6):558–564. doi:10.1111/cge.12561

158. Chen H, Huang XN, Yan W, Chen K, Guo L, Tummalapali L, Dedhar S, St-Arnaud R, Wu C, Sepulveda JL. Role of the integrin-linked kinase/PINCH1/alpha-parvin complex in cardiac myocyte hypertrophy. Lab Invest. 2005;85(11):1342–1356. doi:10.1038/labinvest.3700345

159. Yamada Y, Sakuma J, Takeuchi I, Yasukochi Y, Kato K, Oguri M, Fujimaki T, Horibe H, Muramatsu M, Sawabe M, et al. Identification of EGFLAM, SPATC1L and RNASE13 as novel susceptibility loci for aortic aneurysm in Japanese individuals by exome-wide association studies. Int J Mol Med. 2017;39(5):1091–1100. doi:10.3892/ijmm.2017.2927

160. Hu YW, Guo FX, Xu YJ, Li P, Lu ZF, McVey DG, Zheng L, Wang Q, Ye JH, Kang CM, et al. Long noncoding RNA NEXN-AS1 mitigates atherosclerosis by regulating the actin-binding protein NEXN. J Clin Invest. 2019;129(3):1115–1128. doi:10.1172/JCI98230

161. Bobik A, Kalinina N. Tumor necrosis factor receptor and ligand superfamily family members TNFRSF14 and LIGHT: new players in human atherogenesis. Arterioscler Thromb Vasc Biol. 2001;21(12):1873–1875

162. Schwanekamp JA, Lorts A, Sargent MA, York AJ, Grimes KM, Fischesser DM, Gokey JJ, Whitsett JA, Conway SJ, Molkentin JD. TGFBI functions similar to periostin but is uniquely dispensable during cardiac injury. PLoS One. 2017;12(7):e0181945. doi:10.1371/journal.pone.0181945

163. Liu C, Zhang H, Chen Y, Wang S, Chen Z, Liu Z, Wang J. Identifying RBM47, HCK, CD53, TYROBP, and HAVCR2 as Hub Genes in Advanced Atherosclerotic Plaques by Network-Based Analysis and Validation. Front Genet. 2021;11:602908. doi:10.3389/fgene.2020.602908

164. Schroer AK, Bersi MR, Clark CR, Zhang Q, Sanders LH, Hatzopoulos AK, Force TL, Majka SM, Lal H, Merryman WD. Cadherin-11 blockade reduces inflammation-driven fibrotic remodeling and improves outcomes after myocardial infarction. JCI Insight. 2019;4(18):e131545.doi:10.1172/jci.insight.131545

165. Raza ST, Abbas S, Eba A, Karim F, Wani IA, Rizvi S, Zaidi A, Mahdi F. Association of COL4A1 (rs605143, rs565470) and CD14 (rs2569190) genes polymorphism with coronary artery disease. Mol Cell Biochem. 2018;445(1-2):117–122. doi:10.1007/s11010-017-3257-9

166. Yang W, Ng FL, Chan K, Pu X, Poston RN, Ren M, An W, Zhang R, Wu J, Yan S, et al. Coronary-Heart-Disease-Associated Genetic Variant at the COL4A1/COL4A2 Locus Affects COL4A1/COL4A2 Expression, Vascular Cell Survival, Atherosclerotic Plaque Stability and Risk of Myocardial Infarction. PLoS Genet. 2016;12(7):e1006127. doi:10.1371/journal.pgen.1006127

167. Azuaje F, Zhang L, Jeanty C, Puhl SL, Rodius S, Wagner DR. Analysis of a gene co-expression network establishes robust association between Col5a2 and ischemic heart disease. BMC Med Genomics. 2013;6:13. doi:10.1186/1755-8794-6-13

168. Durbin MD, O’Kane J, Lorentz S, Firulli AB, Ware SM. SHROOM3 is downstream of the planar cell polarity pathway and loss-of-function results in congenital heart defects. Dev Biol. 2020;464(2):124–136. doi:10.1016/j.ydbio.2020.05.013

169. Chowdhury B, Xiang B, Liu M, Hemming R, Dolinsky VW, Triggs-Raine B. Hyaluronidase 2 Deficiency Causes Increased Mesenchymal Cells, Congenital Heart Defects, and Heart Failure. Circ Cardiovasc Genet. 2017;10(1):e001598. doi:10.1161/CIRCGENETICS.116.001598

170. Wang D, Fang J, Lv J, Pan Z, Yin X, Cheng H, Guo X.Novel polymorphisms in PDLIM3 and PDLIM5 gene encoding Z-line proteins increase risk of idiopathic dilated cardiomyopathy. J Cell Mol Med. 2019;23(10):7054–7062. doi:10.1111/jcmm.14607

171. Li J, Su H, Zhu Y, Cao Y, Ma X. ETS2 and microRNA-155 regulate the pathogenesis of heart failure through targeting and regulating GPR18 expression. Exp Ther Med. 2020;19(6):3469–3478. doi:10.3892/etm.2020.8642

172. Lv L, Li T, Li X, Xu C, Liu Q, Jiang H, Li Y, Liu Y, Yan H, Huang Q, et al. The lncRNA Plscr4 Controls Cardiac Hypertrophy by Regulating miR-214. Mol Ther Nucleic Acids. 2018;10:387–397. doi:10.1016/j.omtn.2017.12.018

173. Bertoli-Avella AM, Gillis E, Morisaki H, Verhagen JMA, de Graaf BM, van de Beek G, Gallo E, Kruithof BPT, et al. Mutations in a TGF-β ligand, TGFB3, cause syndromic aortic aneurysms and dissections. J Am Coll Cardiol. 2015;65(13):1324–1336. doi:10.1016/j.jacc.2015.01.040

174. Grossman TR, Gamliel A, Wessells RJ, Taghli-Lamallem O, Jepsen K, Ocorr K, Korenberg JR, Peterson KL, Rosenfeld MG, Bodmer R, et al. Over-expression of DSCAM and COL6A2 cooperatively generates congenital heart defects. PLoS Genet. 2011;7(11):e1002344. doi:10.1371/journal.pgen.1002344

175. Andenæs K, Lunde IG, Mohammadzadeh N, Dahl CP, Aronsen JM, Strand ME, Palmero S, Sjaastad I, Christensen G, Engebretsen KVT, et al. The extracellular matrix proteoglycan fibromodulin is upregulated in clinical and experimental heart failure and affects cardiac remodeling. PLoS One. 2018;13(7):e0201422. doi:10.1371/journal.pone.0201422

176. Chen HX, Yang ZY, Hou HT, Wang J, Wang XL, Yang Q, Liu L, He GW. Novel mutations of TCTN3/LTBP2 with cellular function changes in congenital heart disease associated with polydactyly. J Cell Mol Med. 2020;24(23):13751–13762. doi:10.1111/jcmm.15950

177. Flamant M, Placier S, Rodenas A, Curat CA, Vogel WF, Chatziantoniou C, Dussaule JC. Discoidin domain receptor 1 null mice are protected against hypertension-induced renal disease. J Am Soc Nephrol. 2006;17(12):3374–3381. doi:10.1681/ASN.2006060677

178. Wan F, Letavernier E, Abid S, Houssaini A, Czibik G, Marcos E, Rideau D, Parpaleix A, Lipskaia L, Amsellem V, et al. Extracellular Calpain/Calpastatin Balance Is Involved in the Progression of Pulmonary Hypertension. Am J Respir Cell Mol Biol. 2016;55(3):337–351. doi:10.1165/rcmb.2015-0257OC

179. Zhang Y, Shen J, He X, Zhang K, Wu S, Xiao B, Zhou X, Phillips RS, Gao P, Jeunemaitre X, Zhu D. A rare variant at the KYNU gene is associated with kynureninase activity and essential hypertension in the Han Chinese population. Circ Cardiovasc Genet. 2011;4(6):687–694. doi:10.1161/CIRCGENETICS.110.959064

180. Vallvé JC, Serra N, Zalba G, Fortuño A, Beloqui O, Ferre R, Ribalta J, Masana L. Two variants in the fibulin2 gene are associated with lower systolic blood pressure and decreased risk of hypertension. PLoS One. 2012;7(8):e43051. doi:10.1371/journal.pone.0043051

181. Heximer S, Husain M. A candidate hypertension gene: will SPON1 hold salt and water?. Circ Res. 2007;100(7):940–942. doi:10.1161/01.RES.0000265134.57140.da

182. Selvarajah V, Mäki-Petäjä KM, Pedro L, Bruggraber SFA, Burling K, Goodhart AK, Brown MJ, McEniery CM, Wilkinson IB. Novel Mechanism for Buffering Dietary Salt in Humans: Effects of Salt Loading on Skin Sodium, Vascular Endothelial Growth Factor C, and Blood Pressure. Hypertension. 2017;70(5):930–937. doi:10.1161/HYPERTENSIONAHA.117.10003

183. Jain M, Mann TD, Stulić M, Rao SP, Kirsch A, Pullirsch D, Strobl X, Rath C, Reissig L, Moreth K, et al. RNA editing of Filamin A pre-mRNA regulates vascular contraction and diastolic blood pressure. EMBO J. 2018;37(19):e94813. doi:10.15252/embj.201694813

184. Sun L, Lin P, Chen Y, Yu H, Ren S, Wang J, Zhao L, Du G. miR-182-3p/Myadm contribute to pulmonary artery hypertension vascular remodeling via a KLF4/p21-dependent mechanism. Theranostics. 2020;10(12):5581–5599. doi:10.7150/thno.44687

185. Satomi-Kobayashi S, Ueyama T, Mueller S, Toh R, Masano T, Sakoda T, Rikitake Y, Miyoshi J, Matsubara H, Oh H, et al. Deficiency of nectin-2 leads to cardiac fibrosis and dysfunction under chronic pressure overload. Hypertension. 2009;54(4):825–831. doi:10.1161/HYPERTENSIONAHA.109.130443

186. Jiang J, Nakayama T, Shimodaira M, Sato N, Aoi N, Sato M, Izumi Y, Kasamaki Y, Ohta M, Soma M, et al. Haplotype of smoothelin gene associated with essential hypertension. Hereditas. 2012;149(5):178–185. doi:10.1111/j.1601-5223.2012.02242.x

187. Waghulde H, Cheng X, Galla S, Mell B, Cai J, Pruett-Miller SM, Vazquez G, Patterson A, Vijay Kumar M, Joe B.Attenuation of Microbiotal Dysbiosis and Hypertension in a CRISPR/Cas9 Gene Ablation Rat Model of GPER1. Hypertension. 2018;72(5):1125–1132. doi:10.1161/HYPERTENSIONAHA.118.11175

188. Dahal BK, Heuchel R, Pullamsetti SS, Wilhelm J, Ghofrani HA, Weissmann N, Seeger W, Grimminger F, Schermuly RT. Hypoxic pulmonary hypertension in mice with constitutively active platelet-derived growth factor receptor-β Pulm Circ. 2011;1(2):259–268. doi:10.4103/2045-8932.83448

189. Thun GA, Imboden M, Berger W, Rochat T, Probst-Hensch NM. The association of a variant in the cell cycle control gene CCND1 and obesity on the development of asthma in the Swiss SAPALDIA study. J Asthma. 2013;50(2):147–154. doi:10.3109/02770903.2012.757776

190. Zhang T, Ji C, Shi R. miR-142-3p promotes pancreatic β cell survival through targeting FOXO1 in gestational diabetes mellitus. Int J Clin Exp Pathol. 2019;12(5):1529–1538.

191. Wang H, She G, Zhou W, Liu K, Miao J, Yu B. Expression profile of circular RNAs in placentas of women with gestational diabetes mellitus. Endocr J. 2019;66(5):431–441. doi:10.1507/endocrj.EJ18-0291

192. Pheiffer C, Dias S, Rheeder P, Adam S. MicroRNA Profiling in HIV-Infected South African Women with Gestational Diabetes Mellitus. Mol Diagn Ther. 2019;23(4):499–505. doi:10.1007/s40291-019-00404-2

193. Zhang C, Wang L, Chen J, Song F, Guo Y. Differential Expression of miR-136 in Gestational Diabetes Mellitus Mediates the High-Glucose-Induced Trophoblast Cell Injury through Targeting E2F1. Int J Genomics. 2020;2020:3645371. doi:10.1155/2020/3645371

194. Catanzaro G, Besharat ZM, Chiacchiarini M, Abballe L, Sabato C, Vacca A, Borgiani P, Dotta F, Tesauro M, Po A, et al. Circulating MicroRNAs in Elderly Type 2 Diabetic Patients. Int J Endocrinol. 2018;2018:6872635. doi:10.1155/2018/6872635

195. Demirsoy İ, Ertural DY, Balci Ş Çınkır Ü, Sezer K, Tamer L, Aras N. Profiles of Circulating MiRNAs Following Metformin Treatment in Patients with Type 2 Diabetes. J Med Biochem. 2018;37(4):499–506. doi:10.2478/jomb-2018-0009

196. Kasimiotis H, Myers MA, Argentaro A, Mertin S, Fida S, Ferraro T, Olsson J, Rowley MJ, Harley VR. Sex-determining region Y-related protein SOX13 is a diabetes autoantigen expressed in pancreatic islets. Diabetes. 2000;49(4):555–561. doi:10.2337/diabetes.49.4.555

197. Wang H, Shi J, Li B, Zhou Q, Kong X, Bei Y. MicroRNA Expression Signature in Human Calcific Aortic Valve Disease. Biomed Res Int. 2017;2017:4820275. doi:10.1155/2017/4820275

198. Xie H, Zhang E, Hong N, Fu Q, Li F, Chen S, Yu Y, Sun K. Identification of TBX2 and TBX3 variants in patients with conotruncal heart defects by target sequencing. Hum Genomics. 2018;12(1):44. doi:10.1186/s40246-018-0176-0

199. Lappas M. Runt-related transcription factor 1 (RUNX1) deficiency attenuates inflammation-induced pro-inflammatory and pro-labour mediators in myometrium. Mol Cell Endocrinol. 2018;473:61–71. doi:10.1016/j.mce.2018.01.003

200. Liu R, Wei C, Ma Q, Wang W. Hippo-YAP1 signaling pathway and severe preeclampsia (sPE) in the Chinese population. Pregnancy Hypertens. 2020;19:1–10. doi:10.1016/j.preghy.2019.11.002

